# Extreme conservation and rare evolutionary deviations in mammalian NOTCH3 inform the interpretation of CADASIL pathogenicity

**DOI:** 10.1101/2025.09.28.678708

**Authors:** Khawar Sohail Siddiqui, Haluk Ertan, Yi Ren, Anne Poljak, Tharusha Jayasena, Perminder S. Sachdev

## Abstract

**Background/Objectives:** NOTCH3 is a highly conserved transmembrane receptor whose pathogenic variants cause CADASIL, a hereditary cerebral small-vessel disease characterized by vascular smooth muscle cell degeneration and extracellular NOTCH3 accumulation. Most established CADASIL variants alter cysteine residues within the extracellular EGF-like repeats, but the influence of residue position, disulfide connectivity, surrounding sequence, and domain context on NOTCH3 structure and pathogenicity remains incompletely understood. We used comparative mammalian analysis to identify rare changes within conserved NOTCH3 regions and evaluate their potential relevance to disease-associated molecular mechanisms.

**Methods:** NOTCH3 protein sequences from 113 mammalian species were analysed using multiple-sequence alignment together with complementary structural modelling, intrinsic-disorder prediction, stability, aggregation-propensity, solvent-accessibility, and molecular-dynamics analyses.

**Results:** NOTCH3 showed exceptional overall conservation, including all 204 extracellular cysteine positions across nearly all species examined. Three rare sequence configurations were identified: seven cysteine substitutions and one cysteine deletion in jaguar EGFr13–15; an NRR deletion in Brandt’s bat predicted to increase S2 cleavage-site accessibility; and an exon 16-associated X1 deletion affecting EGFr20–22, previously reported in a human CADASIL pedigree and also found in selected mammals. Computational analyses predicted distinct structural consequences, including disruption of the canonical disulfide framework and increased intrinsic disorder across the affected jaguar EGFr13–15 region, increased S2 cleavage-site accessibility in Brandt’s bat, and a potential compensatory disulfide rearrangement with increased aggregation propensity in the X1 deletion isoform.

**Conclusions:** These findings provide a comparative molecular-pathology framework for interpreting NOTCH3 variation beyond cysteine number alone. They support a context-dependent model in which residue position, disulfide architecture, surrounding sequence, domain organization, aggregation propensity, and proteolytic regulation jointly influence the predicted structural and functional consequences of NOTCH3 variants. The identified variants provide naturally occurring, experimentally testable models for investigating NOTCH3 folding, aggregation, receptor activation, and CADASIL-associated pathogenicity.

## 1. Introduction

> *“It is notorious that man is constructed on the same general type or model as other mammals. All the bones in his skeleton can be compared with corresponding bones in a monkey, bat, or seal. So it is with his muscles, nerves, blood-vessels and internal viscera. The brain, the most important of all the organs, follows the same law …”* [1]

NOTCH3 is a large transmembrane receptor essential for cell signaling and development in mammals. It comprises five major regions: signal peptide, extracellular domain (ECD), negative regulatory region (NRR), transmembrane (TM) region, and intracellular cytoplasmic domain (ICD). The ECD contains 34 epidermal growth factor-like repeats (EGFrs), which contribute to ligand interactions and receptor structure, with conserved cysteine residues forming disulfide bonds that are important for the integrity of the EGF fold [2].

Cerebral autosomal dominant arteriopathy with subcortical infarcts and leukoencephalopathy (CADASIL) is a hereditary small-vessel disease caused predominantly by mutations in the EGF-like repeats of NOTCH3 [3]. Many CADASIL-associated variants affect conserved cysteine residues and are thought to disrupt the canonical disulfide-bonding pattern of the EGF repeats, potentially contributing to altered protein conformation, aggregation, and accumulation of granular osmiophilic material (GOM) in cerebral small-vessel walls. Aggregation is considered an important feature of CADASIL pathology, although the molecular relationship between individual NOTCH3 variants, structural perturbation, aggregation, and clinical disease remains incompletely understood [4–6]. Importantly, the consequences of NOTCH3 cysteine variants are not uniform and may depend on their position within the EGF repeat region and the surrounding structural context [7].

Recent large-scale efforts such as the Zoonomia Project have highlighted the value of mammalian comparative genomics for understanding gene function and disease evolution [8, 9]. Comparative analysis across diverse mammalian species can reveal both deeply conserved sequence features and rare lineage-specific deviations, providing naturally occurring examples of sequence variation that can generate hypotheses concerning protein structure and function. However, lineage-specific variation does not by itself establish adaptive significance and may also reflect neutral divergence, compensatory changes, or other evolutionary processes [10].

Against this background, we performed a detailed comparative bioinformatic analysis of NOTCH3 across 113 mammalian species, complemented by structural modelling, protein-stability and aggregation-prediction analyses, and molecular dynamics simulations. We focused particularly on the highly conserved cysteine framework of the ECD and on rare deviations affecting the EGF repeats and NRR. This analysis identified three notable deviations in the analysed reference sequences: extensive cysteine substitutions/deletion affecting EGFr13–15 in jaguar, a deletion within the NRR of Brandt’s bat, and an exon 16-associated deletion affecting EGFr20–22 that is also observed in selected mammalian lineages and has previously been reported as a human NOTCH3-X1 isoform.

These rare comparative deviations provide sequence contexts in which substantial alterations to otherwise highly conserved NOTCH3 architecture have been retained. Rather than implying that such variants are necessarily adaptive, benign, or pathogenic, we use them as evolutionary examples to examine how changes in cysteine/disulfide architecture, local structural properties, aggregation propensity, and NRR accessibility may relate to NOTCH3 pathology and biology. The findings therefore refine the interpretation of NOTCH3 variation by emphasizing the importance of residue position, surrounding sequence, domain architecture, and evolutionary context when considering CADASIL-associated variants. The study also generates experimentally testable hypotheses concerning the structural and functional consequences of these naturally occurring deviations.

## 2. Materials and Methods

### Bioinformatics analysis

**A total of 113 mammalian NOTCH3 protein sequences were retrieved from the NCBI database. A multiple-sequence alignment (MSA) was performed using Clustal Omega** available at www.ebi.ac.uk/jdispatcher/msa/clustalo [11]. The mammalian NOTCH3 MSA from Clustal-Omega was exported to Jalview 2.11.3.3 available at www.jalview.org for editing and analysis [12]. The Jalview interactive alignment file is available online (see Supplementary Software1). Per-position consensus values across the 113-sequence NOTCH3 alignment were imported into Origin v2020b (OriginLab Corporation, Northampton, MA, USA). All plots feature interactive tooltips for direct data visualization. The Origin interactive file is available online (see Supplementary Software2). Prediction of intrinsically disordered and binding regions in NOTCH3 was carried out using ANCHOR2 and AIUPRED tools available at https://aiupred.elte.hu/ [13, 14]. The structures of various extracellular and cytoplasmic domains of NOTCH3 were generated using AlphaFold3, available at https://alphafoldserver.com/about [15, 16]. Protein structures were analyzed and visualized using ChimeraX available at https://www.rbvi.ucsf.edu/chimerax [17]. The pathogenic mutations of NOTCH3 were retrieved from the respective publications. NOTCH3 structures, including the native form and those harboring mutations and deletions, were generated for EGFr domains using AF3 in segments of shorter repeats, as well as for the NRR region. Predicted stability and flexibility changes associated with missense substitutions were evaluated using DynaMut2 (https://biosig.lab.uq.edu.au/dynamut/) as described previously [18]. Aggregation propensity of WT, deletion, and missense-mutant EGFr constructs was estimated using Aggrescan4D (https://biocomp.chem.uw.edu.pl/a4d/<u>#</u>) using the dynamic-mode option [19]. Potential disulfide-bond geometries were assessed using ChimeraX and Disulfide by Design 2 [20]. The solvent accessible surface area (SASA) of residues in protein structures was calculated using GETAREA available at https://curie.utmb.edu/getarea.html [21]

### Molecular Dynamics Simulations

The crystal structure of the human NRR domain was obtained from PDB entry 4ZLP, with missing residues supplemented using AlphaFold3 (AF3) predictions. The open conformation of the human ADAM10 metalloprotease was derived from PDB entry 8ESV, from which both the heavy and light Fab chains, as well as tetraspanin-15, were removed [22]. The NRR sequence of Brandt’s bat was obtained from the genome project by [23], and its structure was predicted using AF3. The ADAM10 metalloprotease sequence of Brandt’s bat (XP_005883346.1) was retrieved from the NCBI Database, and its structure was modelled based on the human open-form ADAM10 structure (8ESV).

The docking conformations between the NRR and ADAM10 metalloprotease of both human and Brandt’s bat were predicted using AlphaFold3 multimer modeling [16]. Input files for molecular dynamics simulations were generated using the CHARMM-GUI platform [24]. Protein structures were prepared with the Amber ff14SB force field [25], and the protonation state was set to pH 7.0 using CHARMM-GUI. Water molecules were modeled using the TIP3P model [26]. All systems were solvated in a cubic box of TIP3P water with dimensions of 300 Å per side.

All simulations and trajectory analyses were performed using the GROMACS 2024.3-gpuvolta software package [27]. Each system was first subjected to 1000 steps of steepest-descent energy minimization. Equilibration was conducted in two phases. Initially, constant-volume (NVT) equilibration was carried out at 303.15 K using the V-rescale thermostat [28] for **4 ns**, with a time step of **1 fs**. The temperature was maintained at 303.15 K during NVT equilibration and subsequent simulations. This temperature was selected as a consistent simulation condition for comparative analysis of the predicted protein complexes, with all systems simulated under identical conditions to enable direct comparison of their relative structural and dynamic properties.

This was followed by constant-pressure (NPT) equilibration, with pressure maintained at 1 bar using the Parrinello–Rahman barostat [29] and semi-isotropic pressure coupling. NPT equilibration was conducted in four stages, during which position restraints were applied to both backbone and side chains, beginning with force constants of 400 and 40 kJ/mol/nm², respectively (as used in the initial NVT phase), and gradually reduced by 100 and 10 kJ/mol/nm² at each stage.

The Verlet cutoff scheme [30] was employed, with a cutoff radius of 1.2 nm for short-range van der Waals interactions. Long-range electrostatics were treated using the Particle Mesh Ewald (PME) method [31]. All bonds involving hydrogen atoms were constrained using the LINCS algorithm [32].

After equilibration, three independent 200 ns production runs were performed using a 2 fs time step. The SHAKE algorithm was employed to constrain all bonds involving hydrogen atoms [33]. For quantitative comparison, trajectory-derived parameters were averaged across the three independent 200-ns simulations, and variability was expressed as the standard deviation among the three replicates. For graphical presentation, standard-deviation error bars were displayed at every 50th data point to maintain visual clarity.

Following the simulations, trajectories were corrected for periodic boundary conditions using the gmx trjconv tool with the flags -pbc mol -center. Trajectory analyses included the calculation of backbone root mean square deviation (RMSD) using the gmx rms tool. The solvent-accessible surface area (SASA) was computed using gmx sasa, and the distance between the Zn atom of ADAM10 and the cleavage site of the NRR was measured using gmx dist.

## 3. Results

The comparative analysis identified strong overall NOTCH3 conservation and three rare sequence deviations that were selected for structural and computational evaluation.

### 3.1. Mammalian NOTCH3 shows high sequence conservation and conserved domain architecture

We assessed conservation using MSA of NOTCH3 from 113 mammalian species (Supplementary Figure 1) via Jalview (Supplementary Software 1). Combined with published pathogenicity data and domain annotations, results were visualized in Origin plots (Supplementary Software 2). Among primates, identity with human NOTCH3 ranged from 94.7% to 99.8%, while most other mammals showed ≥90% identity, except armadillo (89.7%). EGFr1–12 showed near-perfect identity, and the ICD was also highly conserved (95–100%) (Figure 1a). The full-length AF3-predicted NOTCH3 structure (Figure 1b,c) is provided as an illustrative structural context to facilitate understanding of the sequence, domain organization, and mutation distribution shown in Figure 1a. Although individual EGF-like domains are generally well defined, the relative positions and orientations of domains in the full-length model should be interpreted with caution because of low inter-domain confidence. The high conservation is consistent with strong evolutionary constraint; however, conservation alone does not establish the functional or clinical consequence of an individual variant.

**Figure 1.**
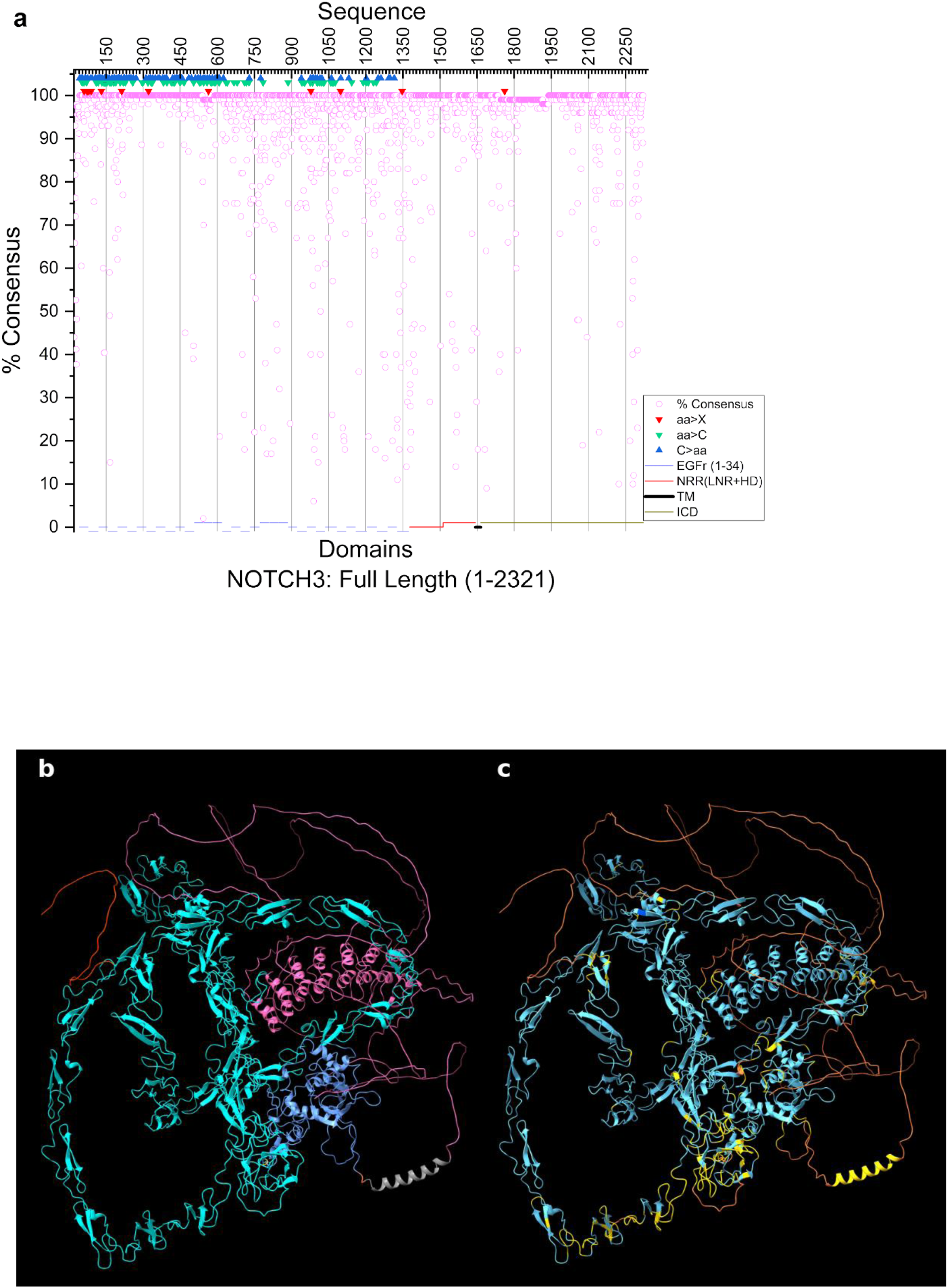
Sequence conservation and structural overview of human NOTCH3**. a**, Origin plot showing the NOTCH3 sequence, domain organization, pathogenic mutations, and percentage consensus across 113 mammalian NOTCH3 sequences. Domains are shown on the X-axis as colored segments: 34 EGF-like repeats (EGFrs) in the extracellular domain (ECD), blue; NRR (LNR + HD), red; transmembrane domain (TM), black; intracellular domain (ICD), green; percentage consensus, pink circles. EGFr10–11 (391–467 aa) are involved in ligand binding. Lifted bars indicate the EGFr regions corresponding to jaguar EGFr13–15 (507–618 aa) and the X1 deletion spanning EGFr20–22 (771–885 aa). Various types of reported CADASIL-associated NOTCH3 mutations are shown as triangles: aa>X, red; aa>C, green; C>aa, blue. The interactive Origin plot features tooltips, where hovering over data points reveals the exact percentage consensus, amino acid sequence, residue name, and mutation information (Supplementary Software 2). Exploded views of EGFr13–15 and EGFr20–22 are shown in Supplementary Figs. 2b and 2c. **b and c**, illustrative structural overview of the AF3-predicted NOTCH3, provided primarily to visualize its domain organization and prediction confidence. The pLDDT profile indicates comparatively high confidence across many of the extracellular EGF-like domains, whereas much of the ICD is predicted with low local confidence; the separate IUPred analysis provides sequence-based evidence of disorder propensity. Accordingly, the full-length model should not be interpreted as a definitive representation of the native global conformation or the relative orientation of individual domains. **b**, color-coded domains—signal peptide (red), EGFr1–34 (cyan), NRR (blue), TM helix (grey), and ICD (pink). **c**, pLDDT confidence scores—blue: >90 (very high); light blue: 70–90 (confident); yellow: 50–70 (low); orange: <50 (very low).

To explore evolutionary insights relevant to CADASIL, we selected four species—human, rhesus macaque, mouse, and naked mole-rat—for focused comparison based on biomedical relevance and unique traits ( **Supplementary Figure 3**). The mouse is widely used in CADASIL models; the rhesus macaque is phylogenetically close to humans; and the naked mole-rat is a long-lived rodent with notable anti-aging traits [35]. Interestingly, naked mole-rat NOTCH3 shares higher sequence identity with human (92.8%) than with mouse (91.0%), a notable exception within the typical 90–99% mammalian identity range. This trend persists in the ECD (EGFr1–34): 92.7% identity with human vs. 89.2% with mouse (Supplementary Software 2). This indicates that naked mole-rat NOTCH3 is modestly more similar in sequence to human NOTCH3 than is mouse NOTCH3. This sequence difference does not, by itself, establish superior structural or functional suitability as a disease model. In the AF3 comparison, the human ECD aligns better with rhesus than with mouse, consistent with sequence identity (Figures 2a, b). The first 11 EGF repeats in human and rhesus superimpose closely, reflecting conserved structure in CADASIL-prone and ligand-binding regions, while divergence beyond EGFr11 suggests differences in inter-repeat orientation, loop flexibility, or species-specific indels in more tolerant mid- and C-terminal regions.

**Figure 2.**
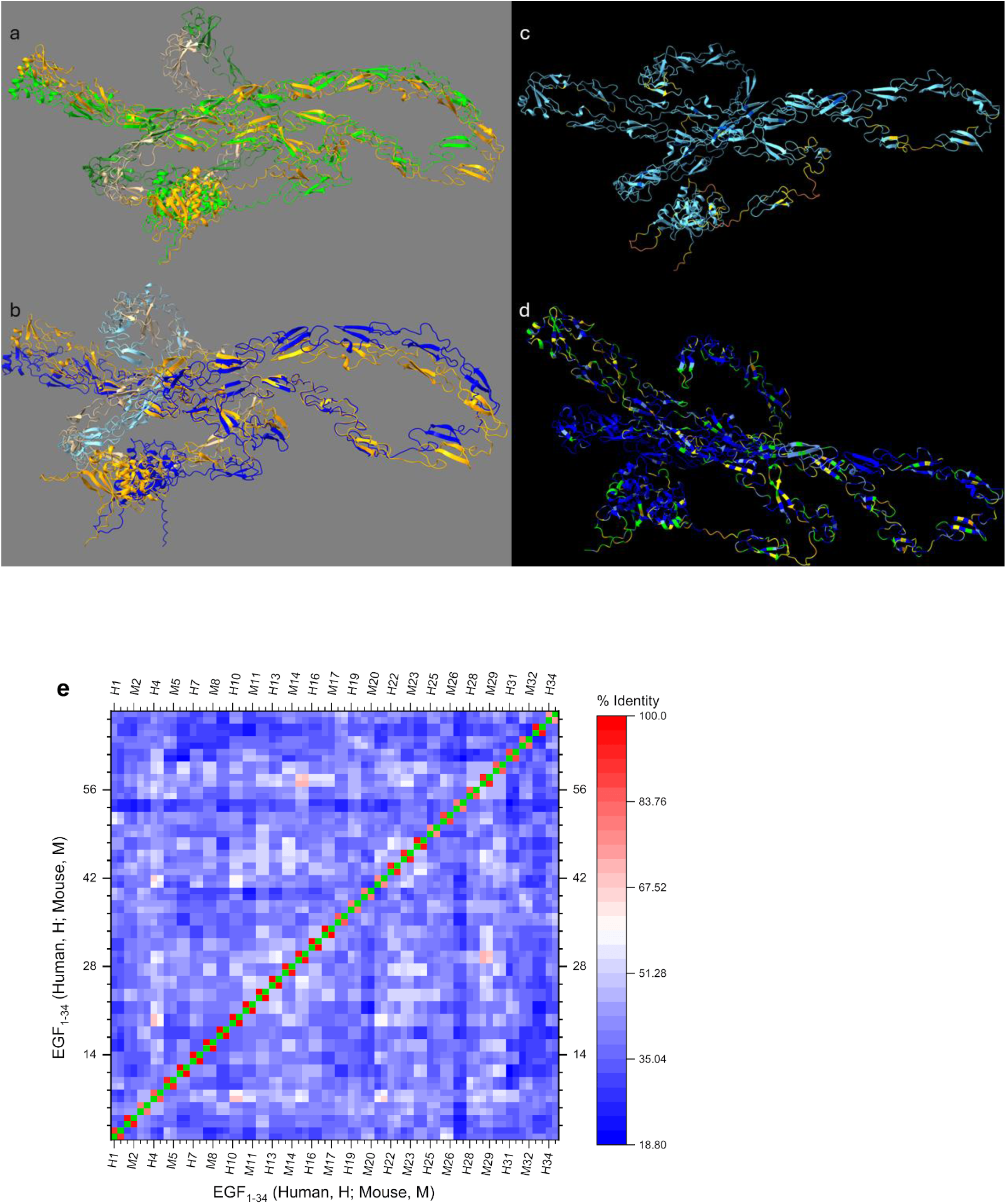
Sequence and structural similarity in ECD. **a**, human and rhesus (RMSD across all 1603 residue pairs, 17.34 Å), **b**, human and mouse (RMSD across all 1603 residue pairs, 23.9 Å). Light orange/dark orange, human; light green/dark green, rhesus, light blue/dark blue, mouse. Darker shades represent EGFr1-11. **c**, human NOTCH3 extracellular domain structure predicted by AlphaFold3. **c**, colored according to pLDDT confidence scores as in Fig. 1. The AlphaFold3-predicted structure of the human NOTCH3 ECD shows high confidence for individual EGFr domains but low confidence in their relative orientation. **d**, human ectodomain showing conservation among >113 mammalian species. Blue, 100%; corn-blue, 98-99%, lime, 90-97%; yellow, 50-89%; <50%, orange. Blue-light blue, high conservation; lime-yellow, medium conservation; orange, low conservation. The relative positioning of EGFr and NRR shows low confidence. **e**, sequence similarity among EGFr1–34 domains in human and mouse NOTCH3. Heatmap showing pairwise sequence identity within human EGFr domains (H vs H), within mouse EGFr domains (M vs M), and between human and mouse EGFr domains (H vs M). The green diagonal represents 100% identity for self-comparisons of the same EGFr domain; the orange-red diagonal indicates sequence identity between corresponding EGFr domains in human and mouse; off-diagonal values above and below the main diagonals show the percentage identity between non-corresponding EGFr domains within and between both species. Structurally, EGFr6 and EGFr31 had an RMSD of 3.815 Å (37 aligned residues), while EGFr4 and EGFr10 had an RMSD of 1.102 Å (38 residues). Other comparisons: EGFr4 vs. EGFr6, 41.7% identity, RMSD 3.81 Å; EGFr6 vs. EGFr10, 35.0%, RMSD 4.2 Å; EGFr4 vs. EGFr31, 50.0%, RMSD 1.0 Å.

The AlphaFold3-predicted structure of the human NOTCH3 ECD (Figure 2c) shows high confidence for individual EGFr domains but low confidence in their relative orientation. A domain-wise conservation map across >113 mammals (Figure 2d) highlights strong conservation in many EGFrs, aligning with sequence similarity and supporting their structural and functional significance. Sequence identity between corresponding EGFr domains in the human and mouse ECD was remarkably high (Figure 2e, diagonal), suggesting evolutionary conservation driven by structural and functional constraints (Figure 2a, b). However, comparing all 34 human and mouse EGFr domains revealed lower pairwise identity (21.6%–68.4%), with high variability across domains (Figure 2e diagonal; Supplementary Figure 4). The mean identity across EGFr1–34 was 41.1%; EGFr6 showed the lowest identity (21.6%) to EGFr31, while EGFr4 had the highest (68.4%) to EGFr10.

To complement sequence analysis, we examined the 3D structures of the extracellular domain (ECD; Fig. 2a, b) and intracellular domain (ICD; Figure 3a), along with IUPRED disorder predictions. AF3 predicted high confidence (pLDDT 70–90) for most of the first 30 EGFr domains, while EGFr20–21 and EGFr30–34 showed lower confidence (Figure 2c), in regions that also showed elevated IUPred-predicted disorder propensity (Figure 3b). Despite high sequence conservation, much of the human NOTCH3 ICD is predicted by AF3 with low confidence, while IUPred independently predicts extensive disorder propensity (Figure 3a,b). Similar disorder was predicted in the ICDs of mouse (Figure 3a), rhesus macaque, and naked mole-rat (Supplementary Figure 3). ANCHOR2 predicts potential disordered binding regions; this is compatible with, but does not demonstrate, partner-induced ordering (Figure 3b, red plot)—a dynamic feature often overlooked in classical structural studies [36]. In contrast, the EGFr1–34 and NRR domains are predicted to be largely ordered (Figure 2c).

**Figure 3.**
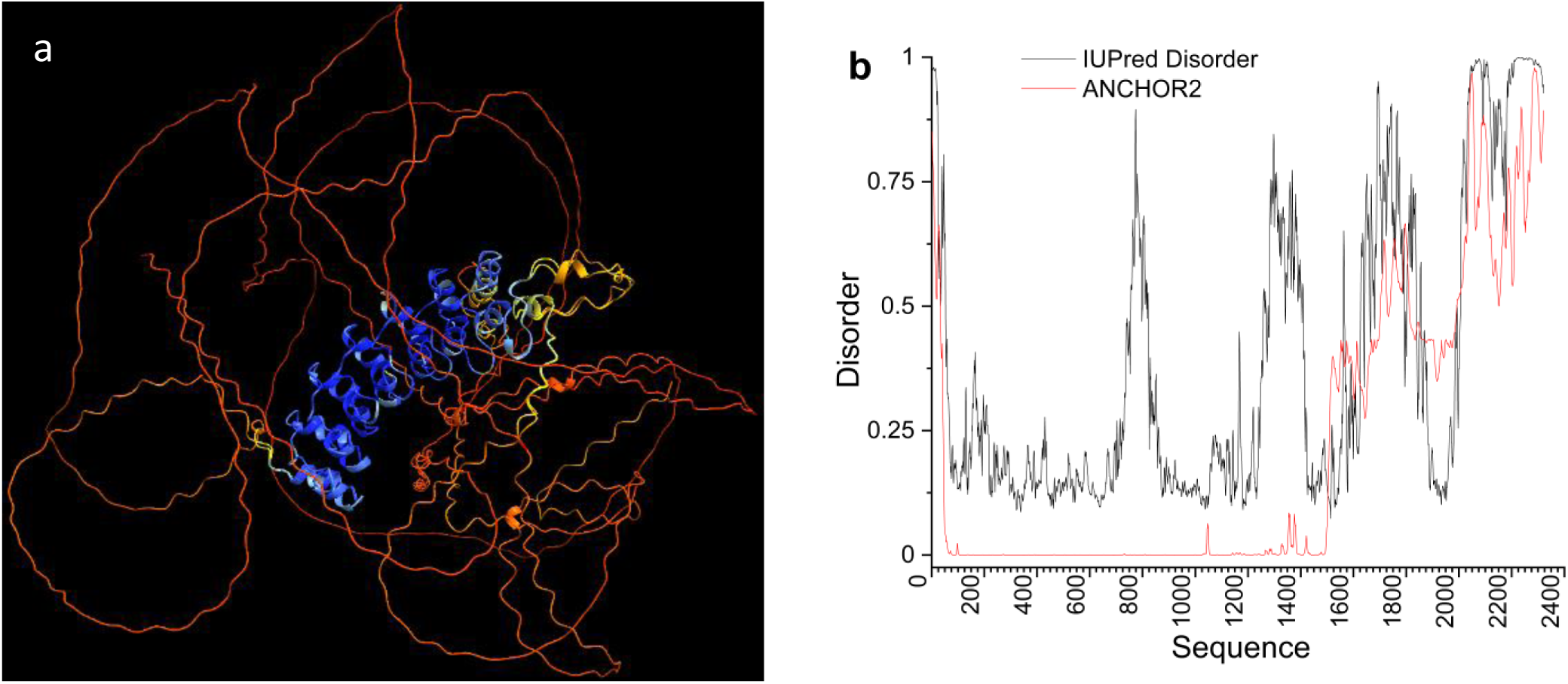
Structure and analysis of intrinsically disordered regions. **a**, superimposed human and mouse NOTCH3 ICD predicted by AF3. Refer to Fig. 1 for pLDDT confidence score. **b**, plot showing intrinsically disordered regions in the NOTCH3 sequence using two different tools: IUPred and ANCHOR2.

### 3.2. Exon 16 Skipping Produces an ECD Deletion Isoform

The MSA revealed key differences in NOTCH3 ECD sequences across mammals, particularly in missense substitutions and deletions. These database-annotated deletion isoforms provide candidate comparative contexts for examining how structural alterations, including those involving conserved cysteine residues and disulfide bonds, may be accommodated within NOTCH3. A deletion corresponding to exon 16 skipping was annotated in several mammalian protein records, including great apes, marmoset, elephants, and vampire bat, but was absent from other primates, bats, and rodents. Two whale isoforms showed a distinct upstream deletion (Figure 4a). A conserved Gly>Asp missense substitution precedes this deletion in all affected species (Figure 4a, arrow).

**Figure 4.**
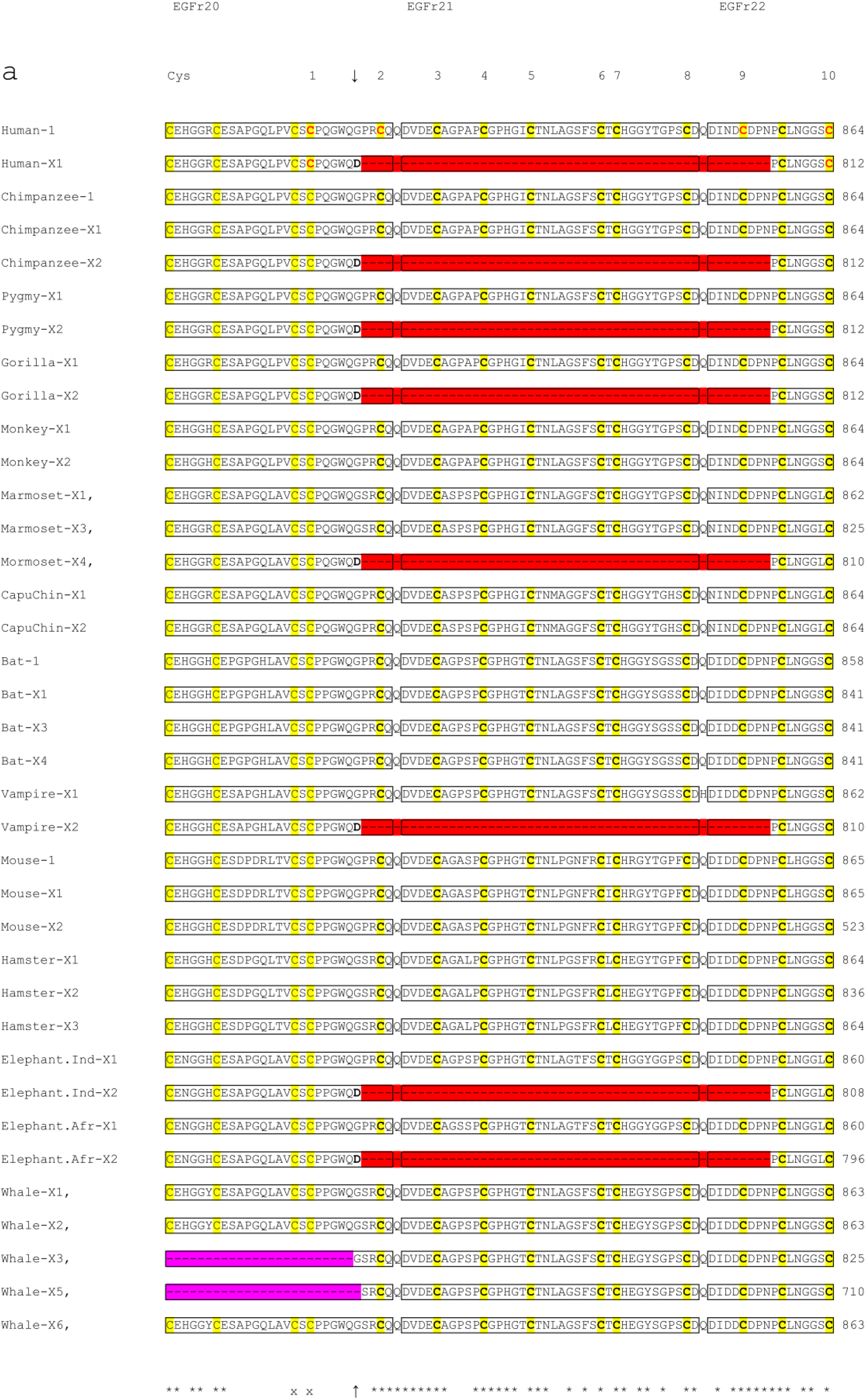
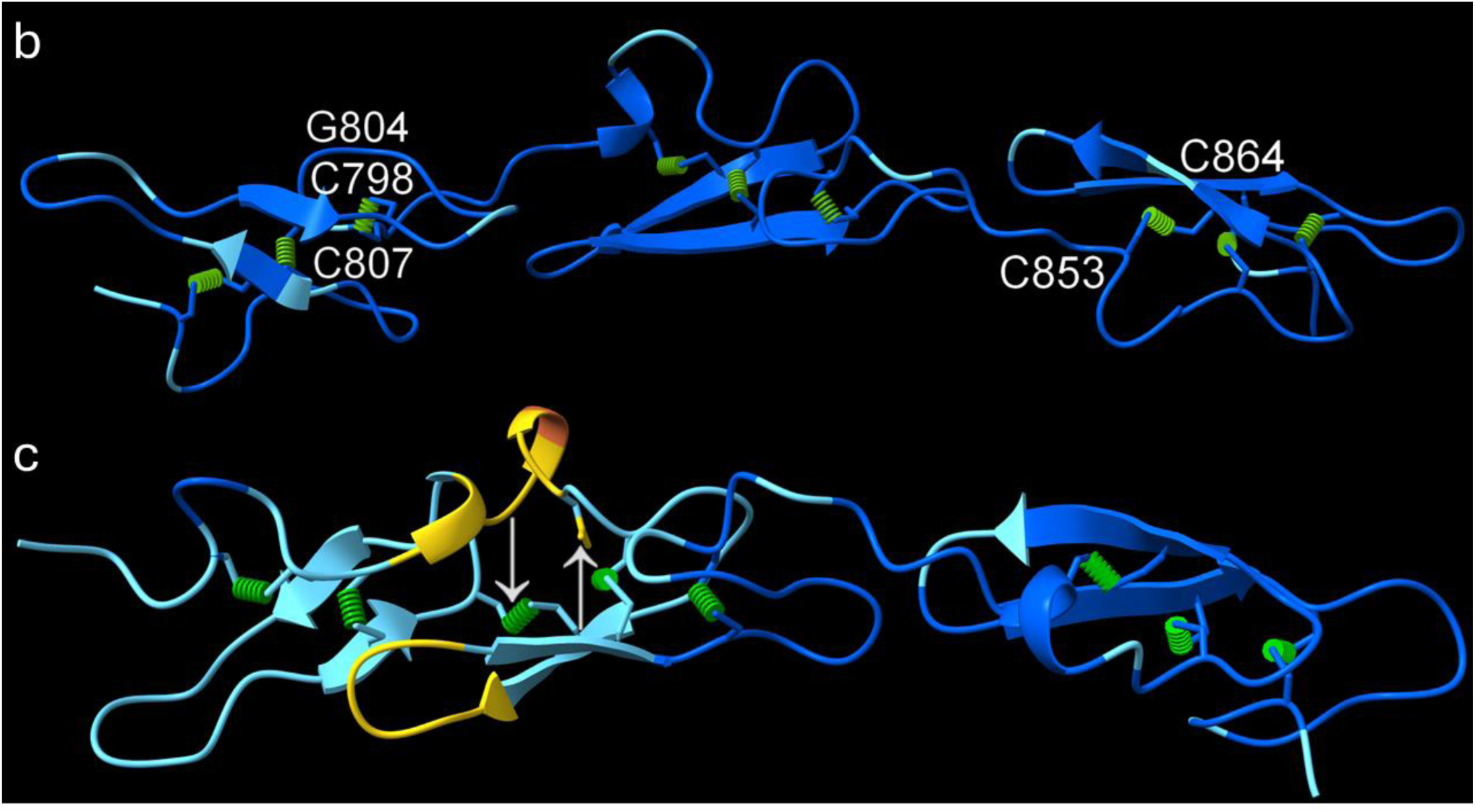
Sequence and structural analysis of the NOTCH3 X1 deletion isoform. a, MSA of the human NOTCH3 X1 isoform compared with NOTCH3 sequences from selected mammals. Chimpanzee, *Pan troglodytes*; pygmy chimpanzee, *Pan paniscus*; gorilla, *Gorilla gorilla gorilla*; monkey, *Rhinopithecus roxellana*; marmoset, *Callithrix jacchus*; capuchin, *Sapajus apella*; bat, *Myotis myotis*; vampire bat, *Desmodus rotundus*; mouse, *Mus musculus*; hamster, *Cricetulus griseus*; Indian elephant, *Elephas maximus indicus*; African elephant, *Loxodonta africana*; and blue whale, *Balaenoptera musculus*. The arrows and bold D indicate the beginning of the deletion. Asterisks indicate conserved residues, whereas x indicates positions where the corresponding cysteine is absent in some sequences. The eight cysteine residues removed by the X1 deletion (human sequence) are shown in bold. The deleted region spans residues P805–N856 and includes eight cysteines. The canonical disulfide bonds C_3_814–C_5_826, C_4_820–C_6_835, and C_7_837–C_8_846 are located within EGFr21, whereas C_1_798–C_2_807 and C_9_853–C_10_864 form disulfide bonds within EGFr20 and EGFr22, respectively, in full-length NOTCH3. In the X1 deletion isoform, Cys_1_798 and Cys_10_812 are predicted to be spatially compatible with the formation of a non-canonical compensatory disulfide bond and are shown in red and bold. b, AF3-predicted structure of the great ape NOTCH3 EGFr20–22 region without the X1 deletion. c, AF3-predicted structure of the great ape NOTCH3 X1 deletion isoform, including EGFr23, colored according to pLDDT confidence: dark blue, >90 (very high); light blue, 70–90 (confident); yellow, 50–70 (low). pLDDT scores are shown as a qualitative measure of prediction confidence. All predicted disulfide bonds are shown as green dashed lines. Cys798 and Cys853 are highlighted because they correspond to cysteine positions affected by the deletion. Cys798 is predicted to form a non-canonical disulfide bond with Cys812 shown by a downward arrow (corresponding to Cys864 in NOTCH3 without the deletion) following the deletion. The G804D substitution is indicated by an upward arrow.

The deletion spans the final four residues of EGFr20, the entire EGFr21, and the first eight residues of EGFr22, eliminating eight cysteine residues: one each in EGFr20 and EGFr22 and all six in EGFr21 (Figure 4a–c). This removes the three canonical disulfide bonds of EGFr21 and one bond each from EGFr20 and EGFr22. Although Cys798 and Cys812 are not paired in the canonical sequence, the deletion substantially alters the spacing of the remaining cysteines and may permit formation of a non-canonical disulfide bond (Figure 4c, inset). ChimeraX and Disulfide by Design 2 [20] analyses indicated that a Cys798–Cys812 disulfide bond is geometrically feasible, with a bond distance of 1.895 Å, χ3 angle of −80.04°, and estimated bond energy of 2.70 kcal/mol, although the geometry suggests some degree of strain. The Sigma B-factor (174.36), which provides an estimate of local atomic mobility in Disulfide by Design 2, indicates relatively restricted local flexibility around the predicted Cys798–Cys812 bond, consistent with a geometry that could support disulfide-bond formation. These observations suggest that, despite the loss of eight cysteine residues, the deletion may not necessarily result in persistent free thiols because local structural rearrangement could permit compensatory disulfide-bond formation. However, experimental studies will be required to establish whether this bond forms in the native protein.

AF3 modelling of the deletion variant predicted local structural changes within the fused EGFr20–22 region (Figure 4c), together with increased aggregation propensity (A4D score, −0.34 versus WT, −0.59). Thus, preservation of a potential compensatory disulfide bond does not necessarily restore the native structural or functional properties of the affected region. The structural and biological implications of this human-associated exon 16-skipping isoform, originally reported by the Hirose group in a CADASIL pedigree [37], therefore warrant further investigation. Importantly, this isoform has been associated with clinical features of CADASIL, including granular osmiophilic material (GOM) deposition and vascular smooth muscle cell degeneration [37], despite the predicted formation of a compensatory Cys798–Cys812 disulfide bond. This observation indicates that preservation of local disulfide-bonding capacity alone is insufficient to predict normal NOTCH3 function or absence of pathogenicity. Interestingly, full-length ECD modelling with AF3 did not predict the Cys798–Cys812 disulfide bond, likely reflecting the low confidence in domain-to-domain positioning in large multidomain models. Collectively, these observations indicate that cysteine conservation or local disulfide-bond formation cannot, by itself, be used as a surrogate for structural integrity or clinical pathogenicity, and that higher-order domain architecture, sequence context and aggregation propensity must also be considered.

### 3.3. Jaguar NOTCH3 Cysteine Variants Disrupt EGFr Integrity

In humans, the three disulfide bonds in each EGFr domain contribute critically to maintaining the structural integrity of the NOTCH3 protein, and mutations affecting cysteine residues are associated with CADASIL [34]. The prevailing hypothesis posits that loss of a disulfide bond or an odd number of cysteines in the ECD promotes high-molecular-weight S–S-linked aggregation [38]. We therefore examined whether naturally occurring cysteine variation in mammalian NOTCH3 is associated with corresponding changes in predicted structural stability and aggregation propensity.

To assess natural variation in the 204 highly conserved NOTCH3 cysteines, we analyzed ECD sequences from 113 mammalian species using Clustal and JALVIEW (Supplementary Software 1). The analysed jaguar reference sequence contained seven cysteine substitutions and one deletion across EGFr13–15, eliminating all six cysteines in EGFr14. Additionally, the last cysteine of EGFr13 (C542) and the first of EGFr15 (C589) were substituted (Figure 5a). This was the only apparent departure from complete cysteine conservation among the analysed reference sequences and represents an unusual set of eight cysteine alterations affecting consecutive EGFr13–15 regions, in sharp contrast to their complete conservation in all other species (Supplementary Software 1). The eight affected cysteines comprise all six cysteines of EGFr14 and one cysteine each from EGFr13 and EGFr15.

**Figure 5.**
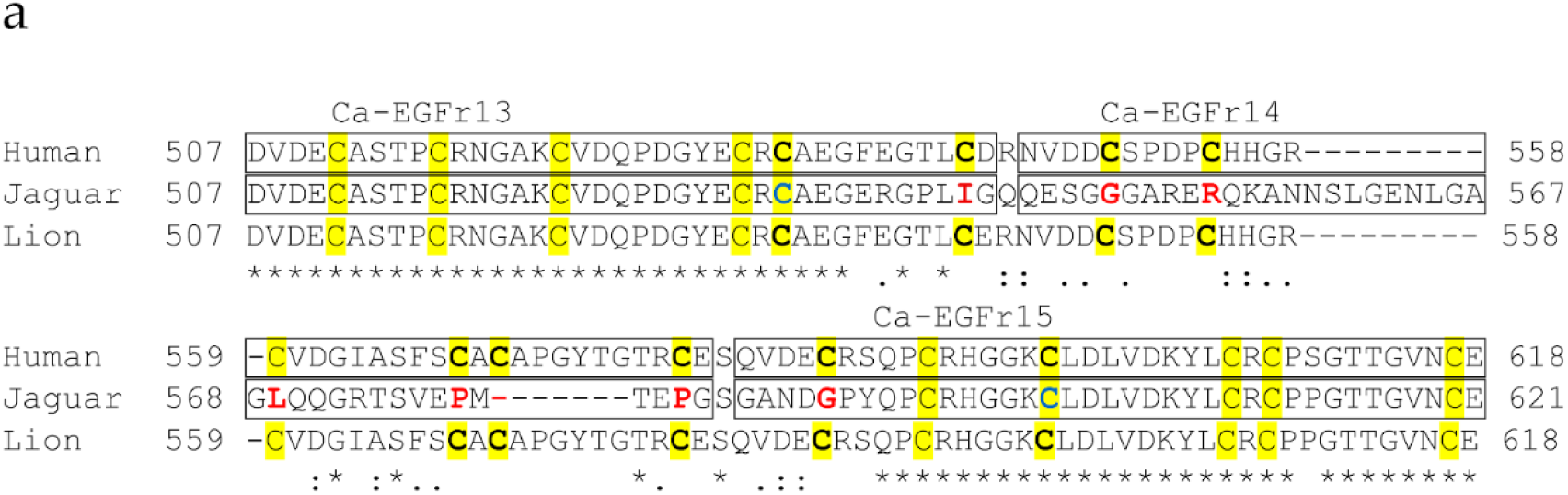
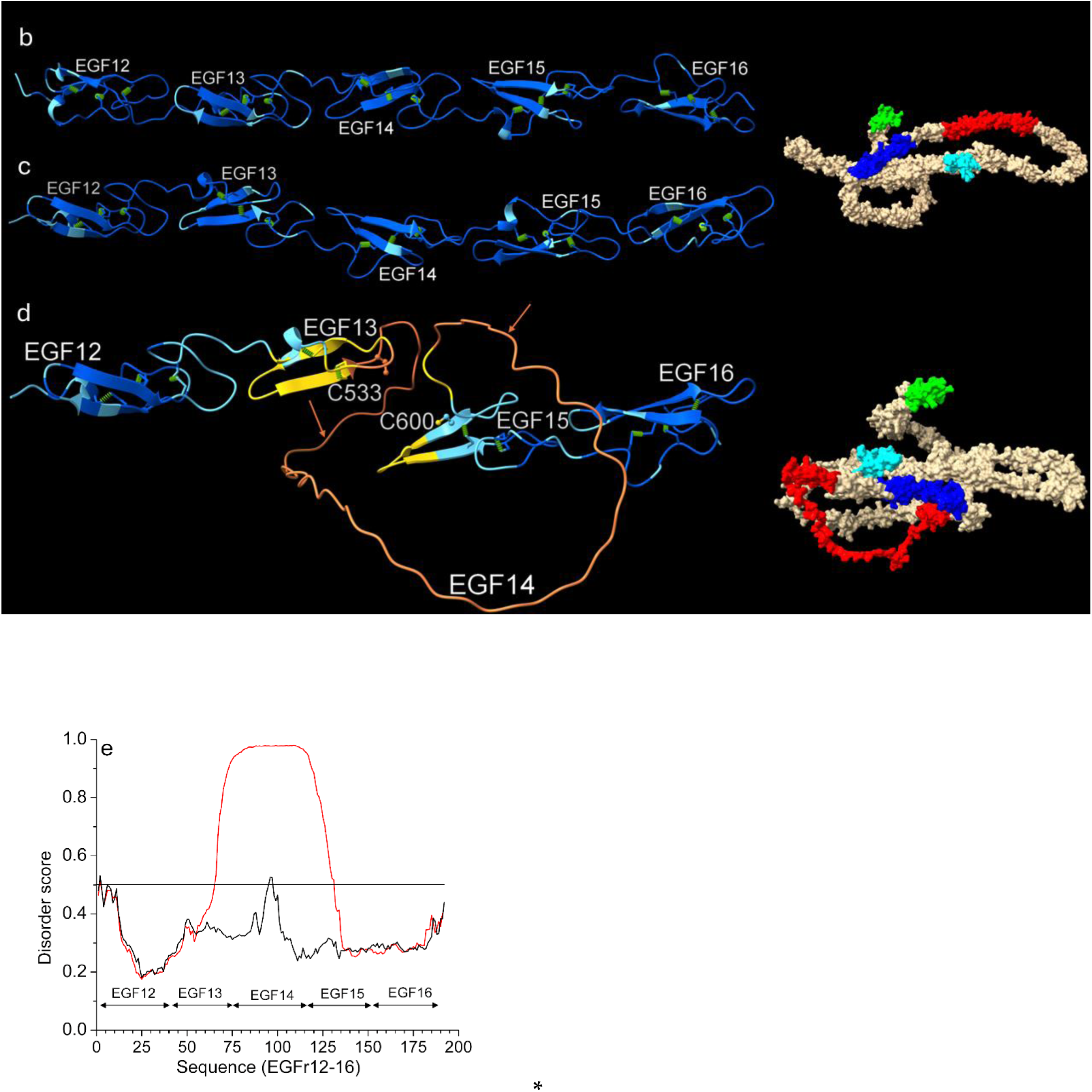
Sequence and structural analysis of cysteine variants in jaguar NOTCH3**. a**, MSA of jaguar EGFr13–15 compared with the corresponding human and lion NOTCH3 sequences. Seven Cys>X substitutions and one deletion in the jaguar sequence are shown in bold red. All six cysteine residues of EGFr14 are absent, resulting in loss of its canonical disulfide-bond network. In EGFr13 and EGFr15, C533 and C600, respectively, lack their canonical cysteine partners because the corresponding partner residues are substituted by I542 and G589, respectively. **b–d**, AF3-predicted structures of NOTCH3 EGFr12–16 from **b**, human; **c**, lion; and **d**, jaguar, colored according to pLDDT confidence as described in Figure 1. pLDDT scores are used as a qualitative measure of prediction confidence. In the jaguar structure, C533 and C600 lack their canonical disulfide-bond partners, while all six cysteines of EGFr14 are absent (between orange arrows). Disulfide bridges predicted within the canonical architecture are shown as green cylinders. EGFr12 and EGFr16, which retain their canonical cysteine/disulfide architecture, provide comparatively well-defined structural regions for reference. Disulfide by Design 2 analysis did not identify the corresponding canonical disulfide bonds involving C533 and C600. However, alternative intramolecular or intermolecular disulfide bonding cannot be excluded experimentally. Inset figures show space-filled AF3 models of the human (upper) and jaguar (lower) ECD, with EGFr13–15 in red, ligand-binding EGFr10–11 in blue, EGFr1 in green, and the remaining EGFrs in cyan and tan. The human EGFr13–15 model is comparatively well defined, whereas the corresponding jaguar region has low prediction confidence; the exact jaguar conformation should not be inferred from this model. e, IUPred disorder-propensity analysis of the jaguar WT sequence (red line) and the corresponding sequence with all eight affected cysteine residues computationally restored (black line), showing markedly reduced disorder propensity following cysteine restoration in the EGFr13–15 region.

Using AF3, we modelled the structural impact of these naturally occurring variants in the jaguar and compared them with structures from humans and lions (Figures 5b–d). In humans and lions, EGFr12–16 maintained compact, folded conformations consistent with intact disulfide-bond networks. In humans and lions, AF3 predicted comparatively compact EGFr12–16 models with intact disulfide-bond networks. In contrast, the jaguar EGFr14 region exhibited very low pLDDT confidence, consistent with increased conformational uncertainty following loss of all three canonical disulfide bonds. EGFr13 and EGFr15 also showed reduced prediction confidence in regions containing unpaired cysteines, whereas EGFr12 and EGFr16 retained comparatively compact, higher-confidence predicted structures, consistent with their preserved cysteine/disulfide architecture (Figure 5d). The reduced pLDDT in the affected region is therefore consistent with loss of the disulfide constraints that normally contribute to the compact EGF fold. Because of the low prediction confidence in the affected region, the precise three-dimensional arrangement of EGFr13–15 in the jaguar should not be inferred from the AF3 model. Rather, the model was used qualitatively to compare prediction confidence between affected and unaffected regions; it was not used to infer a unique conformation or experimentally established compactness.

To independently assess disorder propensity, we analyzed the jaguar EGFr12–16 sequence using IUPred. The jaguar sequence showed elevated predicted disorder propensity within the cysteine-affected EGFr13–15 region, with the highest disorder scores occurring around EGFr14 (Figure 5e). Notably, computational restoration of the eight affected cysteine residues markedly reduced the predicted disorder propensity across the corresponding region, indicating that the jaguar-specific cysteine alterations are associated with increased sequence-based disorder propensity. This sequence-based prediction is concordant with the low AF3 confidence in the same region, although IUPred disorder scores do not constitute experimental evidence of an intrinsically disordered region.

When the WT lion EGFr12–16 region was mutated to carry all eight jaguar Cys>aa substitutions, it showed marked predicted destabilization (ΔΔG = 34.85 kcal/mol), consistent with the reduced structural confidence observed in the corresponding AF3 model. However, A4D scores indicated either no change (WT vs. mutated lion: –0.84) or a slight decrease (WT jaguar: –0.68) in aggregation tendency. Restoring seven Cys residues and retaining the other jaguar-specific sequence changes reduced ΔΔG to 16.34 kcal/mol and the aggregation score to –0.90. Reverting all seven Cys>aa mutations and restoring the deleted cysteine further reduced the predicted destabilization to 4 kcal/mol. DynaMut2 does not determine whether disulfide bonds reform; its outputs were therefore interpreted only as predicted stability and flexibility changes for point-substitution models within the tool’s validated scope. These analyses indicate that restoration of cysteine residues alone did not fully recover the predicted structural properties of the reference sequence, suggesting that the surrounding sequence context also contributes to the structural effects of the jaguar variants.

Importantly, the substantial predicted structural perturbation was not accompanied by increased aggregation propensity. Thus, the jaguar sequence provides a naturally occurring example in which extensive alteration of the canonical cysteine/disulfide architecture is associated with reduced predicted structural compactness but not with increased predicted aggregation tendency. The computational analyses therefore indicate that cysteine status, predicted structural stability, and aggregation propensity are not necessarily coupled in a simple manner.

### 3.4. NRR Deletion in Brandt’s Bat is Predicted to Increase S2 Accessibility

The NRR domain of human NOTCH3 prevents premature receptor activation. It includes three Lin12-Notch repeats (LNRs) and a heterodimerization (HD) domain, which maintain NOTCH3 in an autoinhibited state in the absence of ligand binding (Figure 6a). The NRR shields the S2 cleavage site from ADAM proteases, ensuring NOTCH3 activation primarily occurs upon ligand (e.g., Jagged or Delta-like) engagement. Ligand binding induces mechanical or allosteric disruption of the NRR, exposing the S2 site for ADAM cleavage, followed by γ-secretase cleavage and release of the NOTCH3 ICD [39,40] (Figure 6b). Previous structural studies have shown that the NOTCH3 NRR is intrinsically less stable than the corresponding NRRs of other NOTCH receptors and that this property contributes to its propensity for ligand-independent activation [41]. Constitutive or ligand-independent NOTCH3 signaling has also been reported in pathological contexts [42]. These observations provide an important structural and functional context for evaluating naturally occurring variation within the NOTCH3 NRR.

**Figure 6.**
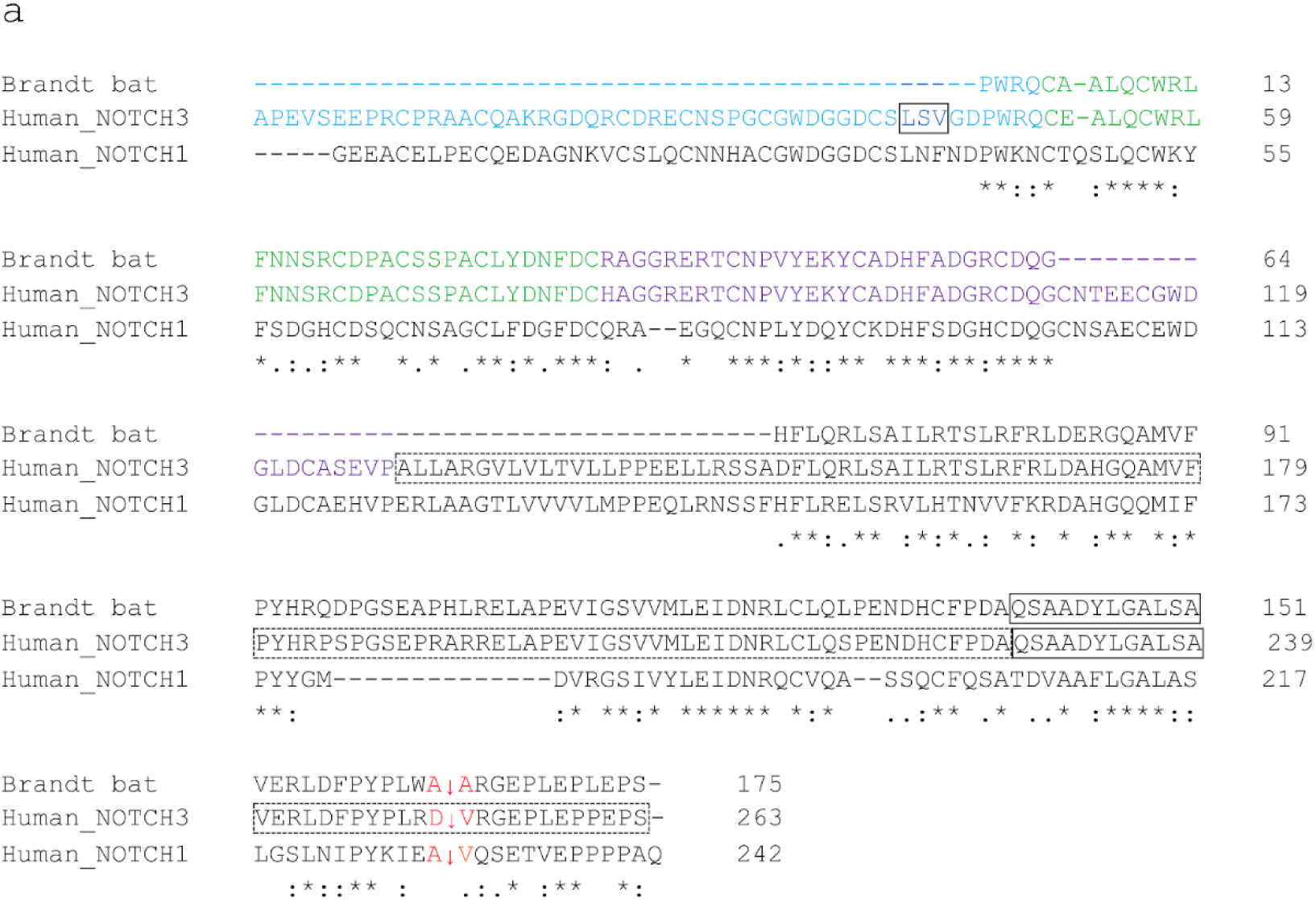
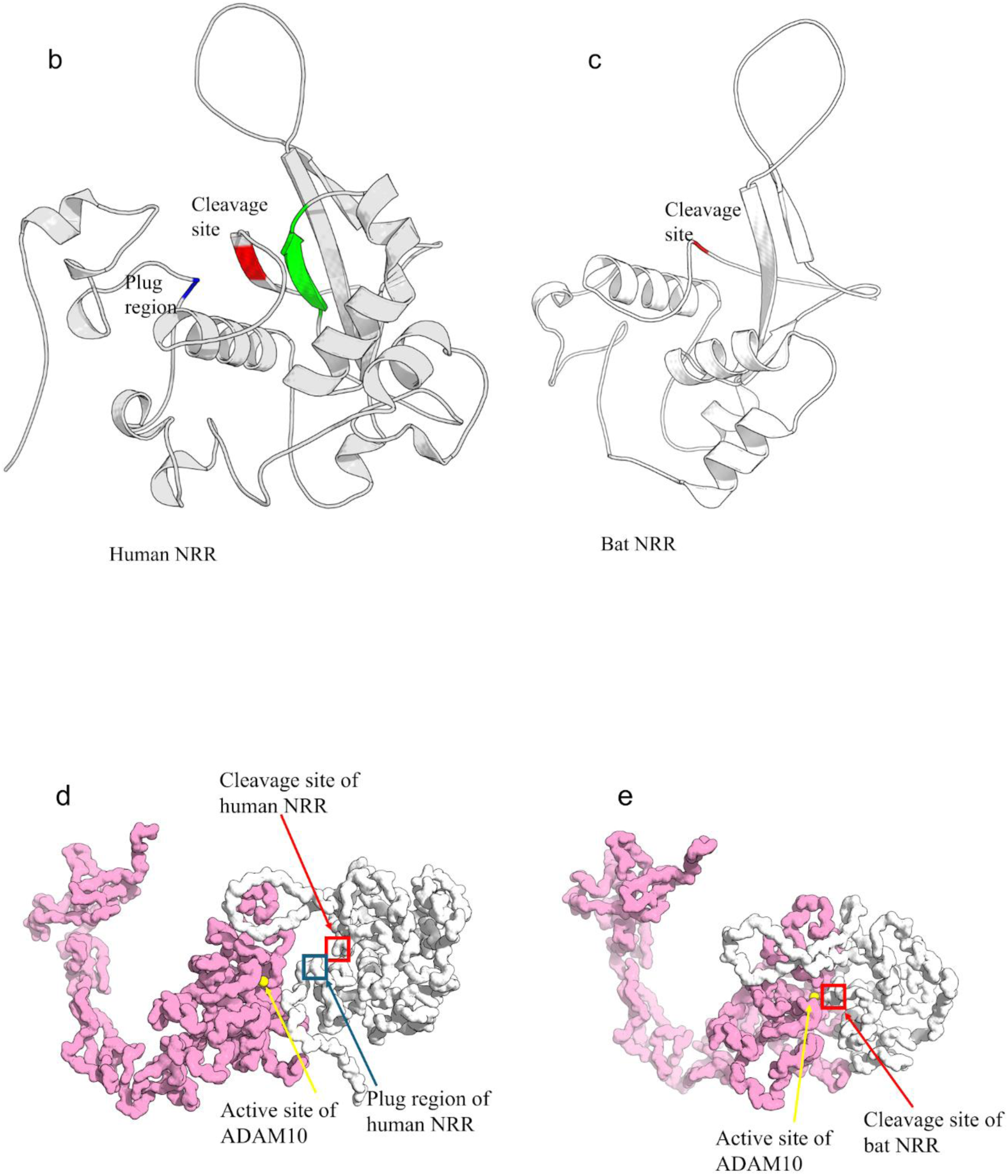
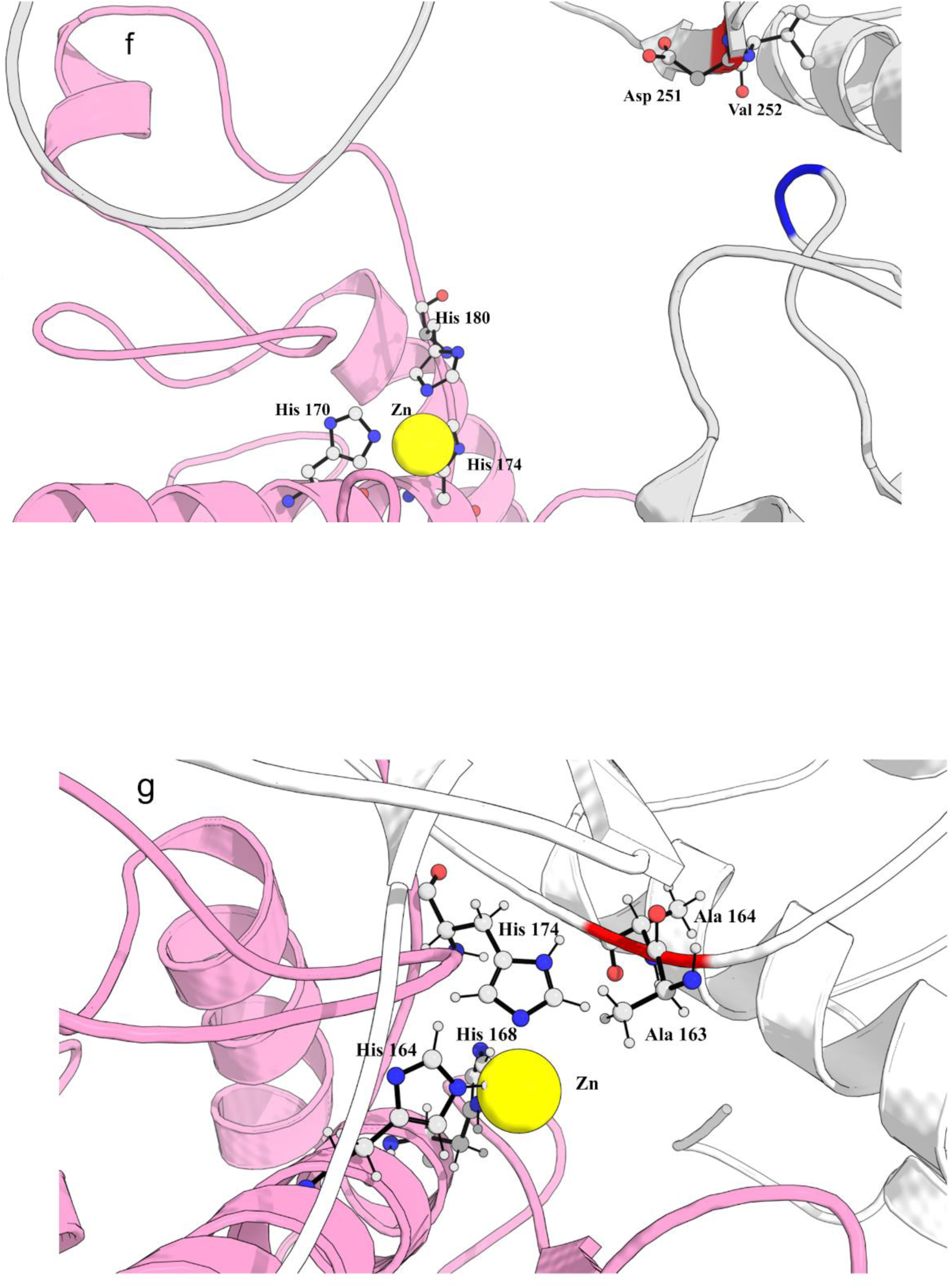
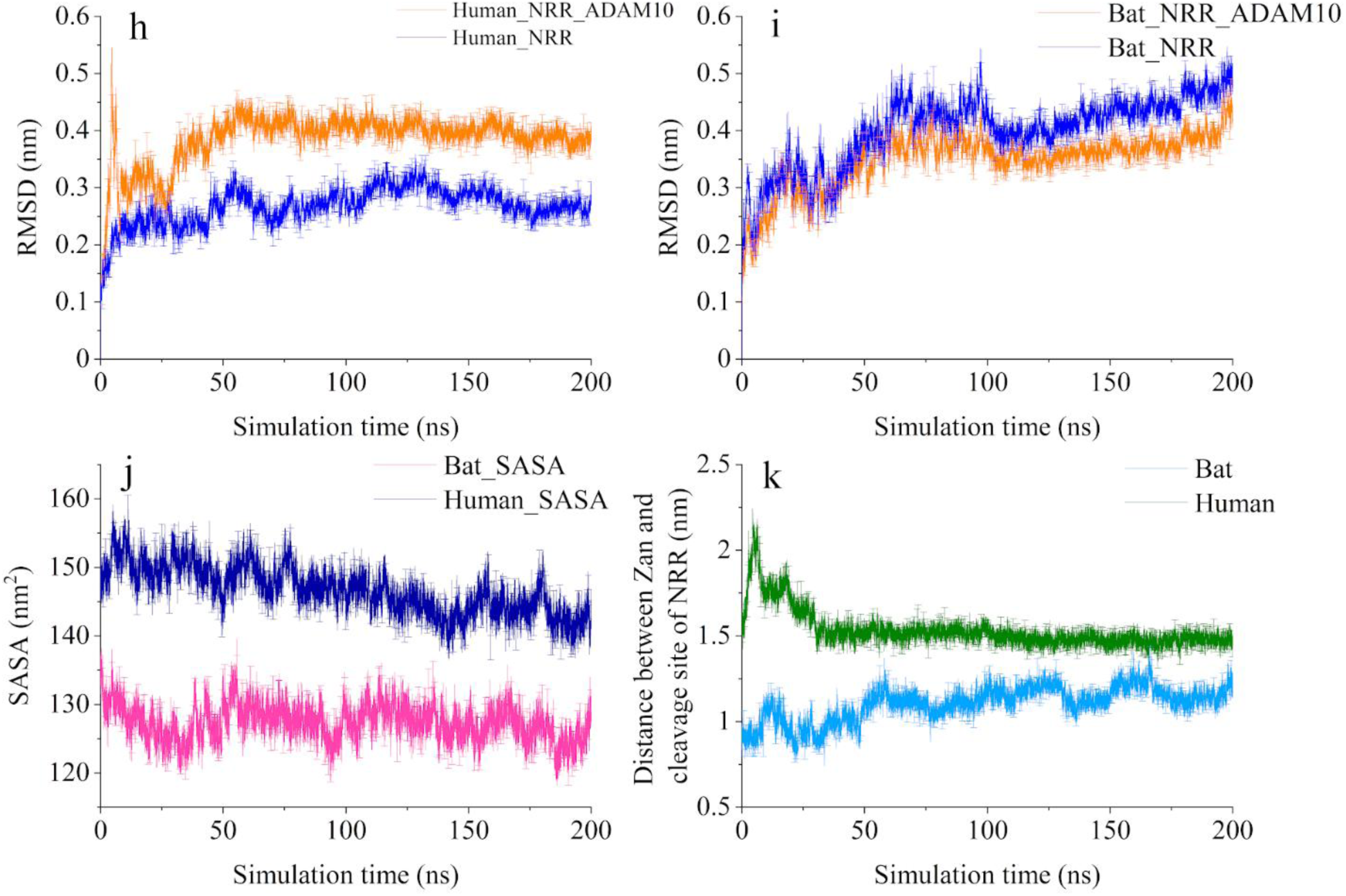
Sequence, structural, and MDS analysis of the NRR-ADAM 10 metalloprotease complexes from human and Brandt’s bat. **a**, shows aligned sequences of human NOTCH1 and NOTCH3 alongside the NRR sequence of Brandt’s bat. Blue, LNR-A showing boxed plug; green, LNR-B; purple, LNR-C; broken box, NRR-HD; solid box, α_3_-helix; and red arrows, S2 cleavage site. **b**, human NRR; showing the S2 cleavage site (red) between a β-strand (green) and plug (blue); **c**, bat NRR structure. The α3-helix is shown below the S2 site. **d-k**, 200 ns simulations for human and bat NRR-ADAM10 complexes. Grey, NRR; pink, ADAM10. **d**, overall view of Human NRR-ADAM10 complex. **e**, overall view of the bat NRR-ADAM10 complex. **f**, close-up of ADAM10-NRR complex in human. **g**, close-up of ADAM10-NRR complex in bat. Red box, S2 cleavage site in NRR. Blue box, plug region of human NRR. The yellow sphere, Zn ion indicates the location of the active site of ADAM10. All ADAM10 were open-forms. **h**, the RMSD of backbone atoms of the human NRR-ADAM10 complex and the human NRR. **i**, the RMSD of backbone atoms of the bat NRR-ADAM10 complex and bat NRR. **j**, the comparison of the solvent accessible surface area (SASA) of bat and human structures. **k**, comparison of distances between Zn ion at the active site centre of ADAM10 metalloprotease and the center of mass of the S2 cleavage site in bat (AA) and human (DV). Solid lines represent the mean of three independent 200-ns MD simulations; error bars indicate standard deviation and are displayed at every 50th data point for clarity.

Multiple sequence alignment (MSA) of mammalian NOTCH3 (Supplementary Software 1) showed that the analysed *Myotis brandtii* NOTCH3 reference sequence lacks two peptide segments within its NRR—one within LNR-A and another within LNR-C (Figure 6a). The apparent LNR-A deletion is particularly notable because it removes a segment positioned over the S2 region and therefore has the potential to alter accessibility of this proteolytic cleavage site. One missing segment in LNR-A (LNF/LSV), which contributes to the structural environment surrounding the S2 site, is absent in *M. brandtii*, potentially increasing exposure of S2 to ADAM proteases (Figures 6b,c). GETAREA analysis further supports this possibility: human S2 residues D152 and V153 show SASA values of 21.9 Å² and 1.8 Å², respectively, whereas the corresponding bat residues A163 and A164 exhibit substantially higher values of 90.3 Å² and 44.1 Å². The markedly greater solvent accessibility of the bat S2 residues is consistent with reduced shielding of this region. All other examined mammals, including vampire bats and *Myotis daubentonii*, *M. myotis*, and *M. kuhlii*, retain the corresponding sequences.

Simulations of the NRR–ADAM10 complex revealed that in the bat model (Figures 6e, g), the S2 site lies closer to the catalytic Zn ion than in the human complex (Figures 6d, f). This difference is consistent with the possibility that the LNR-A deletion alters the local NRR conformation and facilitates access of the protease to S2. Within the sampled 200-ns trajectories, neither complex showed large backbone-RMSD drift; however, these simulations do not establish long-timescale stability of the docked complexes. (Figures 6h, i). The human complex showed a higher overall solvent-accessible surface area (SASA); this indicates greater solvent exposure but does not, by itself, establish weaker binding (Figure 6j). The Zn–S2 distance remained shorter in the Brandt’s bat model than in the human model across the sampled trajectories (initial 8.4 Å; final 11.8 Å; average 11 Å; Figures 6e,g,k) than in the human complex (initial 15.24 Å; final 15 Å; average 15.33 Å; Figures 6d,f,k). However, the distances remained substantially greater than those associated with a catalytically competent configuration. Accordingly, this difference is interpreted only as relative S2 proximity/accessibility and not as evidence of catalytic engagement or ADAM10-mediated cleavage.

Importantly, these computational observations do not establish that the *M. brandtii* receptor undergoes enhanced ADAM10 cleavage or ligand-independent activation *in vivo*. Rather, they identify a naturally occurring evolutionary alteration of the NRR that may modify the balance between receptor autoinhibition and proteolytic activation. Given that human NOTCH3 itself has an unusually labile NRR and a documented propensity for ligand-independent activation [40,41], the *M. brandtii* deletion provides an informative natural experiment for investigating how further modification of NRR architecture can influence this activation threshold. Experimental characterization of the *M. brandtii* NOTCH3 receptor will be required to determine whether the deletion alters ADAM-mediated S2 cleavage, ligand-independent signaling, or other aspects of receptor function. Such studies are beyond the scope of the present comparative analysis but provide a direct experimental avenue arising from this evolutionary observation.

## 4. Discussion

Cross-species protein analyses are essential for uncovering evolutionary conservation and potentially informative natural variation in vascular proteins such as NOTCH3. While aged domestic and wild animals can exhibit cognitive decline and microvascular changes resembling aspects of human small-vessel disease, no naturally occurring animal model has been definitively established for vascular cognitive impairment and dementia (VCID) or CADASIL. In contrast, several experimental rodent models reproduce key features of these conditions, including inflammation, white matter damage, cognitive impairment, vascular smooth muscle cell (VSMC) degeneration, and GOM deposition [43]. Advances in comparative cerebrovascular imaging across larger mammals may further facilitate translational studies of vascular dysfunction [44]. These observations emphasize the potential value of comparative mammalian biology while also highlighting the distinction between naturally occurring variation and experimentally validated disease models.

In this study, we analyzed NOTCH3 sequences from 113 mammalian species to assess conservation, structural variability, and potential relevance to human disease. Recent large-scale efforts such as the Zoonomia Project have demonstrated the power of mammalian comparative genomics for identifying conserved regions and naturally occurring sequence variation [8, 10]. Consistent with this framework, we found exceptionally strong conservation of NOTCH3 across mammals, together with a small number of striking evolutionary deviations. These included eight cysteine alterations affecting consecutive EGFr13–15 regions in jaguar, an EGFr20–22 deletion corresponding to the human X1 isoform in several mammals, and an NRR deletion in Brandt’s bat. Importantly, the occurrence of these variants in naturally evolving mammalian lineages demonstrates that substantial sequence variation can be retained in regions that are otherwise highly conserved, but does not by itself establish that these variants confer an adaptive advantage.

Our analysis focused on three naturally occurring examples with potential relevance to human NOTCH3 biology: cysteine loss or rearrangement within EGF repeats and deletion of part of the NRR. Previous studies have shown that the clinical significance of NOTCH3 cysteine variants can vary according to their position within the EGF repeat region, with variants in EGFr1–6 generally showing stronger associations with CADASIL-related phenotypes [45,46]. Moreover, NOTCH3 variants outside the classical CADASIL-associated regions may still be associated with sporadic stroke, vascular dementia, and MRI features of cerebral small-vessel disease [47], indicating that cysteine variation should not be interpreted as simply pathogenic or benign on the basis of position alone. Together, these observations provide naturally occurring perturbations that can be used to generate testable hypotheses concerning the relationship between sequence, disulfide architecture, structural stability, receptor regulation, and disease-associated variation. The >90% conservation observed across mammalian NOTCH3 underscores the strong evolutionary constraint on the receptor, particularly within its EGF repeats and conserved cysteine framework. At the same time, the rare deviations identified here indicate that evolutionary conservation should not be interpreted as an absolute predictor of pathogenicity. Rather, the structural context of a residue, surrounding sequence, domain architecture, and potentially compensatory changes elsewhere in the protein may influence the consequences of individual variants.

The human X1 isoform, corresponding to exon 16 skipping, provides an informative example. The deletion removes eight cysteine residues from EGFr20–22 and alters the spacing of the remaining cysteines. Our structural analysis suggests that two remaining cysteine residues may form a potential compensatory intramolecular disulfide bond following exon 16 deletion. Nevertheless, the deletion variant showed a higher predicted aggregation propensity than the wild-type protein. This suggests that a potential compensatory disulfide bond alone may be insufficient to restore a wild-type-like predicted structural profile. However, the predicted local structural properties remain altered, indicating that restoration of potential cysteine pairing does not necessarily restore the native architecture of the affected region. Importantly, this isoform has been reported in a CADASIL pedigree in which patients exhibited pathological features including GOM deposition and VSMC degeneration [37]. Thus, preservation of disulfide-bonding capacity alone cannot be considered sufficient evidence of functional normality. Rather, the X1 isoform illustrates that higher-order domain architecture and sequence context may contribute to NOTCH3 pathogenicity in addition to the presence or absence of individual cysteine residues. The recurrence of this exon-skipping configuration across multiple mammalian lineages, despite the otherwise exceptional conservation of the NOTCH3 cysteine framework, provides a striking natural context in which the same extensive loss of eight cysteine residues is retained in different sequence backgrounds. These mammalian X1-like isoforms therefore provide valuable comparative models for investigating how sequence context and compensatory structural rearrangements may influence the consequences of an exon-skipping event that, in humans, has been associated with CADASIL pathology.

The jaguar sequence represents an even more striking departure from the otherwise extreme conservation of NOTCH3 cysteines. Eight cysteine alterations affect EGFr13–15, including loss of all six cysteines in EGFr14 and loss of one cysteine from each of the neighbouring EGFr13 and EGFr15 domains. AF3 modelling indicates reduced structural confidence and compactness in this region, while IUPred3 independently predicts increased disorder propensity that is substantially reduced when the eight affected cysteines are computationally restored. These convergent computational observations support the possibility that disruption of the canonical disulfide network increases conformational flexibility in the affected region. Nevertheless, the precise three-dimensional arrangement of the jaguar EGFr13–15 region cannot be established from AF3, and the predicted disorder should not be regarded as experimentally demonstrated intrinsic disorder.

The jaguar finding should also be interpreted cautiously in evolutionary terms. Its presence in the available jaguar reference sequence makes it a notable comparative observation, but does not establish population-level fixation, phenotypic tolerance, or adaptive significance. Notably, despite the extensive disruption of the canonical cysteine/disulfide architecture predicted for EGFr13–15, the jaguar from which the reference genome was generated was reported as apparently healthy [48], although the possibility of unrecognized or late-onset neurological or vascular abnormalities cannot be excluded. It does not, however, demonstrate that the mutations are functionally advantageous, nor does it imply that the same variants would be benign in humans. The jaguar lineage may possess compensatory sequence changes or other genetic and physiological features that modify their consequences. Alternatively, the observed configuration could represent lineage-specific divergence or genetic drift. Distinguishing among these possibilities will require comparative population-genetic, biochemical, cellular, and physiological studies.

Our computational analyses further indicate that the effects of cysteine restoration are not simply determined by cysteine number. Introducing the jaguar substitutions into the lion sequence produced substantial predicted destabilization, whereas restoration of the affected cysteines progressively reduced the predicted energetic perturbation. At the same time, the tested cysteine restorations did not consistently restore the canonical disulfide architecture, suggesting that the surrounding sequence context may influence the ability of an EGF domain to recover its native structural configuration. The absence of a corresponding increase in predicted aggregation propensity is also notable. These observations suggest that cysteine loss, structural destabilization, conformational flexibility, and aggregation propensity are related but not necessarily interchangeable properties.

The NRR analysis provides a complementary example of naturally occurring variation outside the EGF region. The Brandt’s bat sequence lacks two NRR segments, including a peptide within LNR-A that contributes to shielding the S2 cleavage region. Previous structural studies have established the importance of the NOTCH NRR in maintaining receptor autoinhibition and regulating access to the S2 cleavage site [39, 40]. Structural and functional studies have shown that the NOTCH3 NRR has distinctive stability and activation properties compared with other NOTCH receptors [41,42]. The MD simulations were performed at 303.15 K under identical conditions for comparative analysis; therefore, the observed differences should be interpreted comparatively rather than quantitatively extrapolated to human physiology at 310.15 K. Notably, body temperature in bats, including *Myotis* species, is physiologically variable and can change substantially with activity, ambient temperature, torpor, and hibernation [49]. Our GETAREA analysis indicated substantially greater solvent accessibility of the corresponding S2 residues in Brandt’s bat, while molecular dynamics simulations showed a shorter average Zn–S2 distance than in the human complex.

However, the present MD analysis was limited to RMSD, SASA, and Zn–S2 distance measurements and therefore does not resolve residue-level fluctuations, interfacial interaction networks, or collective conformational dynamics of the NRR–ADAM10 complex. Additional analyses such as RMSF, hydrogen-bond/contact occupancy, PCA, dynamic cross-correlation, free-energy landscape analysis, and energetic characterization would be required to further test the proposed structural mechanism.

These observations are consistent with the hypothesis that the Brandt’s bat deletion may increase accessibility of the S2 region to ADAM10. However, the simulated Zn–S2 distances remained substantially greater than the catalytic distance expected for cleavage, indicating that the simulations do not demonstrate productive proteolysis or constitutive receptor activation. Rather, they suggest that the deletion may alter the conformational environment of the NRR in a manner that could facilitate access to the cleavage site. This interpretation is consistent with previous work showing that structural destabilization of the NRR can promote ligand-independent NOTCH activation [18, 40], while also emphasizing that the specific functional consequence of the Brandt’s bat deletion remains experimentally unresolved.

Importantly, the present results should not be interpreted as demonstrating that Brandt’s bat NOTCH3 is constitutively active. The deletion may alter the energetic or conformational requirements for S2 exposure, but ligand-independent signaling, altered ligand responsiveness, or other regulatory consequences cannot be distinguished using the present computational data. Cellular experiments measuring NOTCH3 cleavage and NICD production would therefore be required to determine the functional consequence of this naturally occurring NRR deletion. A relevant experimental precedent involves the conserved calcium-binding His60 in NOTCH1, which is replaced by arginine in NOTCH3 (Figure 6a). In NOTCH1, substitution of His60 with proline disrupts calcium binding and increases ligand-independent activity approximately 20-fold, demonstrating that relatively small alterations in LNR architecture can markedly affect the threshold for NOTCH activation [40]. This supports the possibility that the *M. brandtii* NRR deletion may similarly modify receptor activation, although its functional consequences remain to be experimentally determined.

Recent studies highlight the importance of tightly regulated NOTCH3 activity in vascular aging and homeostasis. Large-scale human proteomic profiling identified age-related upregulation of NOTCH3 across multiple tissues, with particularly pronounced increases in the aorta, and further showed that NOTCH3 can induce features of vascular endothelial senescence [50]. Conversely, age-related loss of NOTCH3 expression and signaling in brain vessels has been associated with impaired vascular contractility and reduced cerebral blood flow [43], whereas increased NOTCH3 activity can reduce the maximal dilator capacity of cerebral arteries [34]. Together, these apparently contrasting observations emphasize that appropriate regulation of NOTCH3 activity, rather than simply a change in expression level, is important for vascular homeostasis. In this context, the Brandt’s bat NRR deletion is particularly intriguing, as the predicted increase in S2 accessibility raises the possibility that it may modify the threshold for NOTCH3 activation and thereby influence basal or ligand-dependent signaling. Whether such altered regulation contributes to vascular homeostasis or other aspects of bat physiology and longevity [51] remains an interesting and experimentally testable hypothesis.

The coexistence of extensive NOTCH3 conservation with rare naturally occurring deviations has implications for interpreting human disease-associated variants. Our findings do not invalidate the established association between pathogenic NOTCH3 cysteine mutations and CADASIL. Rather, they suggest that the relationship between cysteine status and pathogenicity is more context-dependent than a simple binary model in which every cysteine alteration necessarily produces the same structural or functional outcome. The human X1 deletion, the jaguar cysteine alterations, and the Brandt’s bat NRR deletion represent distinct natural perturbations of NOTCH3 architecture and regulation, each providing a testable example of how sequence context and domain structure may modify the consequences of otherwise substantial alterations.

An important distinction is that presence in a non-human reference sequence does not establish functional tolerance or clinical benignity. If a variant is genuinely retained at the population level, its consequences may be modified by compensatory sequence changes, differences in expression or regulation, species-specific genetic background, or other physiological factors. Conversely, the same sequence alteration introduced into the human NOTCH3 background may have a different consequence. Therefore, the naturally occurring variants identified here should be viewed as evolutionary models for generating hypotheses about structural and functional constraint, rather than as evidence that corresponding human variants are non-pathogenic.

The present study also has several computational limitations. Sequence conservation and comparative analysis identify evolutionary patterns but do not establish molecular function. AF3 provides valuable structural hypotheses, but low-confidence regions, particularly those affected by extensive cysteine loss or deletions, cannot be interpreted as experimentally resolved structures. Similarly, DynaMut2 and Aggrescan4D provide computational estimates of stability and aggregation propensity rather than direct measurements of these properties. The molecular dynamics simulations provide information about conformational behaviour under the simulated conditions, but the 200-ns trajectories and the measured Zn–S2 distances are insufficient to establish productive ADAM10 cleavage or receptor activation. These limitations have therefore been considered when interpreting the structural and functional implications of the identified variants.

These findings may also have implications for the clinical interpretation of NOTCH3 variants. Although cysteine-altering variants within the EGFr domains remain the molecular hallmark of CADASIL, the clinical manifestations and disease severity associated with NOTCH3 variants vary according to their position within the EGF repeat region, with variants in EGFr1–6 generally associated with more severe disease than those in EGFr7–34. Cysteine-sparing and other atypical NOTCH3 variants present additional challenges for clinical classification, particularly when genetic findings are discordant with phenotype, neuroimaging, segregation, or pathological evidence. Importantly, the naturally occurring mammalian variants identified here should not be interpreted as evidence that analogous human variants are benign or non-pathogenic. Rather, they provide comparative evidence that can inform interpretation by highlighting the potential importance of residue position, local sequence environment, domain architecture, and supporting clinical and pathological evidence when assessing the significance of human NOTCH3 variants.

### 4.1. Future directions

The evolutionary deviations identified in this study provide several experimentally testable avenues for future investigation. Recombinant NOTCH3 EGF constructs carrying the jaguar cysteine alterations could be examined for folding, disulfide connectivity, thermal stability, aggregation, and ligand-binding properties. Cellular expression studies could further determine whether the unusual EGFr13–15 configuration affects receptor maturation, trafficking, cleavage, ligand responsiveness, or downstream signaling. The possibility that the conformationally flexible jaguar EGFr13–15 region could acquire an interaction or regulatory function represents an intriguing and experimentally testable hypothesis, particularly because intrinsically disordered and conformationally flexible protein regions can mediate dynamic protein–protein interactions [52]. NOTCH3 also has an established role in skeletal-muscle regeneration: *NOTCH3*-deficient mice exhibit altered satellite-cell dynamics and develop muscle hyperplasia following repetitive muscle injury [53]. Whether the jaguar EGFr13–15 configuration alters NOTCH3 signaling in skeletal muscle or other tissues remains unknown, but provides an additional direction for functional investigation.

The X1 deletion provides a complementary model for determining how extensive cysteine loss and potential compensatory disulfide rearrangement influence NOTCH3 biology. Expression of the X1 isoform in an appropriate NOTCH3 cellular background could determine whether the predicted structural and aggregation changes affect receptor maturation, trafficking, ligand responsiveness, downstream signaling, or extracellular NOTCH3 accumulation.

For the Brandt’s bat NRR deletion, biochemical and cellular assays of ADAM10-dependent S2 cleavage, NICD generation, and downstream NOTCH3 signaling would directly test whether the predicted increase in S2 accessibility translates into altered receptor activation. Comparison of ligand-dependent and ligand-independent signaling would be particularly informative in determining whether the deletion modifies the activation threshold of the receptor.

Comparative population-genetic and phylogenetic analyses would further help distinguish adaptive evolution from lineage-specific divergence or neutral variation. Emerging artificial intelligence and machine-learning approaches may complement comparative genomics and structural modelling in future studies of disease-associated protein variation [54]. Finally, appropriate cellular, animal, and physiological models will be required to determine whether the naturally occurring NOTCH3 variants identified here have measurable functional or organism-level consequences. Together, these approaches provide a path toward understanding how naturally occurring variation in NOTCH3 relates to structural constraint, receptor function, and human vascular disease.

## 5. Conclusions

Mammalian NOTCH3 sequence–structure–function relationships provide an evolutionary framework for investigating naturally occurring variation relevant to human vascular disease, including CADASIL. The three unusual NOTCH3 variants identified in this study are particularly salient against the background of extreme conservation observed across 113 mammalian species.

Our findings refine and expand the context in which NOTCH3 variants can be interpreted by showing that substantial alterations in cysteine content, disulfide architecture, and NRR sequence occur within selected mammalian sequence contexts. The complementary computational analyses further indicate that the predicted structural consequences of these alterations are influenced by residue position, surrounding sequence, higher-order domain architecture, aggregation propensity, and receptor regulatory context. These naturally occurring deviations therefore provide informative comparative models for investigating NOTCH3 structure–function relationships, while their evolutionary retention should not by itself be interpreted as evidence of clinical benignity or adaptive benefit.

Overall, this study demonstrates the value of comparative mammalian analysis for identifying rare natural perturbations of highly conserved disease-associated proteins and translating them into experimentally testable molecular hypotheses. The variants identified here provide specific models for investigating disulfide connectivity and structural flexibility, extracellular NOTCH3 aggregation, receptor processing and trafficking, S2-site accessibility, and NOTCH3 signaling. Such studies may ultimately improve our understanding of how structural and sequence context influence NOTCH3 function and refine the interpretation of human CADASIL-associated variation.

## Supplementary Materials

The following supporting information can be downloaded at https://www.mdpi.com/article/doi/s1, Figure S1: The multiple sequence alignment of human NOTCH3 with 113 mammals using CLUSTAL-Omega.; S2: The Origin plots showing the NOTCH3 sequence with various domains, pathogenic mutations and % consensus with mammalian protein.; S3: MSA and AF3 structures of NOTCH3.; S4: Plot showing the mean percentage identity and standard deviation for each individual EGFr domain.; Table S1: The stability and flexibility scores of jaguar reverse aa>Cys mutations analysed by DynaMut tool.

## Author Contributions

Conceptualization, K.S.S., H.E., P.S. and A.P.; methodology, K.S.S., Y.R., H.E. and A.P.; software, K.S.S., H.E., Y.R; validation, K.S.S., H.E., A.P., T.J. and Y.R.; formal analysis, K.S.S., A.P., T.J. and H.E.; investigation, K.S.S., H.E., P.S., T.J. and A.P; resources, P.S.; data curation, K.S.S. and H.E.; writing—original draft preparation, K.S.S., Y.R. and H.E.; writing—review and editing, A.P., T.J. and P.S.; visualization, K.S.S., H.E. and Y.R; supervision, P.S.; project administration, P.S.; funding acquisition, P.S. All authors have read and agreed to the published version of the manuscript.

## Data Availability Statement

The additional data that supports the findings of this study are available in the form of interactive data files and raw data is available on Figshare as follows:

**Interactive supplementary data files are publicly available through the Figshare records listed below.** https://figshare.com/account/mycontent/items

Multiple Sequence Alignment (MSA) using **Clustal Omega** and presented using Jalview Software. Identifier: 10.6084/m9.figshare.30223741, Interactive Origin plots show the NOTCH3 sequence with various domains, pathogenic mutations and % consensus with 113 mammalian NOTCH3 sequences. Identifier: 10.6084/m9.figshare.30223747

## Acknowledgments

The authors used **ChatGPT (OpenAI)** to assist with English language editing, improvement of scientific writing, and formatting of references. All generated text was critically reviewed, revised, and verified by the authors, who take full responsibility for the final content of the manuscript.

## Abbreviations

NOTCH3: Neurogenic locus notch homolog protein 3
CADASIL: Cerebral autosomal dominant arteriopathy with subcortical infarcts and leukoencephalopathy
EGFr: epidermal growth factor repeats
ECD: extracellular domain
NRR: negative regulatory region
TM: transmembrane
ICD: intracellular cytoplasmic domain
GOM: granular osmiophilic material
VSMC: vascular smooth muscle cell
ADAM: a disintegrin and metalloproteinase
MDS: molecular dynamics simulation
AF3: AlphaFold3
pLDDT: predicted local distance difference test
SASA: solvent accessible surface area
RMSD: root mean square deviation
LNR: Lin12-Notch repeats
HD: heterodimerization domain
VCID: vascular cognitive impairment and dementia

## References

1. Darwin, C. The Descent of Man, and Selection in Relation to Sex, 2nd ed.; John Murray: London, UK, 1874; p. 6.

2. Hosseini-Alghaderi, S.; Baron, M. Notch3 in development, health and disease. Biomolecules 2020, 10, 485. 10.3390/biom10030485.

3. Boston, G.; Jobson, D.; Mizuno, T.; Ihara, M.; Kalaria, R.N. Most common NOTCH3 mutations causing CADASIL or CADASIL-like cerebral small vessel disease: A systematic review. Cereb. Circ. Cogn. Behav. 2024, 6, 100227. 10.1016/j.cccb.2024.100227.

4. Young, K.Z.; Rojas Ramírez, C.; Keep, S.G.; Gatti, J.R.; Lee, S.J.; Zhang, X.; Ivanova, M.I.; Ruotolo, B.T.; Wang, M.M.;, et al. Oligomerization, trans-reduction, and instability of mutant NOTCH3 in inherited vascular dementia. Commun. Biol. 2022, 5, 1331. 10.1038/s42003-022-03259-2.

5. Lee, S.J.; Zhang, X.; Wu, E.; Sukpraphrute, R.; Sukpraphrute, C.; Ye, A.; Wang, M.M. Structural changes in NOTCH3 induced by CADASIL mutations: Role of cysteine and non-cysteine alterations. J. Biol. Chem. 2023, 299, 104838. 10.1016/j.jbc.2023.104838.

6. Dupré, N.; Gueniot, F.; Domenga-Denier, V.; Dubosclard, V.; Nilles, C.; Hill-Eubanks, D.; Morgenthaler-Roth, C.; Nelson, M.T.; Keime, C.; Danglot, L.;, et al. Protein aggregates containing wild-type and mutant NOTCH3 are major drivers of arterial pathology in CADASIL. J. Clin. Invest. 2024, 134, e175789. 10.1172/JCI175789.

7. Mizuta, I.;, et al. Progress to clarify how NOTCH3 mutations lead to CADASIL, a hereditary cerebral small vessel disease. Biomolecules 2024, 14, 127. 10.3390/biom14010127.

8. Christmas, M.J.;, et al. Evolutionary constraint and innovation across hundreds of placental mammals. Science 2023, 380, eabn3943. 10.1126/science.abn3943.

9. Romero, I.J. Seeing humans through an evolutionary lens: A collection of mammalian genomes provides insights into human biology and evolution. Science 2023, 380, 360–361. 10.1126/science.adh0745.

10. Sullivan, P.F.;, et al. Zoonomia: Leveraging base-pair mammalian constraint to understand genetic variation and human disease. Science 2023, 380, 367. 10.1126/science.abn2937.

11. Madeira, F.; Madhusoodanan, N.; Lee, J.; Eusebi, A.; Niewielska, A.; Tivey, A.R.N.; Lopez, R.; Butcher, S. The EMBL-EBI Job Dispatcher sequence analysis tools framework in 2024. Nucleic Acids Res. 2024, 52, W521–W525. 10.1093/nar/gkae241.

12. Waterhouse, A.M.; Procter, J.B.; Martin, D.M.A.; Clamp, M.; Barton, G.J. Jalview Version 2—a multiple sequence alignment editor and analysis workbench. Bioinformatics 2009, 25, 1189–1191. 10.1093/bioinformatics/btp033.

13. Erdős, G.; Dosztányi, Z. AIUPred: Combining energy estimation with deep learning for the enhanced prediction of protein disorder. Nucleic Acids Res. 2024, 52, W176–W181. 10.1093/nar/gkae385.

14. Mészáros, B.; Erdős, G.; Dosztányi, Z. IUPred2A: Context-dependent prediction of protein disorder as a function of redox state and protein binding. Nucleic Acids Res. 2018, 46, W329–W337. 10.1093/nar/gky384.

15. Jumper, J.; Evans, R.; Pritzel, A.; Green, T.; Figurnov, M.; Ronneberger, O.; Tunyasuvunakool, K.; Bates, R.; Žídek, A.; Potapenko, A.;, et al. Highly accurate protein structure prediction with AlphaFold. Nature 2021, 596, 583–589. 10.1038/s41586-021-03819-2.

16. Abramson, J.;, et al. Accurate structure prediction of biomolecular interactions with AlphaFold 3. Nature 2024, 630, 493–500. 10.1038/s41586-024-07487-w.

17. Meng, E.C.; Goddard, T.D.; Pettersen, E.F.; Couch, G.S.; Pearson, Z.J.; Morris, J.H.; Ferrin, T.E. UCSF ChimeraX: Tools for structure building and analysis. Protein Sci. 2023, 32, e4792. 10.1002/pro.4792.

18. Rodrigues, C.H.M.; Pires, D.E.V.; Ascher, D.B. DynaMut2: Assessing changes in stability and flexibility upon single and multiple point missense mutations. Protein Sci. 2021, 30, 60–69. 10.1002/pro.3942.

19. Zalewski, M.; Iglesias, V.; Bárcenas, O.; Ventura, S.; Kmiecik, S. Aggrescan4D: A comprehensive tool for pH-dependent analysis and engineering of protein aggregation propensity. Protein Sci. 2024, 33, e5180. 10.1002/pro.5180.

20. Craig, D.B.; Dombkowski, A.A. Disulfide by Design 2.0: A web-based tool for disulfide engineering in proteins. BMC Bioinformatics 2013, 14, 346. 10.1186/1471-2105-14-346.

21. Fraczkiewicz, R.; Braun, W. Exact and efficient analytical calculation of the accessible surface areas and their gradients for macromolecules. J. Comput. Chem. 1998, 19, 319–333. 10.1002/(SICI)1096-987X(199802)19:3<319::AID-JCC6>3.0.CO;2-W.

22. Lipper, C.H.; Egan, E.D.; Gabriel, K.H.; Blacklow, S.C. Structural basis for membrane-proximal proteolysis of substrates by ADAM10. Cell 2023, 186, 3632–3641.e10. 10.1016/j.cell.2023.06.026.

23. Seim, I.;, et al. Genome analysis reveals insights into physiology and longevity of the Brandt’s bat Myotis brandtii. Nat. Commun. 2013, 4, 2212. 10.1038/ncomms3212.

24. Jo, S.; Kim, T.; Iyer, V.G.; Im, W. CHARMM-GUI: A web-based graphical user interface for CHARMM. J. Comput. Chem. 2008, 29, 1859–1865. 10.1002/jcc.20945.

25. Maier, J.A.; Martinez, C.; Kasavajhala, K.; Wickstrom, L.; Hauser, K.E.; Simmerling, C. ff14SB: Improving the accuracy of protein side chain and backbone parameters from ff99SB. J. Chem. Theory Comput. 2015, 11, 3696–3713. 10.1021/acs.jctc.5b00255.

26. Jorgensen, W.L.; Chandrasekhar, J.; Madura, J.D.; Impey, R.; Klein, M. Refined TIP3P model for water. J. Chem. Phys. 1983, 79, 926–935.

27. Abraham, M.J.;, et al. GROMACS: High performance molecular simulations through multi-level parallelism from laptops to supercomputers. SoftwareX 2015, 1–2, 19–25. 10.1016/j.softx.2015.06.001.

28. Bussi, G.; Donadio, D.; Parrinello, M. Canonical sampling through velocity rescaling. J. Chem. Phys. 2007, 126, 014101. 10.1063/1.2408420.

29. Ke, Q.; Gong, X.; Liao, S.; Duan, C.; Li, L. Effects of thermostats/barostats on physical properties of liquids by molecular dynamics simulations. J. Mol. Liq. 2022, 365, 120116. 10.1016/j.molliq.2022.120116.

30. Grubmüller, H.; Heller, H.; Windemuth, A.; Schulten, K. Generalized Verlet algorithm for efficient molecular dynamics simulations with long-range interactions. Mol. Simul. 1991, 6, 121–142. 10.1080/08927029108022142.

31. Petersen, H.G. Accuracy and efficiency of the particle mesh Ewald method. J. Chem. Phys. 1995, 103, 3668–3679.

32. Hess, B.; Bekker, H.; Berendsen, H.J.; Fraaije, J.G. LINCS: A linear constraint solver for molecular simulations. J. Comput. Chem. 1997, 18, 1463–1472.

33. Ryckaert, J.-P.; Ciccotti, G.; Berendsen, H.J.C. Numerical integration of the cartesian equations of motion of a system with constraints: Molecular dynamics of n-alkanes. J. Comput. Phys. 1977, 23, 327–341. 10.1016/0021-9991(77)90098-5.

34. Baron-Menguy, C.; Domenga-Denier, V.; Ghezali, L.; Faraci, F.M.; Joutel, A. Increased Notch3 activity mediates pathological changes in structure of cerebral arteries. Hypertension 2017, 69, 60–70. 10.1161/HYPERTENSIONAHA.116.08015.

35. Oka, K.; Yamakawa, M.; Kawamura, Y.; Kutsukake, N.; Miura, K. The naked mole-rat as a model for healthy aging. Annu. Rev. Anim. Biosci. 2023, 11, 207–226. 10.1146/annurev-animal-050322-074744.

36. Morris, O.M.; Torpey, J.H.; Isaacson, R.L. Intrinsically disordered proteins: Modes of binding with emphasis on disordered domains. Open Biol. 2021, 11, 210222. 10.1098/rsob.210222.

37. Saiki, S.; Sakai, K.; Saiki, M.; Kitagawa, Y.; Umemori, T.; Murata, K.; Matsui, M.; Hirose, G. Varicose veins associated with CADASIL result from a novel mutation in the Notch3 gene. Neurology 2006, 67, 337–339. 10.1212/01.wnl.0000224758.52970.19.

38. Chabriat, H.; Joutel, A.; Dichgans, M.; Tournier-Lasserve, E.; Bousser, M.-G. CADASIL. Lancet Neurol. 2009, 8, 643–653. 10.1016/S1474-4422(09)70127-9.

39. Gordon, W.R.; Arnett, K.L.; Blacklow, S.C. The molecular logic of Notch signaling—a structural and biochemical perspective. J. Cell Sci. 2008, 121, 3109–3119. 10.1242/jcs.035683.

40. Gordon, W.R.; Roy, M.; Vardar-Ulu, D.; Garfinkel, M.; Mansour, M.R.; Aster, J.C.; Blacklow, S.C. Structure of the Notch1-negative regulatory region: Implications for normal activation and pathogenic signaling in T-ALL. Blood 2009, 113, 4381–4390. 10.1182/blood-2008-08-174748.

41. Xu, X.; Choi, S.H.; Hu, T.; Tiyanont, K.; Habets, R.; Groot, A.J.; Vooijs, M.; Aster, J.C.; Chopra, R.; Fryer, C.;, et al. Insights into autoregulation of Notch3 from structural and functional studies of its negative regulatory region. Structure 2015, 23, 1227–1235. 10.1016/j.str.2015.05.001.

42. Choy, L.; Hagenbeek, T.J.; Solon, M.; French, D.M.; Finkle, D.; Shelton, A.; Venook, R.; Brauer, M.J.; Siebel, C.W. Constitutive NOTCH3 signaling promotes the growth of basal breast cancers. Cancer Res. 2017, 77, 1439–1452. 10.1158/0008-5472.CAN-16-1022.

43. Romay, M.C.; Knutsen, R.H.; Ma, F.; Mompeón, A.; Hernandez, G.E.; Salvador, J.; Mirkov, S.; Batra, A.; Sullivan, D.P.; Procissi, D.;, et al. Age-related loss of Notch3 underlies brain vascular contractility deficiencies, glymphatic dysfunction, and neurodegeneration in mice. J. Clin. Invest. 2024, 134, e166134. 10.1172/JCI166134.

44. Hainsworth, A.H.; Allan, S.M.; Boltze, J.; Cunningham, C.; Farris, C.; Head, E.; Ihara, M.; Isaacs, J.D.; Kalaria, R.N.; Lesnik Oberstein, S.A.M.J.;, et al. Translational models for vascular cognitive impairment: A review including larger species. BMC Med. 2017, 15, 16. 10.1186/s12916-017-0793-9.

45. Rutten, J.W.; Dauwerse, H.G.; Gravesteijn, G.; van Belzen, M.J.; van der Grond, J.; Polke, J.M.; Bernal-Quiros, M.; Lesnik Oberstein, S.A. Archetypal NOTCH3 mutations frequent in public exome: Implications for CADASIL. Ann. Clin. Transl. Neurol. 2016, 3, 844–853. 10.1002/acn3.344.

46. Hack, R.J.; Gravesteijn, G.; Cerfontaine, M.N.; Santcroos, M.A.; Gatti, L.; Kopczak, A.; Bersano, A.; Duering, M.; Rutten, J.W.; Lesnik Oberstein, S.A.J. Three-tiered EGFr domain risk stratification for individualized NOTCH3-small vessel disease prediction. Brain 2023, 146, 2913–2927. 10.1093/brain/awac486.

47. Cho, B.P.H.; Nannoni, S.; Harshfield, E.L.; Tozer, D.; Gräf, S.; Bell, S.; Markus, H.S. NOTCH3 variants are more common than expected in the general population and associated with stroke and vascular dementia: An analysis of 200 000 participants. J. Neurol. Neurosurg. Psychiatry 2021, 92, 694–701. 10.1136/jnnp-2020-325838.

48. Figueiró, H.V.; Li, G.; Trindade, F.J.; Assis, J.; Pais, F.; Fernandes, G.; Santos, S.H.D.; Hughes, G.M.; Komissarov, A.; Antunes, A.;, et al. Genome-wide signatures of complex introgression and adaptive evolution in the big cats. Sci. Adv. 2017, 3, e1700299. 10.1126/sciadv.1700299.

49. Hock, R.J. The metabolic rates and body temperatures of bats. Biol. Bull. 1951, 101, 289–299. 10.2307/1538547.

50. Ding, Y.; Zuo, Y.; Zhang, B.; Fan, Y.; Xu, G.; Cheng, Z.; Ma, S.; Fang, S.; Tian, A.; Gao, D.;, et al. Comprehensive human proteome profiles across a 50-year lifespan reveal aging trajectories and signatures. Cell 2025, 188, 5763–5784.e26. 10.1016/j.cell.2025.06.047.

51. Gorbunova, V.; Seluanov, A.; Kennedy, B.K. The world goes bats: Living longer and tolerating viruses. Cell Metab. 2020, 32, 31–43. 10.1016/j.cmet.2020.06.013.

52. Ginell, G.M.; Emenecker, R.J.; Lotthammer, J.M.; Keeley, A.T.; Plassmeyer, S.P.; Razo, N.; Usher, E.T.; Pelham, J.F.; Holehouse, A.S. Sequence-based prediction of intermolecular interactions driven by disordered regions. Science 2025, 388, eadq8381. 10.1126/science.adq8381.

53. Kitamoto, T.; Hanaoka, K. Notch3 null mutation in mice causes muscle hyperplasia by repetitive muscle regeneration. Stem Cells 2010, 28, 2205–2216. 10.1002/stem.547.

54. Lu, Y.; Chen, J.; Fan, N.; Song, W.; Sheng, H.; Yang, Y.; Wang, J. Machine learning models for drug-drug interaction prediction from computational discovery to clinical application. npj Digit. Med. 2026, 9, 198. 10.1038/s41746-026-02400-3.

