## Supplemental Material for "Extreme conservation and rare evolutionary deviations in mammalian NOTCH3 inform the interpretation of CADASIL pathogenicity"

| | | | | | | | | |
|---|---|---|---|---|---|---|---|---|
| **1: Human** | | | 100.00 | 99.83 | 99.70 | 99.78 | 99.14 | 99.40 |
| 99.10 | 99.44 | 98.88 | 99.10 | 99.31 | 98.78 | 99.22 | 97.37 | 97.85 |
| 99.27 | 98.07 | 98.88 | 95.03 | 97.53 | 97.58 | 98.66 | 94.73 | 95.43 |
| 91.92 | 90.72 | 91.88 | 93.10 | 91.58 | 90.64 | 90.98 | 90.85 | 91.19 |
| 90.72 | 90.76 | 92.26 | 91.63 | 92.20 | 94.10 | 93.99 | 91.75 | 91.79 |
| 90.86 | 91.95 | 92.25 | 91.62 | 92.60 | 91.50 | 95.48 | 91.63 | 95.43 |
| 91.33 | 93.71 | 94.73 | 91.41 | 95.39 | 91.37 | 91.37 | 91.46 | 93.84 |
| 93.71 | 92.33 | 92.33 | 93.04 | 93.05 | 93.34 | 93.26 | 92.87 | 92.53 |
| 92.61 | 93.34 | 92.44 | 92.40 | 93.09 | 93.08 | 93.16 | 94.40 | 90.93 |
| 93.16 | 93.97 | 93.84 | 94.14 | 93.71 | 90.96 | 93.01 | 92.84 | 92.79 |
| 93.06 | 92.62 | 93.11 | 93.15 | 92.75 | 93.12 | 94.14 | 94.32 | 94.41 |
| 93.93 | 93.84 | 93.94 | 93.94 | 93.89 | 93.75 | 93.66 | 93.84 | 93.75 |
| 93.92 | 93.88 | 93.75 | 93.79 | 92.25 | 93.84 | 89.69 | 91.92 | 90.76 |
| **2: Chimp** | | | 99.83 | 100.00 | 99.78 | 99.78 | 99.14 | 99.40 |
| 99.10 | 99.44 | 98.88 | 99.10 | 99.31 | 98.78 | 99.22 | 97.37 | 97.85 |
| 99.27 | 98.07 | 98.88 | 94.99 | 97.53 | 97.58 | 98.66 | 94.73 | 95.43 |
| 91.84 | 90.64 | 91.79 | 93.10 | 91.50 | 90.55 | 90.90 | 90.77 | 91.11 |
| 90.64 | 90.81 | 92.31 | 91.63 | 92.20 | 94.15 | 94.03 | 91.70 | 91.70 |
| 90.90 | 91.95 | 92.25 | 91.67 | 92.60 | 91.50 | 95.48 | 91.71 | 95.43 |
| 91.33 | 93.75 | 94.78 | 91.46 | 95.39 | 91.41 | 91.37 | 91.46 | 93.84 |
| 93.71 | 92.33 | 92.33 | 93.04 | 93.05 | 93.34 | 93.26 | 92.87 | 92.53 |
| 92.61 | 93.34 | 92.44 | 92.40 | 93.09 | 93.08 | 93.25 | 94.44 | 90.93 |
| 93.16 | 94.10 | 93.97 | 94.27 | 93.84 | 90.96 | 93.01 | 92.84 | 92.79 |
| 93.06 | 92.62 | 93.11 | 93.15 | 92.75 | 93.12 | 94.18 | 94.36 | 94.45 |
| 93.97 | 93.89 | 93.99 | 93.99 | 93.93 | 93.79 | 93.71 | 93.88 | 93.79 |
| 93.97 | 93.92 | 93.79 | 93.84 | 92.25 | 93.88 | 89.74 | 91.92 | 90.72 |
| **3: Pygmy-Chimp** | | | 99.70 | 99.78 | 100.00 | 99.66 | 99.01 | 99.27 |
| 98.97 | 99.31 | 98.75 | 98.97 | 99.18 | 98.65 | 99.09 | 97.24 | 97.72 |
| 99.14 | 97.93 | 98.75 | 94.91 | 97.40 | 97.45 | 98.53 | 94.65 | 95.34 |
| 91.75 | 90.51 | 91.71 | 93.01 | 91.41 | 90.42 | 90.77 | 90.64 | 91.02 |
| 90.55 | 90.72 | 92.18 | 91.63 | 92.11 | 94.06 | 93.94 | 91.57 | 91.61 |
| 90.81 | 91.86 | 92.16 | 91.58 | 92.51 | 91.42 | 95.43 | 91.54 | 95.39 |
| 91.29 | 93.66 | 94.65 | 91.37 | 95.30 | 91.33 | 91.28 | 91.46 | 93.76 |
| 93.62 | 92.24 | 92.24 | 92.96 | 92.96 | 93.26 | 93.17 | 92.79 | 92.44 |
| 92.53 | 93.26 | 92.35 | 92.31 | 93.00 | 92.99 | 93.12 | 94.36 | 90.85 |
| 93.07 | 93.92 | 93.79 | 94.10 | 93.66 | 90.87 | 92.93 | 92.75 | 92.71 |
| 92.97 | 92.53 | 93.02 | 93.06 | 92.66 | 93.03 | 94.10 | 94.28 | 94.36 |
| 93.89 | 93.80 | 93.90 | 93.90 | 93.84 | 93.71 | 93.62 | 93.79 | 93.71 |
| 93.88 | 93.84 | 93.71 | 93.75 | 92.21 | 93.79 | 89.69 | 91.83 | 90.67 |
| **4: Gorilla** | | | 99.78 | 99.78 | 99.66 | 100.00 | 99.14 | 99.40 |
| 99.10 | 99.40 | 98.88 | 99.10 | 99.31 | 98.78 | 99.22 | 97.41 | 97.80 |
| 99.27 | 98.07 | 98.88 | 94.99 | 97.49 | 97.53 | 98.66 | 94.69 | 95.39 |
| 91.92 | 90.64 | 91.88 | 93.10 | 91.63 | 90.55 | 90.90 | 90.77 | 91.11 |
| 90.68 | 90.76 | 92.31 | 91.63 | 92.20 | 94.10 | 93.99 | 91.70 | 91.70 |
| 90.86 | 92.03 | 92.25 | 91.62 | 92.60 | 91.50 | 95.48 | 91.67 | 95.43 |
| 91.33 | 93.71 | 94.73 | 91.41 | 95.39 | 91.37 | 91.37 | 91.46 | 93.84 |
| 93.71 | 92.33 | 92.33 | 93.04 | 93.05 | 93.34 | 93.26 | 92.87 | 92.53 |
| 92.61 | 93.34 | 92.44 | 92.40 | 93.09 | 93.08 | 93.16 | 94.49 | 90.89 |
| 93.16 | 93.97 | 93.84 | 94.14 | 93.71 | 90.96 | 93.01 | 92.84 | 92.79 |
| 93.06 | 92.62 | 93.11 | 93.15 | 92.75 | 93.12 | 94.14 | 94.32 | 94.41 |

| | | | | | | | | |
|---|---|---|---|---|---|---|---|---|
| 93.93 | 93.84 | 93.94 | 93.94 | 93.89 | 93.75 | 93.66 | 93.84 | 93.75 |
| 93.92 | 93.88 | 93.75 | 93.79 | 92.21 | 93.84 | 89.69 | 91.92 | 90.67 |
| **5: Siamang-Gibbon** | | | 99.14 | 99.14 | 99.01 | 99.14 | 100.00 | 99.48 |
| 99.18 | 98.84 | 98.32 | 98.49 | 98.71 | 98.17 | 98.62 | 96.85 | 97.29 |
| 98.66 | 97.46 | 98.36 | 94.52 | 96.97 | 97.02 | 98.10 | 94.17 | 94.87 |
| 91.32 | 90.16 | 91.27 | 92.66 | 91.02 | 90.03 | 90.38 | 90.29 | 90.59 |
| 90.11 | 90.55 | 91.92 | 91.28 | 91.85 | 93.67 | 93.51 | 91.18 | 91.26 |
| 90.55 | 91.69 | 91.86 | 91.32 | 92.21 | 91.11 | 94.92 | 91.15 | 94.92 |
| 90.77 | 93.23 | 94.26 | 91.11 | 94.83 | 91.07 | 91.06 | 91.11 | 93.37 |
| 93.19 | 92.03 | 92.03 | 92.65 | 92.66 | 92.96 | 92.74 | 92.40 | 92.14 |
| 92.18 | 92.83 | 92.01 | 91.96 | 92.66 | 92.60 | 92.77 | 94.05 | 90.50 |
| 92.81 | 93.49 | 93.36 | 93.67 | 93.23 | 90.66 | 92.49 | 92.32 | 92.27 |
| 92.58 | 92.10 | 92.63 | 92.67 | 92.23 | 92.77 | 93.58 | 93.80 | 93.89 |
| 93.50 | 93.41 | 93.55 | 93.55 | 93.45 | 93.23 | 93.14 | 93.32 | 93.23 |
| 93.40 | 93.36 | 93.23 | 93.27 | 91.73 | 93.45 | 89.30 | 91.57 | 90.20 |
| **6: Silvery-Gibbon** | | | 99.40 | 99.40 | 99.27 | 99.40 | 99.48 | 100.00 |
| 99.44 | 99.10 | 98.54 | 98.75 | 98.97 | 98.43 | 98.88 | 97.11 | 97.59 |
| 98.92 | 97.74 | 98.62 | 94.69 | 97.23 | 97.28 | 98.36 | 94.43 | 95.13 |
| 91.53 | 90.33 | 91.49 | 92.84 | 91.24 | 90.25 | 90.59 | 90.46 | 90.80 |
| 90.33 | 90.68 | 92.09 | 91.37 | 92.02 | 93.84 | 93.69 | 91.44 | 91.53 |
| 90.68 | 91.82 | 92.08 | 91.49 | 92.43 | 91.33 | 95.17 | 91.41 | 95.13 |
| 91.03 | 93.40 | 94.47 | 91.28 | 95.09 | 91.20 | 91.24 | 91.20 | 93.58 |
| 93.40 | 92.16 | 92.16 | 92.83 | 92.83 | 93.13 | 92.96 | 92.57 | 92.31 |
| 92.31 | 93.04 | 92.18 | 92.14 | 92.87 | 92.73 | 92.90 | 94.27 | 90.67 |
| 92.94 | 93.66 | 93.53 | 93.84 | 93.40 | 90.87 | 92.66 | 92.49 | 92.44 |
| 92.76 | 92.27 | 92.80 | 92.85 | 92.40 | 92.86 | 93.75 | 93.97 | 94.06 |
| 93.67 | 93.58 | 93.73 | 93.73 | 93.63 | 93.49 | 93.40 | 93.58 | 93.49 |
| 93.66 | 93.62 | 93.49 | 93.53 | 91.95 | 93.62 | 89.48 | 91.61 | 90.37 |
| **7: Northern-Gibbon** | | | 99.10 | 99.10 | 98.97 | 99.10 | 99.18 | 99.44 |
| 100.00 | 98.79 | 98.23 | 98.45 | 98.66 | 98.13 | 98.58 | 96.98 | 97.41 |
| 98.62 | 97.46 | 98.31 | 94.52 | 97.10 | 97.15 | 98.01 | 94.26 | 94.96 |
| 91.49 | 90.38 | 91.45 | 92.84 | 91.15 | 90.29 | 90.64 | 90.51 | 90.80 |
| 90.24 | 90.59 | 92.05 | 91.28 | 91.94 | 93.76 | 93.60 | 91.31 | 91.48 |
| 90.60 | 91.73 | 92.03 | 91.41 | 92.34 | 91.29 | 95.00 | 91.24 | 94.96 |
| 90.81 | 93.32 | 94.43 | 91.20 | 94.92 | 91.20 | 91.11 | 91.11 | 93.50 |
| 93.32 | 92.07 | 92.07 | 92.74 | 92.74 | 93.04 | 92.87 | 92.40 | 92.22 |
| 92.14 | 92.96 | 92.01 | 92.05 | 92.79 | 92.82 | 92.86 | 94.01 | 90.50 |
| 92.90 | 93.66 | 93.49 | 93.75 | 93.32 | 90.74 | 92.71 | 92.53 | 92.49 |
| 92.80 | 92.27 | 92.85 | 92.89 | 92.45 | 92.86 | 93.71 | 93.93 | 94.02 |
| 93.58 | 93.50 | 93.64 | 93.64 | 93.54 | 93.27 | 93.19 | 93.36 | 93.27 |
| 93.45 | 93.40 | 93.27 | 93.32 | 91.77 | 93.45 | 89.48 | 91.74 | 90.20 |
| **8: Orangutan** | | | 99.44 | 99.44 | 99.31 | 99.40 | 98.84 | 99.10 |
| 98.79 | 100.00 | 98.58 | 98.79 | 99.01 | 98.52 | 98.97 | 97.15 | 97.63 |
| 98.97 | 97.88 | 98.62 | 94.73 | 97.36 | 97.40 | 98.40 | 94.47 | 95.22 |
| 91.71 | 90.55 | 91.66 | 92.97 | 91.37 | 90.46 | 90.81 | 90.68 | 91.02 |
| 90.51 | 90.63 | 92.13 | 91.46 | 92.11 | 93.97 | 93.86 | 91.44 | 91.48 |
| 90.73 | 91.82 | 92.16 | 91.49 | 92.47 | 91.37 | 95.26 | 91.54 | 95.22 |
| 91.11 | 93.62 | 94.69 | 91.28 | 95.17 | 91.24 | 91.24 | 91.33 | 93.71 |
| 93.62 | 92.16 | 92.16 | 92.91 | 92.92 | 93.22 | 93.13 | 92.74 | 92.44 |
| 92.48 | 93.22 | 92.27 | 92.22 | 92.96 | 92.99 | 93.16 | 94.23 | 90.80 |

| | | | | | | | | |
|---|---|---|---|---|---|---|---|---|
| 93.03 | 93.88 | 93.75 | 94.06 | 93.62 | 90.83 | 92.93 | 92.75 | 92.71 |
| 92.97 | 92.53 | 93.02 | 93.06 | 92.66 | 92.99 | 93.97 | 94.10 | 94.19 |
| 93.80 | 93.71 | 93.81 | 93.81 | 93.76 | 93.66 | 93.58 | 93.75 | 93.66 |
| 93.84 | 93.79 | 93.66 | 93.71 | 92.08 | 93.75 | 89.52 | 91.79 | 90.50 |
| **9: Snubnose-Monkey** | | | 98.88 | 98.88 | 98.75 | 98.88 | 98.32 | 98.54 |
| 98.23 | 98.58 | 100.00 | 98.92 | 99.18 | 98.61 | 99.18 | 96.77 | 97.11 |
| 99.10 | 97.98 | 98.70 | 94.60 | 96.80 | 96.84 | 99.18 | 94.26 | 94.87 |
| 91.40 | 90.07 | 91.36 | 92.75 | 91.15 | 90.07 | 90.42 | 90.33 | 90.59 |
| 90.24 | 90.63 | 91.87 | 91.37 | 92.07 | 93.54 | 93.43 | 90.96 | 91.57 |
| 90.73 | 91.73 | 91.99 | 91.36 | 92.25 | 91.16 | 94.87 | 91.24 | 94.83 |
| 91.16 | 93.14 | 94.26 | 91.28 | 94.79 | 91.15 | 91.19 | 91.20 | 93.41 |
| 93.19 | 92.03 | 92.03 | 92.44 | 92.44 | 92.74 | 92.65 | 92.35 | 91.97 |
| 92.10 | 92.74 | 92.05 | 92.01 | 92.61 | 92.60 | 92.81 | 93.93 | 90.37 |
| 92.77 | 93.40 | 93.27 | 93.58 | 93.19 | 90.61 | 92.62 | 92.45 | 92.40 |
| 92.63 | 92.27 | 92.72 | 92.76 | 92.36 | 92.73 | 93.49 | 93.67 | 93.76 |
| 93.37 | 93.37 | 93.38 | 93.38 | 93.32 | 93.19 | 93.10 | 93.27 | 93.19 |
| 93.36 | 93.32 | 93.19 | 93.23 | 91.69 | 93.27 | 89.21 | 91.48 | 90.28 |
| **10: Rhesus** | | | 99.10 | 99.10 | 98.97 | 99.10 | 98.49 | 98.75 |
| 98.45 | 98.79 | 98.92 | 100.00 | 99.78 | 99.35 | 99.70 | 96.94 | 97.37 |
| 99.31 | 98.12 | 99.18 | 94.65 | 97.06 | 97.10 | 98.70 | 94.34 | 94.96 |
| 91.40 | 90.20 | 91.36 | 92.62 | 91.19 | 90.20 | 90.55 | 90.38 | 90.80 |
| 90.29 | 90.50 | 92.05 | 91.37 | 91.98 | 93.80 | 93.69 | 91.27 | 91.35 |
| 90.60 | 91.69 | 92.03 | 91.36 | 92.30 | 91.20 | 95.05 | 91.15 | 95.00 |
| 90.98 | 93.45 | 94.30 | 91.11 | 94.96 | 91.11 | 91.11 | 91.20 | 93.45 |
| 93.32 | 92.16 | 92.16 | 92.61 | 92.66 | 92.91 | 92.83 | 92.48 | 92.14 |
| 92.22 | 92.91 | 91.96 | 91.92 | 92.66 | 92.73 | 92.73 | 94.05 | 90.72 |
| 92.73 | 93.53 | 93.40 | 93.71 | 93.36 | 90.74 | 92.62 | 92.45 | 92.40 |
| 92.67 | 92.27 | 92.80 | 92.76 | 92.36 | 92.68 | 93.62 | 93.80 | 93.89 |
| 93.58 | 93.54 | 93.60 | 93.60 | 93.58 | 93.45 | 93.32 | 93.49 | 93.40 |
| 93.58 | 93.53 | 93.40 | 93.45 | 91.90 | 93.49 | 89.61 | 91.57 | 90.41 |
| **11: Macaque** | | | 99.31 | 99.31 | 99.18 | 99.31 | 98.71 | 98.97 |
| 98.66 | 99.01 | 99.18 | 99.78 | 100.00 | 99.52 | 99.91 | 97.15 | 97.59 |
| 99.53 | 98.35 | 99.39 | 94.81 | 97.23 | 97.27 | 98.96 | 94.51 | 95.17 |
| 91.62 | 90.46 | 91.58 | 92.84 | 91.41 | 90.41 | 90.80 | 90.59 | 90.97 |
| 90.46 | 90.54 | 92.17 | 91.41 | 91.98 | 93.88 | 93.77 | 91.39 | 91.52 |
| 90.64 | 91.68 | 92.12 | 91.40 | 92.38 | 91.28 | 95.26 | 91.36 | 95.17 |
| 91.19 | 93.57 | 94.52 | 91.19 | 95.17 | 91.15 | 91.15 | 91.24 | 93.62 |
| 93.49 | 92.28 | 92.28 | 92.82 | 92.87 | 93.12 | 93.04 | 92.69 | 92.35 |
| 92.43 | 93.12 | 92.17 | 92.13 | 92.87 | 92.90 | 92.94 | 94.22 | 90.84 |
| 92.89 | 93.75 | 93.62 | 93.92 | 93.49 | 90.87 | 92.79 | 92.62 | 92.57 |
| 92.84 | 92.44 | 92.89 | 92.93 | 92.53 | 92.85 | 93.83 | 94.01 | 94.10 |
| 93.71 | 93.62 | 93.81 | 93.81 | 93.67 | 93.53 | 93.44 | 93.62 | 93.53 |
| 93.70 | 93.66 | 93.53 | 93.57 | 92.03 | 93.70 | 89.69 | 91.65 | 90.59 |
| **12: Pigtail-Macaque** | | | 98.78 | 98.78 | 98.65 | 98.78 | 98.17 | 98.43 |
| 98.13 | 98.52 | 98.61 | 99.35 | 99.52 | 100.00 | 99.43 | 96.78 | 97.13 |
| 99.00 | 98.63 | 99.65 | 94.38 | 97.44 | 97.49 | 99.18 | 94.12 | 94.78 |
| 91.58 | 90.22 | 91.58 | 92.98 | 91.29 | 90.17 | 90.52 | 90.30 | 90.72 |
| 90.59 | 90.48 | 92.10 | 91.44 | 91.76 | 93.60 | 93.67 | 91.27 | 91.43 |
| 90.62 | 91.42 | 91.90 | 91.35 | 92.17 | 91.14 | 94.91 | 91.37 | 94.70 |
| 90.80 | 93.59 | 94.03 | 91.14 | 94.74 | 91.10 | 91.06 | 91.31 | 93.47 |

| | | | | | | | | |
|---|---|---|---|---|---|---|---|---|
| 93.63 | 91.99 | 91.99 | 92.72 | 92.76 | 93.02 | 92.93 | 92.63 | 92.24 |
| 92.37 | 93.02 | 92.42 | 92.38 | 92.76 | 92.89 | 93.19 | 93.79 | 90.68 |
| 92.75 | 93.76 | 93.63 | 93.94 | 93.55 | 91.04 | 92.85 | 92.67 | 92.63 |
| 92.59 | 92.50 | 92.85 | 92.81 | 92.59 | 92.71 | 93.85 | 93.73 | 93.82 |
| 93.42 | 93.38 | 93.67 | 93.67 | 93.42 | 93.63 | 93.50 | 93.72 | 93.59 |
| 93.76 | 93.72 | 93.59 | 93.68 | 91.89 | 93.72 | 89.64 | 91.42 | 90.24 |
| **13: Facicularis-Macaque** | | | 99.22 | 99.22 | 99.09 | 99.22 | 98.62 | 98.88 |
| 98.58 | 98.97 | 99.18 | 99.70 | 99.91 | 99.43 | 100.00 | 97.07 | 97.50 |
| 99.44 | 98.26 | 99.31 | 94.73 | 97.14 | 97.19 | 98.87 | 94.43 | 95.08 |
| 91.53 | 90.37 | 91.49 | 92.75 | 91.32 | 90.33 | 90.72 | 90.50 | 90.88 |
| 90.37 | 90.59 | 92.08 | 91.41 | 92.02 | 93.79 | 93.68 | 91.30 | 91.52 |
| 90.68 | 91.73 | 92.16 | 91.45 | 92.42 | 91.32 | 95.17 | 91.27 | 95.08 |
| 91.15 | 93.49 | 94.43 | 91.24 | 95.08 | 91.19 | 91.19 | 91.28 | 93.54 |
| 93.40 | 92.19 | 92.19 | 92.73 | 92.78 | 93.04 | 92.95 | 92.61 | 92.26 |
| 92.35 | 93.04 | 92.09 | 92.04 | 92.78 | 92.81 | 92.94 | 94.14 | 90.75 |
| 92.81 | 93.66 | 93.53 | 93.84 | 93.40 | 90.78 | 92.71 | 92.53 | 92.48 |
| 92.75 | 92.35 | 92.80 | 92.84 | 92.44 | 92.76 | 93.75 | 93.93 | 94.01 |
| 93.62 | 93.54 | 93.72 | 93.72 | 93.58 | 93.44 | 93.36 | 93.53 | 93.44 |
| 93.62 | 93.57 | 93.44 | 93.49 | 91.94 | 93.62 | 89.61 | 91.56 | 90.50 |
| **14: Jacchus-Marmoset** | | | 97.37 | 97.37 | 97.24 | 97.41 | 96.85 | 97.11 |
| 96.98 | 97.15 | 96.77 | 96.94 | 97.15 | 96.78 | 97.07 | 100.00 | 98.45 |
| 97.07 | 95.90 | 96.75 | 93.91 | 98.01 | 98.14 | 96.58 | 93.91 | 94.35 |
| 90.88 | 89.81 | 90.84 | 92.19 | 90.67 | 89.68 | 90.07 | 89.86 | 90.33 |
| 90.06 | 90.11 | 91.35 | 90.85 | 91.81 | 92.84 | 92.77 | 90.88 | 90.96 |
| 90.25 | 91.47 | 91.52 | 90.84 | 91.82 | 90.64 | 94.44 | 90.80 | 94.39 |
| 90.42 | 92.53 | 93.82 | 90.85 | 94.35 | 90.81 | 90.63 | 90.76 | 93.02 |
| 93.01 | 91.81 | 91.81 | 92.13 | 92.17 | 92.43 | 92.43 | 92.17 | 91.87 |
| 92.04 | 92.52 | 91.39 | 91.35 | 92.26 | 92.34 | 92.20 | 93.36 | 89.89 |
| 92.16 | 92.80 | 92.66 | 92.93 | 92.58 | 90.39 | 92.40 | 92.19 | 92.14 |
| 92.54 | 91.97 | 92.37 | 92.46 | 92.10 | 92.12 | 92.97 | 93.02 | 93.19 |
| 92.67 | 92.59 | 92.68 | 92.68 | 92.72 | 92.62 | 92.58 | 92.71 | 92.71 |
| 92.80 | 92.75 | 92.62 | 92.71 | 91.08 | 92.58 | 89.21 | 91.14 | 89.85 |
| **15: Squirrel-Monkey** | | | 97.85 | 97.85 | 97.72 | 97.80 | 97.29 | 97.59 |
| 97.41 | 97.63 | 97.11 | 97.37 | 97.59 | 97.13 | 97.50 | 98.45 | 100.00 |
| 97.46 | 96.23 | 97.15 | 94.21 | 98.70 | 98.83 | 96.97 | 93.96 | 94.87 |
| 91.45 | 90.29 | 91.40 | 92.58 | 91.19 | 90.16 | 90.59 | 90.38 | 90.76 |
| 90.51 | 90.50 | 91.70 | 91.33 | 91.89 | 93.37 | 93.34 | 91.14 | 91.22 |
| 90.64 | 91.43 | 91.65 | 91.45 | 91.95 | 90.81 | 94.83 | 90.98 | 94.83 |
| 90.85 | 93.14 | 94.08 | 91.07 | 94.74 | 91.07 | 90.93 | 91.15 | 93.28 |
| 93.27 | 92.16 | 92.16 | 92.52 | 92.44 | 92.74 | 92.78 | 92.53 | 92.10 |
| 92.27 | 92.78 | 91.75 | 91.70 | 92.53 | 92.69 | 92.68 | 93.71 | 90.28 |
| 92.51 | 93.32 | 93.19 | 93.49 | 93.14 | 90.87 | 92.71 | 92.45 | 92.40 |
| 92.80 | 92.23 | 92.72 | 92.80 | 92.36 | 92.47 | 93.32 | 93.45 | 93.54 |
| 93.19 | 93.15 | 93.25 | 93.25 | 93.24 | 93.14 | 93.10 | 93.23 | 93.14 |
| 93.32 | 93.27 | 93.14 | 93.23 | 91.60 | 93.14 | 89.43 | 91.53 | 90.41 |
| **16: Green-Monkey** | | | 99.27 | 99.27 | 99.14 | 99.27 | 98.66 | 98.92 |
| 98.62 | 98.97 | 99.10 | 99.31 | 99.53 | 99.00 | 99.44 | 97.07 | 97.46 |
| 100.00 | 98.35 | 99.09 | 94.78 | 97.15 | 97.19 | 98.92 | 94.47 | 95.13 |
| 91.79 | 90.59 | 91.75 | 92.84 | 91.58 | 90.51 | 90.85 | 90.77 | 91.06 |
| 90.64 | 90.59 | 92.26 | 91.46 | 92.07 | 93.89 | 93.77 | 91.44 | 91.53 |

| | | | | | | | | |
|---|---|---|---|---|---|---|---|---|
| 90.68 | 91.77 | 92.12 | 91.45 | 92.43 | 91.33 | 95.22 | 91.50 | 95.26 |
| 91.20 | 93.49 | 94.52 | 91.24 | 95.13 | 91.20 | 91.19 | 91.33 | 93.67 |
| 93.53 | 92.37 | 92.37 | 92.87 | 92.87 | 93.17 | 93.09 | 92.74 | 92.40 |
| 92.48 | 93.17 | 92.22 | 92.18 | 92.92 | 92.95 | 92.99 | 94.18 | 90.67 |
| 92.94 | 93.75 | 93.62 | 93.93 | 93.53 | 90.96 | 92.93 | 92.75 | 92.71 |
| 92.97 | 92.57 | 93.02 | 93.06 | 92.66 | 92.90 | 93.92 | 94.10 | 94.19 |
| 93.71 | 93.63 | 93.73 | 93.73 | 93.67 | 93.53 | 93.45 | 93.62 | 93.53 |
| 93.71 | 93.66 | 93.53 | 93.58 | 92.03 | 93.62 | 89.74 | 91.74 | 90.67 |
| **17: Colobus-Primate** | | | 98.07 | 98.07 | 97.93 | 98.07 | 97.46 | 97.74 |
| 97.46 | 97.88 | 97.98 | 98.12 | 98.35 | 98.63 | 98.26 | 95.90 | 96.23 |
| 98.35 | 100.00 | 98.78 | 93.49 | 96.62 | 96.67 | 98.59 | 93.06 | 93.88 |
| 90.35 | 89.20 | 90.35 | 91.88 | 89.99 | 89.01 | 89.29 | 89.39 | 89.47 |
| 89.05 | 89.35 | 91.33 | 90.39 | 90.65 | 92.68 | 92.73 | 90.38 | 90.33 |
| 89.50 | 90.33 | 90.80 | 90.24 | 91.14 | 91.15 | 93.83 | 90.43 | 93.79 |
| 89.74 | 92.44 | 93.06 | 90.01 | 93.74 | 89.92 | 89.87 | 90.30 | 92.77 |
| 92.68 | 91.07 | 91.07 | 91.74 | 91.75 | 92.07 | 91.98 | 91.65 | 91.23 |
| 91.42 | 92.07 | 91.28 | 91.23 | 91.84 | 91.98 | 92.20 | 92.72 | 89.37 |
| 91.87 | 92.73 | 92.58 | 92.92 | 92.54 | 89.55 | 91.78 | 91.64 | 91.63 |
| 91.77 | 91.49 | 92.01 | 91.97 | 91.54 | 91.83 | 92.82 | 92.86 | 92.96 |
| 92.49 | 92.44 | 92.77 | 92.77 | 92.49 | 92.63 | 92.58 | 92.82 | 92.63 |
| 92.82 | 92.77 | 92.63 | 92.63 | 90.98 | 92.63 | 88.33 | 90.56 | 89.03 |
| **18: Gelada-Primate** | | | 98.88 | 98.88 | 98.75 | 98.88 | 98.36 | 98.62 |
| 98.31 | 98.62 | 98.70 | 99.18 | 99.39 | 99.65 | 99.31 | 96.75 | 97.15 |
| 99.09 | 98.78 | 100.00 | 94.45 | 97.50 | 97.54 | 99.27 | 94.15 | 94.85 |
| 91.58 | 90.21 | 91.58 | 92.97 | 91.28 | 90.16 | 90.47 | 90.34 | 90.71 |
| 90.38 | 90.57 | 92.17 | 91.48 | 91.89 | 93.71 | 93.75 | 91.30 | 91.55 |
| 90.71 | 91.59 | 92.03 | 91.43 | 92.29 | 91.27 | 94.90 | 91.42 | 94.81 |
| 90.97 | 93.45 | 94.10 | 91.22 | 94.81 | 91.18 | 91.18 | 91.35 | 93.62 |
| 93.58 | 91.97 | 91.97 | 92.80 | 92.80 | 93.10 | 93.02 | 92.72 | 92.37 |
| 92.46 | 93.10 | 92.46 | 92.42 | 92.85 | 92.97 | 93.18 | 93.87 | 90.71 |
| 92.79 | 93.66 | 93.53 | 93.84 | 93.45 | 90.66 | 92.75 | 92.58 | 92.53 |
| 92.71 | 92.40 | 92.93 | 92.88 | 92.49 | 92.75 | 93.75 | 93.84 | 93.92 |
| 93.58 | 93.49 | 93.70 | 93.70 | 93.53 | 93.49 | 93.40 | 93.62 | 93.49 |
| 93.66 | 93.62 | 93.49 | 93.49 | 91.97 | 93.58 | 89.68 | 91.50 | 90.36 |
| **19: Galago-Primate** | | | 95.03 | 94.99 | 94.91 | 94.99 | 94.52 | 94.69 |
| 94.52 | 94.73 | 94.60 | 94.65 | 94.81 | 94.38 | 94.73 | 93.91 | 94.21 |
| 94.78 | 93.49 | 94.45 | 100.00 | 93.67 | 93.80 | 94.54 | 97.75 | 94.77 |
| 90.35 | 89.36 | 90.30 | 92.18 | 90.26 | 89.49 | 89.66 | 89.27 | 89.97 |
| 89.32 | 89.89 | 90.82 | 90.67 | 91.15 | 92.66 | 92.54 | 90.39 | 90.74 |
| 89.81 | 91.03 | 91.11 | 90.53 | 91.29 | 90.33 | 93.87 | 90.66 | 93.91 |
| 90.20 | 92.35 | 93.52 | 90.58 | 93.78 | 90.45 | 90.45 | 90.71 | 92.62 |
| 92.44 | 91.41 | 91.32 | 91.60 | 91.65 | 91.90 | 91.94 | 91.43 | 91.17 |
| 91.30 | 92.03 | 91.36 | 91.40 | 91.86 | 91.76 | 92.15 | 92.83 | 89.78 |
| 92.15 | 92.79 | 92.66 | 92.88 | 92.70 | 90.25 | 92.00 | 91.79 | 91.78 |
| 91.88 | 91.61 | 92.10 | 91.97 | 91.83 | 92.06 | 92.70 | 92.80 | 92.88 |
| 92.49 | 92.40 | 92.63 | 92.63 | 92.49 | 92.74 | 92.57 | 92.74 | 92.70 |
| 92.74 | 92.70 | 92.57 | 92.61 | 90.85 | 92.66 | 88.72 | 91.12 | 89.79 |
| **20: Capuchin-Monkey** | | | 97.53 | 97.53 | 97.40 | 97.49 | 96.97 | 97.23 |
| 97.10 | 97.36 | 96.80 | 97.06 | 97.23 | 97.44 | 97.14 | 98.01 | 98.70 |
| 97.15 | 96.62 | 97.50 | 93.67 | 100.00 | 99.70 | 97.33 | 93.41 | 94.37 |

| | | | | | | | | |
|---|---|---|---|---|---|---|---|---|
| 91.23 | 89.99 | 91.23 | 92.49 | 90.80 | 89.86 | 90.21 | 90.08 | 90.62 |
| 90.21 | 90.40 | 91.73 | 90.96 | 91.54 | 93.02 | 93.10 | 91.04 | 91.08 |
| 90.49 | 91.12 | 91.42 | 91.17 | 91.73 | 90.79 | 94.77 | 90.90 | 94.68 |
| 90.54 | 92.88 | 93.59 | 91.05 | 94.68 | 90.96 | 90.75 | 90.88 | 93.10 |
| 93.23 | 91.89 | 91.89 | 92.37 | 92.29 | 92.59 | 92.54 | 92.33 | 91.94 |
| 92.07 | 92.63 | 92.07 | 92.03 | 92.37 | 92.58 | 92.75 | 93.22 | 90.06 |
| 92.32 | 93.19 | 93.06 | 93.36 | 92.97 | 90.48 | 92.58 | 92.32 | 92.27 |
| 92.62 | 92.10 | 92.66 | 92.66 | 92.23 | 92.27 | 93.14 | 93.14 | 93.23 |
| 92.88 | 92.84 | 92.97 | 92.97 | 92.93 | 93.06 | 93.01 | 93.19 | 93.14 |
| 93.23 | 93.19 | 93.06 | 93.10 | 91.41 | 92.93 | 89.38 | 91.29 | 89.84 |
| **21: Tufted-Capuchin** | | | 97.58 | 97.58 | 97.45 | 97.53 | 97.02 | 97.28 |
| 97.15 | 97.40 | 96.84 | 97.10 | 97.27 | 97.49 | 97.19 | 98.14 | 98.83 |
| 97.19 | 96.67 | 97.54 | 93.80 | 99.70 | 100.00 | 97.37 | 93.54 | 94.50 |
| 91.23 | 89.99 | 91.23 | 92.58 | 90.89 | 89.95 | 90.29 | 90.08 | 90.71 |
| 90.21 | 90.36 | 91.64 | 91.01 | 91.58 | 93.15 | 93.23 | 91.04 | 91.08 |
| 90.45 | 91.25 | 91.46 | 91.17 | 91.77 | 90.83 | 94.77 | 90.82 | 94.68 |
| 90.45 | 93.01 | 93.63 | 91.01 | 94.68 | 91.09 | 90.70 | 90.92 | 93.19 |
| 93.32 | 91.84 | 91.84 | 92.46 | 92.37 | 92.67 | 92.63 | 92.42 | 92.03 |
| 92.16 | 92.72 | 92.16 | 92.12 | 92.46 | 92.67 | 92.84 | 93.35 | 90.02 |
| 92.40 | 93.23 | 93.10 | 93.41 | 93.10 | 90.53 | 92.62 | 92.36 | 92.31 |
| 92.66 | 92.14 | 92.71 | 92.71 | 92.27 | 92.36 | 93.27 | 93.27 | 93.36 |
| 93.01 | 92.97 | 93.10 | 93.10 | 93.06 | 93.14 | 93.10 | 93.27 | 93.23 |
| 93.32 | 93.27 | 93.14 | 93.19 | 91.50 | 93.06 | 89.47 | 91.37 | 89.97 |
| **22: Francoisi-Langur** | | | 98.66 | 98.66 | 98.53 | 98.66 | 98.10 | 98.36 |
| 98.01 | 98.40 | 99.18 | 98.70 | 98.96 | 99.18 | 98.87 | 96.58 | 96.97 |
| 98.92 | 98.59 | 99.27 | 94.54 | 97.33 | 97.37 | 100.00 | 94.06 | 94.85 |
| 91.53 | 90.16 | 91.53 | 92.84 | 91.19 | 90.12 | 90.42 | 90.29 | 90.62 |
| 90.25 | 90.53 | 92.12 | 91.35 | 91.93 | 93.49 | 93.57 | 91.21 | 91.60 |
| 90.66 | 91.55 | 91.90 | 91.34 | 92.16 | 91.14 | 94.77 | 91.33 | 94.68 |
| 91.02 | 93.27 | 94.15 | 91.22 | 94.68 | 91.18 | 91.18 | 91.26 | 93.62 |
| 93.54 | 92.10 | 92.10 | 92.67 | 92.68 | 92.97 | 92.89 | 92.63 | 92.20 |
| 92.37 | 92.97 | 92.38 | 92.33 | 92.72 | 92.88 | 93.09 | 93.74 | 90.36 |
| 92.79 | 93.53 | 93.40 | 93.71 | 93.36 | 90.61 | 92.80 | 92.62 | 92.57 |
| 92.71 | 92.44 | 92.97 | 92.93 | 92.53 | 92.75 | 93.62 | 93.66 | 93.75 |
| 93.36 | 93.36 | 93.53 | 93.53 | 93.32 | 93.36 | 93.27 | 93.49 | 93.36 |
| 93.53 | 93.49 | 93.36 | 93.36 | 91.84 | 93.40 | 89.55 | 91.46 | 90.36 |
| **23: Loris** | | | 94.73 | 94.73 | 94.65 | 94.69 | 94.17 | 94.43 |
| 94.26 | 94.47 | 94.26 | 94.34 | 94.51 | 94.12 | 94.43 | 93.91 | 93.96 |
| 94.47 | 93.06 | 94.15 | 97.75 | 93.41 | 93.54 | 94.06 | 100.00 | 94.56 |
| 89.96 | 89.23 | 89.91 | 91.74 | 89.78 | 89.27 | 89.57 | 89.10 | 89.62 |
| 88.92 | 89.32 | 90.30 | 90.36 | 90.71 | 92.31 | 92.15 | 89.73 | 90.52 |
| 89.29 | 90.85 | 90.55 | 90.01 | 90.90 | 89.72 | 93.48 | 90.48 | 93.48 |
| 89.89 | 91.96 | 93.13 | 90.28 | 93.39 | 89.89 | 90.01 | 90.28 | 92.19 |
| 92.05 | 90.80 | 90.72 | 91.25 | 91.30 | 91.55 | 91.64 | 91.26 | 90.82 |
| 91.13 | 91.73 | 91.01 | 90.96 | 91.52 | 91.55 | 91.71 | 92.62 | 89.83 |
| 92.06 | 92.35 | 92.13 | 92.40 | 92.18 | 89.86 | 91.66 | 91.48 | 91.35 |
| 91.49 | 91.17 | 91.76 | 91.71 | 91.44 | 91.97 | 92.22 | 92.36 | 92.45 |
| 92.19 | 92.06 | 92.24 | 92.24 | 92.14 | 92.31 | 92.13 | 92.31 | 92.26 |
| 92.35 | 92.31 | 92.22 | 92.22 | 90.54 | 92.22 | 88.72 | 90.60 | 89.66 |

| | | | | | | | | |
|---|---|---|---|---|---|---|---|---|
| **24: Lemur** | | | 95.43 | 95.43 | 95.34 | 95.39 | 94.87 | 95.13 |
| 94.96 | 95.22 | 94.87 | 94.96 | 95.17 | 94.78 | 95.08 | 94.35 | 94.87 |
| 95.13 | 93.88 | 94.85 | 94.77 | 94.37 | 94.50 | 94.85 | 94.56 | 100.00 |
| 90.70 | 89.64 | 90.70 | 92.18 | 90.49 | 89.81 | 90.03 | 89.59 | 90.15 |
| 89.89 | 90.24 | 91.04 | 90.98 | 91.24 | 93.23 | 93.25 | 90.66 | 90.83 |
| 90.16 | 91.21 | 91.47 | 91.10 | 91.52 | 90.37 | 94.18 | 91.15 | 94.53 |
| 90.68 | 92.97 | 93.74 | 90.89 | 94.18 | 90.72 | 90.58 | 90.80 | 92.89 |
| 92.84 | 91.29 | 91.29 | 92.22 | 92.35 | 92.52 | 92.56 | 92.31 | 91.96 |
| 92.05 | 92.65 | 91.79 | 91.74 | 92.44 | 92.73 | 92.16 | 93.45 | 90.10 |
| 92.20 | 93.10 | 92.97 | 93.27 | 92.97 | 90.22 | 92.40 | 92.31 | 92.27 |
| 92.41 | 92.14 | 92.58 | 92.45 | 92.36 | 92.12 | 92.92 | 93.02 | 93.10 |
| 93.10 | 92.97 | 93.20 | 93.20 | 93.02 | 93.31 | 93.36 | 93.44 | 93.23 |
| 93.40 | 93.36 | 93.23 | 93.27 | 91.73 | 93.14 | 89.03 | 91.05 | 89.89 |

| | | | | | | | | |
|---|---|---|---|---|---|---|---|---|
| **25: Myotis-Bat** | | | 91.92 | 91.84 | 91.75 | 91.92 | 91.32 | 91.53 |
| 91.49 | 91.71 | 91.40 | 91.40 | 91.62 | 91.58 | 91.53 | 90.88 | 91.45 |
| 91.79 | 90.35 | 91.58 | 90.35 | 91.23 | 91.23 | 91.53 | 89.96 | 90.70 |
| 100.00 | 93.30 | 99.91 | 93.17 | 98.06 | 93.47 | 93.60 | 93.73 | 93.86 |
| 95.98 | 88.11 | 89.53 | 88.73 | 89.24 | 93.08 | 92.93 | 88.40 | 88.48 |
| 88.16 | 88.82 | 89.04 | 88.67 | 89.21 | 88.33 | 91.40 | 91.16 | 91.40 |
| 88.34 | 92.75 | 90.74 | 88.64 | 91.27 | 88.34 | 88.63 | 88.51 | 93.04 |
| 92.88 | 90.42 | 90.42 | 92.15 | 92.19 | 92.41 | 92.28 | 91.85 | 91.85 |
| 91.67 | 92.32 | 91.55 | 91.55 | 92.06 | 92.62 | 92.62 | 90.72 | 87.60 |
| 92.31 | 92.88 | 92.79 | 93.05 | 92.62 | 89.54 | 92.61 | 92.44 | 92.39 |
| 92.42 | 92.17 | 92.47 | 92.55 | 92.35 | 92.31 | 92.66 | 92.82 | 92.91 |
| 92.91 | 92.91 | 92.84 | 92.84 | 92.91 | 92.66 | 92.62 | 92.79 | 92.66 |
| 92.75 | 92.75 | 92.62 | 92.62 | 91.14 | 92.53 | 87.87 | 91.24 | 88.44 |

| | | | | | | | | |
|---|---|---|---|---|---|---|---|---|
| **26: Sturnira-Bat** | | | 90.72 | 90.64 | 90.51 | 90.64 | 90.16 | 90.33 |
| 90.38 | 90.55 | 90.07 | 90.20 | 90.46 | 90.22 | 90.37 | 89.81 | 90.29 |
| 90.59 | 89.20 | 90.21 | 89.36 | 89.99 | 89.99 | 90.16 | 89.23 | 89.64 |
| 93.30 | 100.00 | 93.25 | 92.34 | 93.12 | 97.20 | 96.85 | 98.19 | 95.54 |
| 91.84 | 86.86 | 88.32 | 87.78 | 87.70 | 91.70 | 91.56 | 87.46 | 87.40 |
| 86.95 | 87.54 | 87.63 | 87.38 | 87.98 | 86.96 | 90.25 | 89.92 | 90.38 |
| 87.00 | 91.57 | 89.83 | 87.52 | 90.16 | 87.39 | 87.55 | 87.56 | 91.96 |
| 92.09 | 89.46 | 89.46 | 90.91 | 90.92 | 91.17 | 91.04 | 90.61 | 90.57 |
| 90.61 | 91.09 | 90.17 | 90.17 | 90.83 | 91.52 | 91.39 | 89.19 | 86.54 |
| 91.12 | 91.96 | 91.96 | 92.21 | 91.61 | 88.64 | 91.66 | 91.92 | 91.86 |
| 91.53 | 91.65 | 91.65 | 91.70 | 91.96 | 91.25 | 91.74 | 91.66 | 91.75 |
| 91.53 | 91.49 | 91.73 | 91.73 | 91.49 | 91.79 | 91.83 | 92.00 | 91.83 |
| 91.96 | 91.96 | 91.87 | 91.83 | 90.00 | 91.57 | 86.87 | 90.52 | 87.49 |

| | | | | | | | | |
|---|---|---|---|---|---|---|---|---|
| **27: Daubentonii-Bat** | | | 91.88 | 91.79 | 91.71 | 91.88 | 91.27 | 91.49 |
| 91.45 | 91.66 | 91.36 | 91.36 | 91.58 | 91.58 | 91.49 | 90.84 | 91.40 |
| 91.75 | 90.35 | 91.58 | 90.30 | 91.23 | 91.23 | 91.53 | 89.91 | 90.70 |
| 99.91 | 93.25 | 100.00 | 93.22 | 98.06 | 93.51 | 93.56 | 93.69 | 93.82 |
| 96.03 | 88.11 | 89.53 | 88.73 | 89.24 | 93.08 | 92.97 | 88.44 | 88.53 |
| 88.16 | 88.82 | 89.04 | 88.67 | 89.21 | 88.33 | 91.36 | 91.16 | 91.36 |
| 88.34 | 92.75 | 90.74 | 88.64 | 91.23 | 88.34 | 88.63 | 88.51 | 92.99 |
| 92.83 | 90.38 | 90.38 | 92.15 | 92.19 | 92.41 | 92.28 | 91.85 | 91.85 |
| 91.67 | 92.32 | 91.55 | 91.55 | 92.06 | 92.62 | 92.62 | 90.72 | 87.64 |
| 92.31 | 92.79 | 92.70 | 92.96 | 92.57 | 89.50 | 92.57 | 92.39 | 92.34 |
| 92.38 | 92.13 | 92.42 | 92.51 | 92.31 | 92.31 | 92.62 | 92.78 | 92.86 |

| | | | | | | | | |
|---|---|---|---|---|---|---|---|---|
| 92.91 | 92.91 | 92.84 | 92.84 | 92.91 | 92.62 | 92.57 | 92.75 | 92.62 |
| 92.70 | 92.70 | 92.57 | 92.57 | 91.14 | 92.49 | 87.91 | 91.24 | 88.44 |
| **28: Flying-Fox** | | | 93.10 | 93.10 | 93.01 | 93.10 | 92.66 | 92.84 |
| 92.84 | 92.97 | 92.75 | 92.62 | 92.84 | 92.98 | 92.75 | 92.19 | 92.58 |
| 92.84 | 91.88 | 92.97 | 92.18 | 92.49 | 92.58 | 92.84 | 91.74 | 92.18 |
| 93.17 | 92.34 | 93.22 | 100.00 | 93.14 | 92.60 | 92.47 | 92.25 | 92.28 |
| 92.05 | 88.90 | 90.48 | 89.21 | 89.98 | 94.94 | 94.71 | 89.53 | 89.57 |
| 88.90 | 89.64 | 89.82 | 89.50 | 90.04 | 89.03 | 92.36 | 92.72 | 92.23 |
| 89.03 | 94.83 | 91.87 | 89.08 | 92.32 | 89.04 | 89.25 | 89.16 | 94.59 |
| 94.70 | 91.46 | 91.55 | 93.63 | 93.67 | 93.93 | 93.76 | 93.45 | 93.11 |
| 93.28 | 93.84 | 93.34 | 93.34 | 93.76 | 94.02 | 94.14 | 91.11 | 89.02 |
| 93.75 | 95.09 | 95.04 | 95.25 | 94.78 | 90.48 | 94.48 | 94.31 | 94.30 |
| 93.94 | 94.09 | 94.03 | 94.07 | 94.39 | 93.71 | 94.70 | 94.59 | 94.68 |
| 94.76 | 94.68 | 94.49 | 94.49 | 94.72 | 94.61 | 94.53 | 94.70 | 94.53 |
| 94.74 | 94.74 | 94.61 | 94.61 | 92.84 | 94.44 | 88.91 | 93.28 | 89.73 |
| **29: Brown-Bat** | | | 91.58 | 91.50 | 91.41 | 91.63 | 91.02 | 91.24 |
| 91.15 | 91.37 | 91.15 | 91.19 | 91.41 | 91.29 | 91.32 | 90.67 | 91.19 |
| 91.58 | 89.99 | 91.28 | 90.26 | 90.80 | 90.89 | 91.19 | 89.78 | 90.49 |
| 98.06 | 93.12 | 98.06 | 93.14 | 100.00 | 93.34 | 93.47 | 93.56 | 93.74 |
| 96.38 | 87.73 | 89.27 | 88.64 | 89.03 | 93.03 | 92.85 | 88.53 | 88.44 |
| 87.78 | 88.60 | 88.86 | 88.55 | 89.09 | 88.17 | 90.98 | 91.00 | 91.19 |
| 88.16 | 92.58 | 90.61 | 88.30 | 90.98 | 88.00 | 88.26 | 88.51 | 92.95 |
| 92.75 | 90.25 | 90.25 | 91.94 | 91.94 | 92.20 | 92.07 | 91.68 | 91.68 |
| 91.51 | 92.11 | 91.42 | 91.42 | 91.81 | 92.50 | 92.50 | 90.60 | 87.60 |
| 92.19 | 92.58 | 92.58 | 92.75 | 92.45 | 89.64 | 92.49 | 92.27 | 92.27 |
| 92.51 | 92.10 | 92.43 | 92.56 | 92.18 | 92.19 | 92.45 | 92.69 | 92.73 |
| 92.90 | 92.86 | 92.67 | 92.67 | 92.82 | 92.41 | 92.36 | 92.54 | 92.41 |
| 92.58 | 92.58 | 92.45 | 92.45 | 90.93 | 92.36 | 87.96 | 91.16 | 88.30 |
| **30: Spearnose-Bat** | | | 90.64 | 90.55 | 90.42 | 90.55 | 90.03 | 90.25 |
| 90.29 | 90.46 | 90.07 | 90.20 | 90.41 | 90.17 | 90.33 | 89.68 | 90.16 |
| 90.51 | 89.01 | 90.16 | 89.49 | 89.86 | 89.95 | 90.12 | 89.27 | 89.81 |
| 93.47 | 97.20 | 93.51 | 92.60 | 93.34 | 100.00 | 97.50 | 97.59 | 96.06 |
| 92.02 | 86.60 | 88.37 | 87.60 | 88.01 | 92.04 | 91.91 | 87.41 | 87.40 |
| 86.69 | 87.72 | 87.85 | 87.38 | 88.16 | 87.05 | 90.33 | 90.09 | 90.42 |
| 86.87 | 91.74 | 89.53 | 87.30 | 90.25 | 87.30 | 87.42 | 87.47 | 91.96 |
| 92.09 | 89.54 | 89.54 | 90.96 | 90.96 | 91.22 | 91.13 | 90.70 | 90.79 |
| 90.83 | 91.17 | 90.39 | 90.39 | 90.87 | 91.52 | 91.61 | 89.32 | 86.54 |
| 91.25 | 92.09 | 92.09 | 92.30 | 91.92 | 88.73 | 91.70 | 91.83 | 91.73 |
| 91.53 | 91.60 | 91.57 | 91.65 | 91.79 | 91.38 | 91.96 | 91.96 | 92.01 |
| 91.83 | 91.83 | 92.03 | 92.03 | 91.79 | 91.83 | 91.83 | 92.00 | 91.87 |
| 92.00 | 92.00 | 91.87 | 91.87 | 90.04 | 91.87 | 86.92 | 90.57 | 87.41 |
| **31: Vampire-Bat** | | | 90.98 | 90.90 | 90.77 | 90.90 | 90.38 | 90.59 |
| 90.64 | 90.81 | 90.42 | 90.55 | 90.80 | 90.52 | 90.72 | 90.07 | 90.59 |
| 90.85 | 89.29 | 90.47 | 89.66 | 90.21 | 90.29 | 90.42 | 89.57 | 90.03 |
| 93.60 | 96.85 | 93.56 | 92.47 | 93.47 | 97.50 | 100.00 | 97.20 | 96.28 |
| 92.02 | 86.77 | 88.54 | 87.65 | 88.14 | 92.00 | 91.82 | 87.41 | 87.40 |
| 86.87 | 87.72 | 87.89 | 87.46 | 88.20 | 87.09 | 90.33 | 90.22 | 90.38 |
| 86.91 | 91.79 | 89.88 | 87.52 | 90.20 | 87.34 | 87.64 | 87.30 | 92.44 |
| 92.44 | 89.76 | 89.76 | 91.17 | 91.22 | 91.39 | 91.39 | 90.83 | 90.87 |
| 90.83 | 91.43 | 90.52 | 90.52 | 91.18 | 91.69 | 91.82 | 89.41 | 87.02 |

| | | | | | | | | |
|---|---|---|---|---|---|---|---|---|
| 91.38 | 92.17 | 92.17 | 92.38 | 91.92 | 88.94 | 91.79 | 91.87 | 91.77 |
| 91.57 | 91.56 | 91.65 | 91.74 | 91.83 | 91.51 | 92.13 | 92.09 | 92.18 |
| 91.83 | 91.79 | 92.03 | 92.03 | 91.79 | 92.09 | 92.00 | 92.17 | 92.04 |
| 92.35 | 92.35 | 92.22 | 92.22 | 90.30 | 91.92 | 87.18 | 90.74 | 87.49 |
| **32: Fruit-Bat** | | | 90.85 | 90.77 | 90.64 | 90.77 | 90.29 | 90.46 |
| 90.51 | 90.68 | 90.33 | 90.38 | 90.59 | 90.30 | 90.50 | 89.86 | 90.38 |
| 90.77 | 89.39 | 90.34 | 89.27 | 90.08 | 90.08 | 90.29 | 89.10 | 89.59 |
| 93.73 | 98.19 | 93.69 | 92.25 | 93.56 | 97.59 | 97.20 | 100.00 | 95.89 |
| 92.15 | 87.21 | 88.37 | 88.08 | 88.05 | 91.83 | 91.56 | 87.46 | 87.53 |
| 87.30 | 87.85 | 87.85 | 87.81 | 88.24 | 87.31 | 90.33 | 89.92 | 90.51 |
| 87.13 | 91.57 | 89.79 | 87.91 | 90.25 | 87.69 | 87.77 | 87.86 | 92.26 |
| 92.31 | 89.46 | 89.46 | 90.91 | 90.87 | 91.17 | 91.09 | 90.57 | 90.57 |
| 90.61 | 91.13 | 90.30 | 90.30 | 90.87 | 91.61 | 91.43 | 89.28 | 86.46 |
| 91.34 | 91.92 | 91.92 | 92.12 | 91.57 | 88.86 | 91.74 | 91.92 | 91.82 |
| 91.61 | 91.60 | 91.65 | 91.74 | 91.87 | 91.47 | 91.70 | 91.70 | 91.79 |
| 91.66 | 91.62 | 91.73 | 91.73 | 91.62 | 91.70 | 91.79 | 91.92 | 91.74 |
| 91.92 | 91.87 | 91.74 | 91.74 | 90.00 | 91.61 | 87.05 | 90.52 | 87.62 |
| **33: Pteronotus-Bat** | | | 91.19 | 91.11 | 91.02 | 91.11 | 90.59 | 90.80 |
| 90.80 | 91.02 | 90.59 | 90.80 | 90.97 | 90.72 | 90.88 | 90.33 | 90.76 |
| 91.06 | 89.47 | 90.71 | 89.97 | 90.62 | 90.71 | 90.62 | 89.62 | 90.15 |
| 93.86 | 95.54 | 93.82 | 92.28 | 93.74 | 96.06 | 96.28 | 95.89 | 100.00 |
| 92.31 | 87.04 | 88.66 | 87.82 | 88.42 | 92.04 | 92.02 | 88.05 | 87.79 |
| 87.13 | 88.08 | 88.17 | 87.72 | 88.52 | 87.52 | 90.76 | 90.26 | 90.85 |
| 87.44 | 91.59 | 89.87 | 87.60 | 90.63 | 87.61 | 87.82 | 87.60 | 92.65 |
| 92.36 | 89.81 | 89.77 | 91.29 | 91.38 | 91.55 | 91.46 | 91.07 | 91.12 |
| 91.03 | 91.50 | 91.08 | 91.08 | 91.29 | 92.15 | 91.93 | 89.90 | 87.08 |
| 91.67 | 91.85 | 91.85 | 92.06 | 91.72 | 88.73 | 91.97 | 92.06 | 92.01 |
| 92.25 | 91.80 | 92.17 | 92.17 | 92.01 | 91.71 | 91.89 | 92.17 | 92.26 |
| 92.00 | 91.87 | 92.19 | 92.19 | 91.96 | 92.15 | 92.11 | 92.23 | 92.15 |
| 92.32 | 92.28 | 92.15 | 92.15 | 90.54 | 91.89 | 87.75 | 90.86 | 88.04 |
| **34: Bat** | | | 90.72 | 90.64 | 90.55 | 90.68 | 90.11 | 90.33 |
| 90.24 | 90.51 | 90.24 | 90.29 | 90.46 | 90.59 | 90.37 | 90.06 | 90.51 |
| 90.64 | 89.05 | 90.38 | 89.32 | 90.21 | 90.21 | 90.25 | 88.92 | 89.89 |
| 95.98 | 91.84 | 96.03 | 92.05 | 96.38 | 92.02 | 92.02 | 92.15 | 92.31 |
| 100.00 | 86.93 | 88.61 | 87.80 | 88.09 | 92.00 | 91.73 | 87.61 | 87.38 |
| 86.98 | 87.71 | 87.93 | 87.61 | 88.15 | 87.14 | 90.55 | 90.18 | 90.51 |
| 87.91 | 91.62 | 89.50 | 87.54 | 90.55 | 87.41 | 87.45 | 87.54 | 91.97 |
| 92.01 | 89.46 | 89.46 | 90.88 | 90.79 | 91.05 | 90.92 | 90.49 | 90.58 |
| 90.32 | 90.96 | 90.79 | 90.79 | 90.75 | 91.64 | 91.82 | 89.79 | 86.75 |
| 91.20 | 91.83 | 91.83 | 91.92 | 91.79 | 88.76 | 91.66 | 91.44 | 91.35 |
| 91.44 | 91.18 | 91.45 | 91.58 | 91.35 | 91.11 | 91.79 | 91.88 | 91.97 |
| 91.75 | 91.71 | 91.68 | 91.68 | 91.71 | 91.79 | 91.83 | 92.01 | 91.92 |
| 91.97 | 91.97 | 91.83 | 91.92 | 90.11 | 91.62 | 87.45 | 90.29 | 87.81 |
| **35: House-Mouse** | | | 90.76 | 90.81 | 90.72 | 90.76 | 90.55 | 90.68 |
| 90.59 | 90.63 | 90.63 | 90.50 | 90.54 | 90.48 | 90.59 | 90.11 | 90.50 |
| 90.59 | 89.35 | 90.57 | 89.89 | 90.40 | 90.36 | 90.53 | 89.32 | 90.24 |
| 88.11 | 86.86 | 88.11 | 88.90 | 87.73 | 86.60 | 86.77 | 87.21 | 87.04 |
| 86.93 | 100.00 | 89.22 | 93.65 | 94.22 | 90.07 | 89.84 | 88.22 | 91.21 |
| 99.40 | 94.05 | 94.36 | 96.24 | 94.23 | 93.44 | 90.77 | 87.77 | 90.90 |
| 88.59 | 89.73 | 91.24 | 96.42 | 90.77 | 96.81 | 96.20 | 93.48 | 89.21 |

| | | | | | | | | |
|---|---|---|---|---|---|---|---|---|
| 89.12 | 88.29 | 88.25 | 88.67 | 88.76 | 88.97 | 88.88 | 88.59 | 88.37 |
| 88.41 | 88.97 | 87.75 | 87.79 | 88.89 | 88.88 | 88.84 | 89.48 | 87.02 |
| 88.74 | 89.73 | 89.56 | 89.77 | 89.25 | 87.19 | 89.16 | 88.90 | 88.69 |
| 88.81 | 88.52 | 89.03 | 89.03 | 88.90 | 88.74 | 89.38 | 89.55 | 89.60 |
| 89.90 | 89.94 | 89.23 | 89.23 | 90.07 | 89.47 | 89.47 | 89.56 | 89.47 |
| 89.60 | 89.56 | 89.43 | 89.43 | 87.61 | 89.25 | 85.92 | 87.98 | 86.20 |
| **36: Degu-Mouse** | | | 92.26 | 92.31 | 92.18 | 92.31 | 91.92 | 92.09 |
| 92.05 | 92.13 | 91.87 | 92.05 | 92.17 | 92.10 | 92.08 | 91.35 | 91.70 |
| 92.26 | 91.33 | 92.17 | 90.82 | 91.73 | 91.64 | 92.12 | 90.30 | 91.04 |
| 89.53 | 88.32 | 89.53 | 90.48 | 89.27 | 88.37 | 88.54 | 88.37 | 88.66 |
| 88.61 | 89.22 | 100.00 | 89.83 | 90.02 | 91.43 | 91.26 | 93.39 | 90.00 |
| 89.19 | 90.07 | 90.20 | 89.84 | 90.17 | 89.36 | 92.09 | 89.14 | 92.27 |
| 89.20 | 91.05 | 91.69 | 89.53 | 92.01 | 89.71 | 89.58 | 89.57 | 91.09 |
| 91.05 | 89.63 | 89.67 | 90.27 | 90.40 | 90.57 | 90.27 | 90.10 | 89.70 |
| 89.88 | 90.35 | 89.52 | 89.56 | 90.10 | 90.52 | 90.38 | 91.07 | 88.20 |
| 90.21 | 91.31 | 91.09 | 91.27 | 90.96 | 88.93 | 90.65 | 90.61 | 90.53 |
| 90.28 | 90.40 | 90.49 | 90.41 | 90.57 | 90.21 | 91.01 | 91.04 | 91.13 |
| 91.13 | 91.22 | 90.86 | 90.86 | 91.26 | 90.96 | 90.79 | 90.96 | 91.01 |
| 90.96 | 90.92 | 90.79 | 90.88 | 89.25 | 90.70 | 87.09 | 89.48 | 87.98 |
| **37: Gerbil** | | | 91.63 | 91.63 | 91.63 | 91.63 | 91.28 | 91.37 |
| 91.28 | 91.46 | 91.37 | 91.37 | 91.41 | 91.44 | 91.41 | 90.85 | 91.33 |
| 91.46 | 90.39 | 91.48 | 90.67 | 90.96 | 91.01 | 91.35 | 90.36 | 90.98 |
| 88.73 | 87.78 | 88.73 | 89.21 | 88.64 | 87.60 | 87.65 | 88.08 | 87.82 |
| 87.80 | 93.65 | 89.83 | 100.00 | 94.39 | 90.55 | 90.32 | 89.40 | 91.74 |
| 93.74 | 94.36 | 94.53 | 94.60 | 94.66 | 93.74 | 91.63 | 88.38 | 91.85 |
| 89.51 | 90.08 | 91.72 | 94.48 | 91.59 | 94.30 | 94.34 | 98.83 | 89.86 |
| 89.73 | 89.51 | 89.47 | 89.75 | 89.80 | 90.06 | 90.01 | 89.63 | 89.45 |
| 89.37 | 90.10 | 88.62 | 88.67 | 89.93 | 89.84 | 89.66 | 90.74 | 88.33 |
| 89.74 | 90.03 | 89.86 | 90.11 | 89.73 | 88.44 | 90.07 | 89.94 | 89.94 |
| 89.97 | 89.77 | 90.23 | 90.14 | 89.90 | 89.74 | 89.82 | 89.99 | 90.03 |
| 90.29 | 90.34 | 89.92 | 89.92 | 90.42 | 89.99 | 89.90 | 90.03 | 89.90 |
| 90.08 | 90.03 | 89.90 | 89.95 | 88.14 | 89.82 | 86.80 | 88.85 | 87.03 |
| **38: Desert-Hamsater** | | | 92.20 | 92.20 | 92.11 | 92.20 | 91.85 | 92.02 |
| 91.94 | 92.11 | 92.07 | 91.98 | 91.98 | 91.76 | 92.02 | 91.81 | 91.89 |
| 92.07 | 90.65 | 91.89 | 91.15 | 91.54 | 91.58 | 91.93 | 90.71 | 91.24 |
| 89.24 | 87.70 | 89.24 | 89.98 | 89.03 | 88.01 | 88.14 | 88.05 | 88.42 |
| 88.09 | 94.22 | 90.02 | 94.39 | 100.00 | 91.34 | 91.15 | 89.36 | 92.42 |
| 94.31 | 97.19 | 96.46 | 94.74 | 96.50 | 95.35 | 92.16 | 89.00 | 92.24 |
| 89.52 | 90.94 | 92.19 | 94.74 | 92.11 | 94.78 | 94.83 | 94.31 | 90.74 |
| 90.64 | 89.46 | 89.41 | 90.20 | 90.21 | 90.46 | 90.38 | 89.99 | 89.99 |
| 89.86 | 90.46 | 88.77 | 88.81 | 90.29 | 90.46 | 90.06 | 91.10 | 87.78 |
| 89.93 | 90.68 | 90.51 | 90.76 | 90.38 | 88.57 | 90.60 | 90.55 | 90.54 |
| 90.48 | 90.37 | 90.75 | 90.75 | 90.55 | 89.93 | 90.64 | 90.65 | 90.78 |
| 91.08 | 91.17 | 90.54 | 90.54 | 91.26 | 90.68 | 90.47 | 90.64 | 90.68 |
| 90.77 | 90.73 | 90.60 | 90.64 | 88.80 | 90.47 | 86.95 | 89.16 | 87.24 |
| **39: Common-Mink** | | | 94.10 | 94.15 | 94.06 | 94.10 | 93.67 | 93.84 |
| 93.76 | 93.97 | 93.54 | 93.80 | 93.88 | 93.60 | 93.79 | 92.84 | 93.37 |
| 93.89 | 92.68 | 93.71 | 92.66 | 93.02 | 93.15 | 93.49 | 92.31 | 93.23 |
| 93.08 | 91.70 | 93.08 | 94.94 | 93.03 | 92.04 | 92.00 | 91.83 | 92.04 |
| 92.00 | 90.07 | 91.43 | 90.55 | 91.34 | 100.00 | 99.26 | 90.61 | 90.18 |

| | | | | | | | | |
|---|---|---|---|---|---|---|---|---|
| 89.98 | 91.00 | 90.96 | 90.72 | 91.09 | 90.16 | 93.10 | 92.85 | 93.15 |
| 90.00 | 99.13 | 92.70 | 90.37 | 93.02 | 90.29 | 90.46 | 90.33 | 95.17 |
| 95.24 | 92.04 | 92.04 | 94.50 | 94.42 | 94.72 | 94.59 | 94.37 | 93.98 |
| 94.11 | 94.63 | 93.52 | 93.47 | 94.42 | 94.37 | 94.45 | 92.94 | 90.03 |
| 94.28 | 98.10 | 97.92 | 98.27 | 97.79 | 91.11 | 94.72 | 94.41 | 94.33 |
| 94.33 | 94.11 | 94.37 | 94.50 | 94.33 | 94.28 | 97.71 | 97.80 | 97.84 |
| 99.35 | 99.61 | 97.40 | 97.40 | 99.61 | 97.10 | 96.93 | 97.10 | 97.14 |
| 97.14 | 97.14 | 97.01 | 97.01 | 95.06 | 97.36 | 89.85 | 93.33 | 90.66 |
| **40: Lutreola-Mink** | | | 93.99 | 94.03 | 93.94 | 93.99 | 93.51 | 93.69 |
| 93.60 | 93.86 | 93.43 | 93.69 | 93.77 | 93.67 | 93.68 | 92.77 | 93.34 |
| 93.77 | 92.73 | 93.75 | 92.54 | 93.10 | 93.23 | 93.57 | 92.15 | 93.25 |
| 92.93 | 91.56 | 92.97 | 94.71 | 92.85 | 91.91 | 91.82 | 91.56 | 92.02 |
| 91.73 | 89.84 | 91.26 | 90.32 | 91.15 | 99.26 | 100.00 | 90.56 | 90.08 |
| 89.80 | 90.82 | 90.86 | 90.53 | 91.00 | 90.02 | 93.08 | 92.84 | 93.08 |
| 89.80 | 98.57 | 92.54 | 90.19 | 93.04 | 90.14 | 90.27 | 90.14 | 94.89 |
| 94.84 | 91.76 | 91.76 | 94.49 | 94.40 | 94.71 | 94.58 | 94.40 | 93.97 |
| 94.15 | 94.62 | 93.68 | 93.64 | 94.36 | 94.49 | 94.70 | 92.97 | 89.81 |
| 94.48 | 97.61 | 97.44 | 97.79 | 97.35 | 90.75 | 94.36 | 94.10 | 94.06 |
| 94.19 | 93.84 | 94.23 | 94.36 | 94.02 | 94.48 | 97.31 | 97.49 | 97.53 |
| 98.87 | 99.31 | 97.50 | 97.50 | 99.48 | 96.75 | 96.57 | 96.75 | 96.75 |
| 96.79 | 96.79 | 96.66 | 96.66 | 95.26 | 96.92 | 89.69 | 93.45 | 90.46 |
| **41: Guinea-Pig** | | | 91.75 | 91.70 | 91.57 | 91.70 | 91.18 | 91.44 |
| 91.31 | 91.44 | 90.96 | 91.27 | 91.39 | 91.27 | 91.30 | 90.88 | 91.14 |
| 91.44 | 90.38 | 91.30 | 90.39 | 91.04 | 91.04 | 91.21 | 89.73 | 90.66 |
| 88.40 | 87.46 | 88.44 | 89.53 | 88.53 | 87.41 | 87.41 | 87.46 | 88.05 |
| 87.61 | 88.22 | 93.39 | 89.40 | 89.36 | 90.61 | 90.56 | 100.00 | 89.34 |
| 88.23 | 89.07 | 89.12 | 89.06 | 89.25 | 88.45 | 91.40 | 88.28 | 91.75 |
| 88.59 | 90.01 | 90.78 | 88.84 | 91.36 | 88.80 | 88.54 | 89.18 | 90.00 |
| 90.10 | 88.55 | 88.59 | 89.44 | 89.44 | 89.74 | 89.57 | 89.18 | 89.18 |
| 89.01 | 89.66 | 88.91 | 88.86 | 89.44 | 89.43 | 89.47 | 90.72 | 87.93 |
| 89.82 | 90.14 | 89.92 | 90.23 | 89.83 | 88.10 | 89.61 | 89.57 | 89.57 |
| 89.46 | 89.48 | 89.45 | 89.46 | 89.52 | 89.73 | 90.01 | 90.05 | 90.09 |
| 90.26 | 90.39 | 89.95 | 89.95 | 90.48 | 89.97 | 90.01 | 90.10 | 90.01 |
| 90.05 | 90.01 | 89.88 | 89.92 | 88.33 | 89.75 | 87.00 | 88.53 | 87.34 |
| **42: Blind-Mole** | | | 91.79 | 91.70 | 91.61 | 91.70 | 91.26 | 91.53 |
| 91.48 | 91.48 | 91.57 | 91.35 | 91.52 | 91.43 | 91.52 | 90.96 | 91.22 |
| 91.53 | 90.33 | 91.55 | 90.74 | 91.08 | 91.08 | 91.60 | 90.52 | 90.83 |
| 88.48 | 87.40 | 88.53 | 89.57 | 88.44 | 87.40 | 87.40 | 87.53 | 87.79 |
| 87.38 | 91.21 | 90.00 | 91.74 | 92.42 | 90.18 | 90.08 | 89.34 | 100.00 |
| 91.39 | 92.30 | 92.47 | 91.86 | 92.61 | 91.69 | 91.70 | 88.10 | 91.79 |
| 89.49 | 89.93 | 92.18 | 91.91 | 91.66 | 92.12 | 92.03 | 91.43 | 89.80 |
| 89.71 | 88.97 | 88.97 | 89.39 | 89.39 | 89.69 | 89.56 | 89.26 | 88.96 |
| 89.13 | 89.65 | 88.82 | 88.86 | 89.52 | 89.47 | 89.43 | 90.72 | 88.01 |
| 89.73 | 89.75 | 89.54 | 89.74 | 89.49 | 87.73 | 89.36 | 89.49 | 89.40 |
| 89.34 | 89.22 | 89.77 | 89.64 | 89.44 | 89.77 | 89.45 | 89.58 | 89.62 |
| 90.01 | 90.10 | 89.69 | 89.69 | 90.06 | 89.58 | 89.45 | 89.67 | 89.49 |
| 89.67 | 89.62 | 89.54 | 89.54 | 87.90 | 89.54 | 86.59 | 87.87 | 87.22 |
| **43: Caroli-Mouse** | | | 90.86 | 90.90 | 90.81 | 90.86 | 90.55 | 90.68 |
| 90.60 | 90.73 | 90.73 | 90.60 | 90.64 | 90.62 | 90.68 | 90.25 | 90.64 |
| 90.68 | 89.50 | 90.71 | 89.81 | 90.49 | 90.45 | 90.66 | 89.29 | 90.16 |

| | | | | | | | | |
|---|---|---|---|---|---|---|---|---|
| 88.16 | 86.95 | 88.16 | 88.90 | 87.78 | 86.69 | 86.87 | 87.30 | 87.13 |
| 86.98 | 99.40 | 89.19 | 93.74 | 94.31 | 89.98 | 89.80 | 88.23 | 91.39 |
| 100.00 | 94.15 | 94.32 | 96.29 | 94.23 | 93.40 | 90.82 | 87.61 | 90.95 |
| 88.82 | 89.64 | 91.20 | 96.42 | 90.82 | 96.94 | 96.33 | 93.57 | 89.21 |
| 89.12 | 88.13 | 88.09 | 88.63 | 88.73 | 88.94 | 88.81 | 88.51 | 88.29 |
| 88.33 | 88.89 | 87.80 | 87.85 | 88.81 | 88.85 | 88.75 | 89.58 | 87.07 |
| 88.66 | 89.51 | 89.34 | 89.55 | 89.04 | 86.97 | 89.08 | 88.90 | 88.81 |
| 88.88 | 88.64 | 89.15 | 89.11 | 88.86 | 88.66 | 89.25 | 89.43 | 89.47 |
| 89.82 | 89.86 | 89.19 | 89.19 | 89.99 | 89.43 | 89.43 | 89.51 | 89.43 |
| 89.56 | 89.51 | 89.38 | 89.38 | 87.62 | 89.17 | 85.80 | 87.94 | 86.12 |
| **44: Chinese-Hamster** | | | 91.95 | 91.95 | 91.86 | 92.03 | 91.69 | 91.82 |
| 91.73 | 91.82 | 91.73 | 91.69 | 91.68 | 91.42 | 91.73 | 91.47 | 91.43 |
| 91.77 | 90.33 | 91.59 | 91.03 | 91.12 | 91.25 | 91.55 | 90.85 | 91.21 |
| 88.82 | 87.54 | 88.82 | 89.64 | 88.60 | 87.72 | 87.72 | 87.85 | 88.08 |
| 87.71 | 94.05 | 90.07 | 94.36 | 97.19 | 91.00 | 90.82 | 89.07 | 92.30 |
| 94.15 | 100.00 | 96.46 | 94.66 | 96.46 | 95.31 | 92.12 | 88.84 | 92.30 |
| 89.49 | 90.48 | 92.20 | 94.66 | 92.08 | 94.57 | 94.62 | 94.31 | 90.05 |
| 89.96 | 89.38 | 89.34 | 89.70 | 89.74 | 90.00 | 89.91 | 89.66 | 89.49 |
| 89.57 | 90.00 | 88.87 | 88.91 | 89.83 | 90.04 | 89.86 | 91.07 | 87.67 |
| 89.68 | 90.39 | 90.13 | 90.38 | 90.04 | 88.28 | 90.17 | 90.13 | 90.07 |
| 89.97 | 89.90 | 90.41 | 90.37 | 90.13 | 89.68 | 90.30 | 90.36 | 90.40 |
| 90.75 | 90.83 | 90.21 | 90.21 | 90.92 | 90.30 | 90.09 | 90.26 | 90.30 |
| 90.39 | 90.35 | 90.22 | 90.26 | 88.43 | 90.13 | 86.70 | 88.87 | 86.91 |
| **45: Snow-Vole** | | | 92.25 | 92.25 | 92.16 | 92.25 | 91.86 | 92.08 |
| 92.03 | 92.16 | 91.99 | 92.03 | 92.12 | 91.90 | 92.16 | 91.52 | 91.65 |
| 92.12 | 90.80 | 92.03 | 91.11 | 91.42 | 91.46 | 91.90 | 90.55 | 91.47 |
| 89.04 | 87.63 | 89.04 | 89.82 | 88.86 | 87.85 | 87.89 | 87.85 | 88.17 |
| 87.93 | 94.36 | 90.20 | 94.53 | 96.46 | 90.96 | 90.86 | 89.12 | 92.47 |
| 94.32 | 96.46 | 100.00 | 94.83 | 98.83 | 97.70 | 92.34 | 88.80 | 92.38 |
| 89.58 | 90.57 | 92.59 | 94.79 | 92.21 | 94.79 | 95.09 | 94.36 | 90.40 |
| 90.35 | 89.38 | 89.38 | 89.87 | 89.92 | 90.17 | 90.17 | 89.87 | 89.61 |
| 89.79 | 90.26 | 88.78 | 88.83 | 90.05 | 90.39 | 89.94 | 91.32 | 88.23 |
| 89.68 | 90.48 | 90.30 | 90.56 | 90.17 | 88.19 | 90.22 | 90.26 | 90.20 |
| 90.15 | 90.03 | 90.46 | 90.37 | 90.26 | 89.64 | 90.39 | 90.40 | 90.44 |
| 90.83 | 90.83 | 90.38 | 90.38 | 90.92 | 90.39 | 90.26 | 90.43 | 90.43 |
| 90.52 | 90.48 | 90.35 | 90.48 | 88.60 | 90.26 | 86.91 | 88.79 | 87.22 |
| **46: Woodmouse** | | | 91.62 | 91.67 | 91.58 | 91.62 | 91.32 | 91.49 |
| 91.41 | 91.49 | 91.36 | 91.36 | 91.40 | 91.35 | 91.45 | 90.84 | 91.45 |
| 91.45 | 90.24 | 91.43 | 90.53 | 91.17 | 91.17 | 91.34 | 90.01 | 91.10 |
| 88.67 | 87.38 | 88.67 | 89.50 | 88.55 | 87.38 | 87.46 | 87.81 | 87.72 |
| 87.61 | 96.24 | 89.84 | 94.60 | 94.74 | 90.72 | 90.53 | 89.06 | 91.86 |
| 96.29 | 94.66 | 94.83 | 100.00 | 94.83 | 94.00 | 91.45 | 88.45 | 91.71 |
| 89.33 | 90.42 | 91.80 | 96.42 | 91.45 | 96.76 | 96.42 | 94.51 | 89.82 |
| 89.72 | 88.85 | 88.81 | 89.61 | 89.66 | 89.92 | 89.83 | 89.57 | 89.27 |
| 89.31 | 89.92 | 88.39 | 88.44 | 89.75 | 89.61 | 89.35 | 90.64 | 87.93 |
| 89.56 | 90.16 | 89.98 | 90.20 | 89.77 | 87.88 | 89.89 | 89.72 | 89.64 |
| 89.79 | 89.46 | 90.05 | 89.97 | 89.72 | 89.52 | 89.85 | 89.95 | 89.99 |
| 90.55 | 90.60 | 89.91 | 89.91 | 90.64 | 90.07 | 90.07 | 90.16 | 90.07 |
| 90.11 | 90.07 | 89.94 | 89.94 | 88.21 | 89.90 | 86.79 | 88.88 | 86.81 |

| | | | | | | | | |
|---|---|---|---|---|---|---|---|---|
| **47: Vole** | | | 92.60 | 92.60 | 92.51 | 92.60 | 92.21 | 92.43 |
| 92.34 | 92.47 | 92.25 | 92.30 | 92.38 | 92.17 | 92.42 | 91.82 | 91.95 |
| 92.43 | 91.14 | 92.29 | 91.29 | 91.73 | 91.77 | 92.16 | 90.90 | 91.52 |
| 89.21 | 87.98 | 89.21 | 90.04 | 89.09 | 88.16 | 88.20 | 88.24 | 88.52 |
| 88.15 | 94.23 | 90.17 | 94.66 | 96.50 | 91.09 | 91.00 | 89.25 | 92.61 |
| 94.23 | 96.46 | 98.83 | 94.83 | 100.00 | 97.96 | 92.69 | 88.89 | 92.65 |
| 89.58 | 90.70 | 92.90 | 94.88 | 92.56 | 94.79 | 95.18 | 94.45 | 90.53 |
| 90.49 | 89.56 | 89.56 | 90.05 | 90.10 | 90.35 | 90.35 | 90.01 | 89.88 |
| 89.92 | 90.44 | 89.05 | 89.09 | 90.18 | 90.56 | 90.21 | 91.54 | 88.54 |
| 89.99 | 90.66 | 90.48 | 90.74 | 90.31 | 88.37 | 90.31 | 90.35 | 90.30 |
| 90.24 | 90.12 | 90.55 | 90.46 | 90.35 | 89.95 | 90.70 | 90.71 | 90.75 |
| 90.92 | 90.97 | 90.56 | 90.56 | 91.06 | 90.61 | 90.48 | 90.66 | 90.66 |
| 90.79 | 90.74 | 90.70 | 90.74 | 88.82 | 90.48 | 87.05 | 89.14 | 87.52 |

| | | | | | | | | |
|---|---|---|---|---|---|---|---|---|
| **48: Prairie-Vole** | | | 91.50 | 91.50 | 91.42 | 91.50 | 91.11 | 91.33 |
| 91.29 | 91.37 | 91.16 | 91.20 | 91.28 | 91.14 | 91.32 | 90.64 | 90.81 |
| 91.33 | 91.15 | 91.27 | 90.33 | 90.79 | 90.83 | 91.14 | 89.72 | 90.37 |
| 88.33 | 86.96 | 88.33 | 89.03 | 88.17 | 87.05 | 87.09 | 87.31 | 87.52 |
| 87.14 | 93.44 | 89.36 | 93.74 | 95.35 | 90.16 | 90.02 | 88.45 | 91.69 |
| 93.40 | 95.31 | 97.70 | 94.00 | 97.96 | 100.00 | 91.64 | 87.99 | 91.59 |
| 88.60 | 89.73 | 91.67 | 94.04 | 91.55 | 94.00 | 94.34 | 93.61 | 89.56 |
| 89.43 | 88.65 | 88.65 | 89.11 | 89.15 | 89.41 | 89.41 | 89.07 | 88.89 |
| 88.98 | 89.50 | 88.10 | 88.15 | 89.28 | 89.45 | 89.14 | 90.35 | 87.60 |
| 88.97 | 89.73 | 89.56 | 89.77 | 89.43 | 87.45 | 89.42 | 89.38 | 89.29 |
| 89.41 | 89.12 | 89.67 | 89.59 | 89.38 | 88.92 | 89.69 | 89.74 | 89.78 |
| 90.00 | 90.04 | 89.62 | 89.62 | 90.13 | 89.60 | 89.47 | 89.64 | 89.64 |
| 89.77 | 89.73 | 89.60 | 89.69 | 87.88 | 89.47 | 86.10 | 88.24 | 86.38 |

| | | | | | | | | |
|---|---|---|---|---|---|---|---|---|
| **49: Arctic-Squirrel** | | | 95.48 | 95.48 | 95.43 | 95.48 | 94.92 | 95.17 |
| 95.00 | 95.26 | 94.87 | 95.05 | 95.26 | 94.91 | 95.17 | 94.44 | 94.83 |
| 95.22 | 93.83 | 94.90 | 93.87 | 94.77 | 94.77 | 94.77 | 93.48 | 94.18 |
| 91.40 | 90.25 | 91.36 | 92.36 | 90.98 | 90.33 | 90.33 | 90.33 | 90.76 |
| 90.55 | 90.77 | 92.09 | 91.63 | 92.16 | 93.10 | 93.08 | 91.40 | 91.70 |
| 90.82 | 92.12 | 92.34 | 91.45 | 92.69 | 91.64 | 100.00 | 91.24 | 98.54 |
| 91.38 | 92.62 | 94.65 | 91.59 | 99.53 | 91.63 | 91.50 | 91.63 | 93.11 |
| 93.01 | 91.94 | 91.94 | 92.39 | 92.35 | 92.65 | 92.61 | 92.18 | 92.01 |
| 92.01 | 92.70 | 91.88 | 91.88 | 92.61 | 92.86 | 92.55 | 93.84 | 90.11 |
| 92.47 | 93.14 | 92.97 | 93.19 | 92.84 | 90.66 | 92.71 | 92.40 | 92.36 |
| 92.72 | 92.18 | 92.72 | 92.76 | 92.36 | 92.38 | 92.93 | 93.06 | 93.15 |
| 92.89 | 92.89 | 92.82 | 92.82 | 92.93 | 92.97 | 92.93 | 93.06 | 93.06 |
| 93.14 | 93.10 | 92.97 | 93.19 | 91.47 | 92.62 | 89.26 | 90.92 | 89.59 |

| | | | | | | | | |
|---|---|---|---|---|---|---|---|---|
| **50: Spanish-Mole** | | | 91.63 | 91.71 | 91.54 | 91.67 | 91.15 | 91.41 |
| 91.24 | 91.54 | 91.24 | 91.15 | 91.36 | 91.37 | 91.27 | 90.80 | 90.98 |
| 91.50 | 90.43 | 91.42 | 90.66 | 90.90 | 90.82 | 91.33 | 90.48 | 91.15 |
| 91.16 | 89.92 | 91.16 | 92.72 | 91.00 | 90.09 | 90.22 | 89.92 | 90.26 |
| 90.18 | 87.77 | 89.14 | 88.38 | 89.00 | 92.85 | 92.84 | 88.28 | 88.10 |
| 87.61 | 88.84 | 88.80 | 88.45 | 88.89 | 87.99 | 91.24 | 100.00 | 91.45 |
| 88.64 | 92.59 | 90.92 | 88.38 | 91.28 | 87.99 | 88.21 | 88.38 | 92.94 |
| 92.81 | 90.05 | 90.05 | 92.59 | 92.68 | 92.89 | 92.72 | 92.12 | 92.20 |
| 91.99 | 92.80 | 91.82 | 91.77 | 92.51 | 93.10 | 92.71 | 90.58 | 87.80 |
| 92.71 | 92.98 | 92.98 | 93.20 | 92.89 | 89.14 | 93.02 | 92.85 | 92.85 |
| 92.90 | 92.76 | 93.16 | 93.12 | 92.85 | 92.58 | 92.72 | 92.94 | 92.94 |

| | | | | | | | | |
|---|---|---|---|---|---|---|---|---|
| 92.81 | 92.77 | 92.84 | 92.84 | 92.81 | 92.55 | 92.46 | 92.59 | 92.55 |
| 92.72 | 92.72 | 92.59 | 92.68 | 91.07 | 92.55 | 87.40 | 90.86 | 88.34 |
| **51: Squirrel** | | | 95.43 | 95.43 | 95.39 | 95.43 | 94.92 | 95.13 |
| 94.96 | 95.22 | 94.83 | 95.00 | 95.17 | 94.70 | 95.08 | 94.39 | 94.83 |
| 95.26 | 93.79 | 94.81 | 93.91 | 94.68 | 94.68 | 94.68 | 93.48 | 94.53 |
| 91.40 | 90.38 | 91.36 | 92.23 | 91.19 | 90.42 | 90.38 | 90.51 | 90.85 |
| 90.51 | 90.90 | 92.27 | 91.85 | 92.24 | 93.15 | 93.08 | 91.75 | 91.79 |
| 90.95 | 92.30 | 92.38 | 91.71 | 92.65 | 91.59 | 98.54 | 91.45 | 100.00 |
| 91.29 | 92.66 | 94.61 | 91.68 | 98.49 | 91.68 | 91.63 | 91.81 | 93.02 |
| 92.93 | 91.81 | 91.81 | 92.31 | 92.31 | 92.61 | 92.52 | 92.18 | 91.97 |
| 92.01 | 92.61 | 91.83 | 91.83 | 92.48 | 92.73 | 92.55 | 93.67 | 90.16 |
| 92.47 | 93.10 | 92.88 | 93.15 | 92.88 | 90.74 | 92.75 | 92.45 | 92.40 |
| 92.85 | 92.31 | 92.85 | 92.85 | 92.45 | 92.38 | 93.06 | 93.19 | 93.28 |
| 92.93 | 92.93 | 92.86 | 92.86 | 93.06 | 93.01 | 92.97 | 93.10 | 93.10 |
| 93.10 | 93.06 | 92.93 | 93.06 | 91.47 | 92.62 | 89.26 | 91.05 | 89.76 |
| **52: Pacific-Mouse** | | | 91.33 | 91.33 | 91.29 | 91.33 | 90.77 | 91.03 |
| 90.81 | 91.11 | 91.16 | 90.98 | 91.19 | 90.80 | 91.15 | 90.42 | 90.85 |
| 91.20 | 89.74 | 90.97 | 90.20 | 90.54 | 90.45 | 91.02 | 89.89 | 90.68 |
| 88.34 | 87.00 | 88.34 | 89.03 | 88.16 | 86.87 | 86.91 | 87.13 | 87.44 |
| 87.91 | 88.59 | 89.20 | 89.51 | 89.52 | 90.00 | 89.80 | 88.59 | 89.49 |
| 88.82 | 89.49 | 89.58 | 89.33 | 89.58 | 88.60 | 91.38 | 88.64 | 91.29 |
| 100.00 | 89.77 | 91.98 | 89.08 | 91.38 | 88.91 | 88.91 | 89.16 | 89.62 |
| 89.56 | 88.62 | 88.62 | 89.07 | 89.08 | 89.38 | 89.29 | 88.82 | 88.64 |
| 88.64 | 89.38 | 88.71 | 88.67 | 89.21 | 89.18 | 89.23 | 90.65 | 87.55 |
| 89.35 | 89.64 | 89.51 | 89.73 | 89.56 | 87.51 | 89.46 | 89.42 | 89.33 |
| 89.36 | 89.16 | 89.44 | 89.44 | 89.33 | 89.26 | 89.21 | 89.36 | 89.44 |
| 89.97 | 89.75 | 89.66 | 89.66 | 89.79 | 89.73 | 89.69 | 89.77 | 89.77 |
| 89.86 | 89.82 | 89.69 | 89.73 | 88.04 | 89.51 | 86.08 | 87.84 | 86.86 |
| **53: Badger** | | | 93.71 | 93.75 | 93.66 | 93.71 | 93.23 | 93.40 |
| 93.32 | 93.62 | 93.14 | 93.45 | 93.57 | 93.59 | 93.49 | 92.53 | 93.14 |
| 93.49 | 92.44 | 93.45 | 92.35 | 92.88 | 93.01 | 93.27 | 91.96 | 92.97 |
| 92.75 | 91.57 | 92.75 | 94.83 | 92.58 | 91.74 | 91.79 | 91.57 | 91.59 |
| 91.62 | 89.73 | 91.05 | 90.08 | 90.94 | 99.13 | 98.57 | 90.01 | 89.93 |
| 89.64 | 90.48 | 90.57 | 90.42 | 90.70 | 89.73 | 92.62 | 92.59 | 92.66 |
| 89.77 | 100.00 | 92.39 | 90.03 | 92.58 | 89.91 | 90.16 | 89.86 | 94.72 |
| 95.01 | 91.86 | 91.86 | 94.10 | 94.06 | 94.32 | 94.19 | 93.93 | 93.63 |
| 93.71 | 94.23 | 93.25 | 93.21 | 94.02 | 94.15 | 94.40 | 92.46 | 89.81 |
| 93.97 | 97.93 | 97.76 | 98.14 | 97.63 | 90.66 | 94.44 | 94.18 | 94.17 |
| 93.94 | 93.96 | 94.03 | 94.07 | 94.09 | 93.97 | 97.55 | 97.32 | 97.40 |
| 98.92 | 98.88 | 97.14 | 97.14 | 98.88 | 96.94 | 96.86 | 97.03 | 97.03 |
| 97.07 | 97.07 | 96.94 | 96.94 | 94.75 | 97.42 | 89.48 | 93.11 | 90.21 |
| **54: Beaver** | | | 94.73 | 94.78 | 94.65 | 94.73 | 94.26 | 94.47 |
| 94.43 | 94.69 | 94.26 | 94.30 | 94.52 | 94.03 | 94.43 | 93.82 | 94.08 |
| 94.52 | 93.06 | 94.10 | 93.52 | 93.59 | 93.63 | 94.15 | 93.13 | 93.74 |
| 90.74 | 89.83 | 90.74 | 91.87 | 90.61 | 89.53 | 89.88 | 89.79 | 89.87 |
| 89.50 | 91.24 | 91.69 | 91.72 | 92.19 | 92.70 | 92.54 | 90.78 | 92.18 |
| 91.20 | 92.20 | 92.59 | 91.80 | 92.90 | 91.67 | 94.65 | 90.92 | 94.61 |
| 91.98 | 92.39 | 100.00 | 91.80 | 94.61 | 91.67 | 91.84 | 91.54 | 92.44 |
| 92.35 | 91.45 | 91.45 | 91.73 | 91.69 | 92.03 | 91.86 | 91.56 | 91.43 |
| 91.43 | 91.94 | 90.65 | 90.70 | 91.77 | 91.85 | 91.89 | 93.35 | 90.05 |

| | | | | | | | | |
|---|---|---|---|---|---|---|---|---|
| 92.10 | 92.66 | 92.44 | 92.66 | 92.22 | 90.21 | 92.05 | 92.05 | 91.96 |
| 91.80 | 91.78 | 92.02 | 92.02 | 91.96 | 92.10 | 92.18 | 92.36 | 92.44 |
| 92.57 | 92.49 | 92.24 | 92.24 | 92.57 | 92.18 | 92.05 | 92.22 | 92.13 |
| 92.35 | 92.31 | 92.18 | 92.22 | 90.63 | 92.13 | 88.90 | 90.42 | 89.54 |
| **55: Nile-Rat** | | | 91.41 | 91.46 | 91.37 | 91.41 | 91.11 | 91.28 |
| 91.20 | 91.28 | 91.28 | 91.11 | 91.19 | 91.14 | 91.24 | 90.85 | 91.07 |
| 91.24 | 90.01 | 91.22 | 90.58 | 91.05 | 91.01 | 91.22 | 90.28 | 90.89 |
| 88.64 | 87.52 | 88.64 | 89.08 | 88.30 | 87.30 | 87.52 | 87.91 | 87.60 |
| 87.54 | 96.42 | 89.53 | 94.48 | 94.74 | 90.37 | 90.19 | 88.84 | 91.91 |
| 96.42 | 94.66 | 94.79 | 96.42 | 94.88 | 94.04 | 91.59 | 88.38 | 91.68 |
| 89.08 | 90.03 | 91.80 | 100.00 | 91.59 | 96.72 | 96.76 | 94.22 | 89.69 |
| 89.60 | 88.73 | 88.69 | 89.45 | 89.45 | 89.67 | 89.54 | 89.15 | 89.06 |
| 89.02 | 89.62 | 88.32 | 88.36 | 89.41 | 89.45 | 89.31 | 90.43 | 87.38 |
| 89.31 | 90.03 | 89.86 | 90.03 | 89.56 | 87.79 | 89.77 | 89.47 | 89.42 |
| 89.45 | 89.25 | 89.75 | 89.80 | 89.47 | 89.31 | 89.82 | 89.99 | 90.03 |
| 90.25 | 90.25 | 89.75 | 89.75 | 90.29 | 89.77 | 89.77 | 89.86 | 89.77 |
| 89.90 | 89.86 | 89.73 | 89.73 | 87.96 | 89.69 | 86.71 | 88.45 | 86.90 |
| **56: Marmot-Rodent** | | | 95.39 | 95.39 | 95.30 | 95.39 | 94.83 | 95.09 |
| 94.92 | 95.17 | 94.79 | 94.96 | 95.17 | 94.74 | 95.08 | 94.35 | 94.74 |
| 95.13 | 93.74 | 94.81 | 93.78 | 94.68 | 94.68 | 94.68 | 93.39 | 94.18 |
| 91.27 | 90.16 | 91.23 | 92.32 | 90.98 | 90.25 | 90.20 | 90.25 | 90.63 |
| 90.55 | 90.77 | 92.01 | 91.59 | 92.11 | 93.02 | 93.04 | 91.36 | 91.66 |
| 90.82 | 92.08 | 92.21 | 91.45 | 92.56 | 91.55 | 99.53 | 91.28 | 98.49 |
| 91.38 | 92.58 | 94.61 | 91.59 | 100.00 | 91.55 | 91.59 | 91.59 | 92.98 |
| 92.93 | 91.68 | 91.68 | 92.31 | 92.27 | 92.57 | 92.61 | 92.10 | 91.92 |
| 91.92 | 92.70 | 91.70 | 91.70 | 92.57 | 92.78 | 92.42 | 93.67 | 90.07 |
| 92.42 | 93.14 | 92.97 | 93.19 | 92.80 | 90.35 | 92.62 | 92.32 | 92.27 |
| 92.63 | 92.10 | 92.63 | 92.68 | 92.32 | 92.34 | 92.88 | 92.98 | 93.06 |
| 92.80 | 92.80 | 92.73 | 92.73 | 92.85 | 92.88 | 92.84 | 92.97 | 92.97 |
| 93.06 | 93.01 | 92.88 | 93.01 | 91.39 | 92.53 | 89.17 | 90.83 | 89.59 |
| **57: Mastomys-Mouse** | | | 91.37 | 91.41 | 91.33 | 91.37 | 91.07 | 91.20 |
| 91.20 | 91.24 | 91.15 | 91.11 | 91.15 | 91.10 | 91.19 | 90.81 | 91.07 |
| 91.20 | 89.92 | 91.18 | 90.45 | 90.96 | 91.09 | 91.18 | 89.89 | 90.72 |
| 88.34 | 87.39 | 88.34 | 89.04 | 88.00 | 87.30 | 87.34 | 87.69 | 87.61 |
| 87.41 | 96.81 | 89.71 | 94.30 | 94.78 | 90.29 | 90.14 | 88.80 | 92.12 |
| 96.94 | 94.57 | 94.79 | 96.76 | 94.79 | 94.00 | 91.63 | 87.99 | 91.68 |
| 88.91 | 89.91 | 91.67 | 96.72 | 91.55 | 100.00 | 96.80 | 94.09 | 89.52 |
| 89.48 | 88.64 | 88.60 | 89.14 | 89.24 | 89.45 | 89.36 | 89.06 | 88.85 |
| 88.93 | 89.45 | 88.27 | 88.32 | 89.24 | 89.45 | 89.23 | 90.22 | 87.55 |
| 89.27 | 89.95 | 89.78 | 89.99 | 89.52 | 87.40 | 89.64 | 89.47 | 89.42 |
| 89.53 | 89.25 | 89.71 | 89.75 | 89.42 | 89.35 | 89.69 | 89.74 | 89.78 |
| 90.04 | 90.17 | 89.66 | 89.66 | 90.21 | 89.78 | 89.82 | 89.91 | 89.82 |
| 89.91 | 89.87 | 89.74 | 89.74 | 88.01 | 89.65 | 86.33 | 88.15 | 86.77 |
| **58: Rat** | | | 91.37 | 91.37 | 91.28 | 91.37 | 91.06 | 91.24 |
| 91.11 | 91.24 | 91.19 | 91.11 | 91.15 | 91.06 | 91.19 | 90.63 | 90.93 |
| 91.19 | 89.87 | 91.18 | 90.45 | 90.75 | 90.70 | 91.18 | 90.01 | 90.58 |
| 88.63 | 87.55 | 88.63 | 89.25 | 88.26 | 87.42 | 87.64 | 87.77 | 87.82 |
| 87.45 | 96.20 | 89.58 | 94.34 | 94.83 | 90.46 | 90.27 | 88.54 | 92.03 |
| 96.33 | 94.62 | 95.09 | 96.42 | 95.18 | 94.34 | 91.50 | 88.21 | 91.63 |
| 88.91 | 90.16 | 91.84 | 96.76 | 91.59 | 96.80 | 100.00 | 94.17 | 89.56 |

| | | | | | | | | |
|---|---|---|---|---|---|---|---|---|
| 89.51 | 88.99 | 88.94 | 89.19 | 89.28 | 89.49 | 89.41 | 89.06 | 88.85 |
| 88.93 | 89.49 | 88.36 | 88.40 | 89.28 | 89.27 | 89.18 | 90.21 | 87.37 |
| 89.18 | 90.03 | 89.86 | 90.08 | 89.55 | 87.75 | 89.42 | 89.20 | 89.13 |
| 89.28 | 88.96 | 89.45 | 89.49 | 89.20 | 89.18 | 89.77 | 89.87 | 89.91 |
| 90.34 | 90.30 | 89.79 | 89.79 | 90.39 | 89.81 | 89.77 | 89.94 | 89.86 |
| 89.99 | 89.94 | 89.81 | 89.81 | 88.01 | 89.77 | 86.49 | 88.41 | 86.77 |
| **59: Sand-Rat** | | | 91.46 | 91.46 | 91.46 | 91.46 | 91.11 | 91.20 |
| 91.11 | 91.33 | 91.20 | 91.20 | 91.24 | 91.31 | 91.28 | 90.76 | 91.15 |
| 91.33 | 90.30 | 91.35 | 90.71 | 90.88 | 90.92 | 91.26 | 90.28 | 90.80 |
| 88.51 | 87.56 | 88.51 | 89.16 | 88.51 | 87.47 | 87.30 | 87.86 | 87.60 |
| 87.54 | 93.48 | 89.57 | 98.83 | 94.31 | 90.33 | 90.14 | 89.18 | 91.43 |
| 93.57 | 94.31 | 94.36 | 94.51 | 94.45 | 93.61 | 91.63 | 88.38 | 91.81 |
| 89.16 | 89.86 | 91.54 | 94.22 | 91.59 | 94.09 | 94.17 | 100.00 | 89.73 |
| 89.60 | 89.47 | 89.43 | 89.54 | 89.58 | 89.84 | 89.80 | 89.50 | 89.37 |
| 89.32 | 89.88 | 88.49 | 88.54 | 89.71 | 89.67 | 89.53 | 90.52 | 87.94 |
| 89.48 | 89.86 | 89.69 | 89.94 | 89.56 | 88.31 | 90.03 | 89.68 | 89.68 |
| 89.97 | 89.51 | 90.06 | 90.01 | 89.64 | 89.48 | 89.60 | 89.77 | 89.82 |
| 90.03 | 90.12 | 89.83 | 89.83 | 90.21 | 89.73 | 89.60 | 89.82 | 89.64 |
| 89.82 | 89.77 | 89.64 | 89.64 | 87.92 | 89.69 | 86.67 | 88.72 | 86.85 |
| **60: Camel** | | | 93.84 | 93.84 | 93.76 | 93.84 | 93.37 | 93.58 |
| 93.50 | 93.71 | 93.41 | 93.45 | 93.62 | 93.47 | 93.54 | 93.02 | 93.28 |
| 93.67 | 92.77 | 93.62 | 92.62 | 93.10 | 93.19 | 93.62 | 92.19 | 92.89 |
| 93.04 | 91.96 | 92.99 | 94.59 | 92.95 | 91.96 | 92.44 | 92.26 | 92.65 |
| 91.97 | 89.21 | 91.09 | 89.86 | 90.74 | 95.17 | 94.89 | 90.00 | 89.80 |
| 89.21 | 90.05 | 90.40 | 89.82 | 90.53 | 89.56 | 93.11 | 92.94 | 93.02 |
| 89.62 | 94.72 | 92.44 | 89.69 | 92.98 | 89.52 | 89.56 | 89.73 | 100.00 |
| 99.61 | 92.05 | 92.05 | 95.75 | 95.84 | 95.97 | 96.06 | 95.58 | 95.32 |
| 95.41 | 96.14 | 93.34 | 93.34 | 95.84 | 96.27 | 94.41 | 92.34 | 89.47 |
| 95.75 | 95.16 | 95.07 | 95.33 | 94.90 | 90.89 | 96.36 | 96.32 | 96.24 |
| 96.28 | 96.02 | 96.33 | 96.33 | 96.23 | 95.66 | 95.07 | 95.14 | 95.31 |
| 94.96 | 94.96 | 94.93 | 94.93 | 94.96 | 95.03 | 94.98 | 95.16 | 94.94 |
| 95.16 | 95.07 | 95.03 | 94.94 | 93.36 | 94.64 | 89.37 | 93.02 | 90.36 |
| **61: Wild-Camel** | | | 93.71 | 93.71 | 93.62 | 93.71 | 93.19 | 93.40 |
| 93.32 | 93.62 | 93.19 | 93.32 | 93.49 | 93.63 | 93.40 | 93.01 | 93.27 |
| 93.53 | 92.68 | 93.58 | 92.44 | 93.23 | 93.32 | 93.54 | 92.05 | 92.84 |
| 92.88 | 92.09 | 92.83 | 94.70 | 92.75 | 92.09 | 92.44 | 92.31 | 92.36 |
| 92.01 | 89.12 | 91.05 | 89.73 | 90.64 | 95.24 | 94.84 | 90.10 | 89.71 |
| 89.12 | 89.96 | 90.35 | 89.72 | 90.49 | 89.43 | 93.01 | 92.81 | 92.93 |
| 89.56 | 95.01 | 92.35 | 89.60 | 92.93 | 89.48 | 89.51 | 89.60 | 99.61 |
| 100.00 | 92.17 | 92.17 | 95.58 | 95.67 | 95.80 | 95.88 | 95.41 | 95.15 |
| 95.23 | 95.97 | 93.47 | 93.47 | 95.62 | 96.14 | 94.54 | 92.21 | 89.42 |
| 95.57 | 95.35 | 95.26 | 95.56 | 95.09 | 90.66 | 96.51 | 96.47 | 96.46 |
| 96.28 | 96.25 | 96.28 | 96.24 | 96.38 | 95.49 | 95.26 | 95.03 | 95.20 |
| 94.94 | 94.98 | 94.84 | 94.84 | 94.98 | 95.22 | 95.26 | 95.44 | 95.22 |
| 95.44 | 95.35 | 95.31 | 95.22 | 93.23 | 94.92 | 89.48 | 92.85 | 90.34 |
| **62: African-Elephant** | | | 92.33 | 92.33 | 92.24 | 92.33 | 92.03 | 92.16 |
| 92.07 | 92.16 | 92.03 | 92.16 | 92.28 | 91.99 | 92.19 | 91.81 | 92.16 |
| 92.37 | 91.07 | 91.97 | 91.41 | 91.89 | 91.84 | 92.10 | 90.80 | 91.29 |
| 90.42 | 89.46 | 90.38 | 91.46 | 90.25 | 89.54 | 89.76 | 89.46 | 89.81 |
| 89.46 | 88.29 | 89.63 | 89.51 | 89.46 | 92.04 | 91.76 | 88.55 | 88.97 |

| | | | | | | | | |
|---|---|---|---|---|---|---|---|---|
| 88.13 | 89.38 | 89.38 | 88.85 | 89.56 | 88.65 | 91.94 | 90.05 | 91.81 |
| 88.62 | 91.86 | 91.45 | 88.73 | 91.68 | 88.64 | 88.99 | 89.47 | 92.05 |
| 92.17 | 100.00 | 99.87 | 91.16 | 91.17 | 91.33 | 91.42 | 91.04 | 90.95 |
| 90.91 | 91.42 | 90.69 | 90.69 | 91.25 | 91.33 | 91.59 | 90.70 | 88.00 |
| 91.32 | 92.17 | 92.04 | 92.33 | 91.99 | 94.94 | 91.82 | 91.78 | 91.76 |
| 91.75 | 91.59 | 91.79 | 91.71 | 91.69 | 91.32 | 91.86 | 91.79 | 91.88 |
| 92.01 | 91.83 | 91.76 | 91.76 | 91.88 | 91.91 | 91.95 | 92.08 | 92.08 |
| 92.04 | 91.99 | 91.86 | 91.99 | 89.89 | 91.95 | 89.46 | 90.20 | 89.62 |
| **63: Indian-Elephant** | | | <u>92.33</u> | 92.33 | 92.24 | 92.33 | 92.03 | 92.16 |
| 92.07 | 92.16 | 92.03 | 92.16 | 92.28 | 91.99 | 92.19 | 91.81 | 92.16 |
| 92.37 | 91.07 | 91.97 | 91.32 | 91.89 | 91.84 | 92.10 | 90.72 | 91.29 |
| 90.42 | 89.46 | 90.38 | 91.55 | 90.25 | 89.54 | 89.76 | 89.46 | 89.77 |
| 89.46 | 88.25 | 89.67 | 89.47 | 89.41 | 92.04 | 91.76 | 88.59 | 88.97 |
| 88.09 | 89.34 | 89.38 | 88.81 | 89.56 | 88.65 | 91.94 | 90.05 | 91.81 |
| 88.62 | 91.86 | 91.45 | 88.69 | 91.68 | 88.60 | 88.94 | 89.43 | 92.05 |
| 92.17 | 99.87 | 100.00 | 91.12 | 91.12 | 91.29 | 91.38 | 90.99 | 90.91 |
| 90.86 | 91.38 | 90.69 | 90.69 | 91.21 | 91.33 | 91.54 | 90.70 | 88.00 |
| 91.23 | 92.17 | 92.04 | 92.33 | 91.99 | 94.86 | 91.82 | 91.78 | 91.76 |
| 91.75 | 91.59 | 91.79 | 91.71 | 91.69 | 91.23 | 91.86 | 91.79 | 91.88 |
| 92.01 | 91.83 | 91.76 | 91.76 | 91.88 | 91.91 | 91.95 | 92.08 | 92.08 |
| 92.04 | 91.99 | 91.86 | 91.99 | 89.89 | 91.95 | 89.46 | 90.16 | 89.62 |
| **64: Cow** | | | <u>93.04</u> | 93.04 | 92.96 | 93.04 | 92.65 | 92.83 |
| 92.74 | 92.91 | 92.44 | 92.61 | 92.82 | 92.72 | 92.73 | 92.13 | 92.52 |
| 92.87 | 91.74 | 92.80 | 91.60 | 92.37 | 92.46 | 92.67 | 91.25 | 92.22 |
| 92.15 | 90.91 | 92.15 | 93.63 | 91.94 | 90.96 | 91.17 | 90.91 | 91.29 |
| 90.88 | 88.67 | 90.27 | 89.75 | 90.20 | 94.50 | 94.49 | 89.44 | 89.39 |
| 88.63 | 89.70 | 89.87 | 89.61 | 90.05 | 89.11 | 92.39 | 92.59 | 92.31 |
| 89.07 | 94.10 | 91.73 | 89.45 | 92.31 | 89.14 | 89.19 | 89.54 | 95.75 |
| 95.58 | 91.16 | 91.12 | 100.00 | 99.31 | 99.70 | 99.14 | 98.20 | 98.41 |
| 98.03 | 99.14 | 92.34 | 92.30 | 98.76 | 95.91 | 93.50 | 91.76 | 88.81 |
| 95.04 | 94.15 | 94.06 | 94.36 | 93.97 | 90.10 | 95.53 | 95.53 | 95.49 |
| 95.58 | 95.31 | 95.71 | 95.54 | 95.49 | 95.04 | 94.19 | 94.32 | 94.50 |
| 94.45 | 94.32 | 94.28 | 94.28 | 94.32 | 93.97 | 94.02 | 94.10 | 93.93 |
| 94.10 | 94.06 | 93.93 | 93.93 | 92.55 | 93.89 | 88.74 | 92.33 | 89.41 |
| **65: Water-Buffalo** | | | <u>93.05</u> | 93.05 | 92.96 | 93.05 | 92.66 | 92.83 |
| 92.74 | 92.92 | 92.44 | 92.66 | 92.87 | 92.76 | 92.78 | 92.17 | 92.44 |
| 92.87 | 91.75 | 92.80 | 91.65 | 92.29 | 92.37 | 92.68 | 91.30 | 92.35 |
| 92.19 | 90.92 | 92.19 | 93.67 | 91.94 | 90.96 | 91.22 | 90.87 | 91.38 |
| 90.79 | 88.76 | 90.40 | 89.80 | 90.21 | 94.42 | 94.40 | 89.44 | 89.39 |
| 88.73 | 89.74 | 89.92 | 89.66 | 90.10 | 89.15 | 92.35 | 92.68 | 92.31 |
| 89.08 | 94.06 | 91.69 | 89.45 | 92.27 | 89.24 | 89.28 | 89.58 | 95.84 |
| 95.67 | 91.17 | 91.12 | 99.31 | 100.00 | 99.53 | 99.23 | 98.33 | 98.50 |
| 98.16 | 99.31 | 92.39 | 92.34 | 98.89 | 96.13 | 93.49 | 91.81 | 89.03 |
| 95.09 | 94.06 | 93.97 | 94.28 | 93.89 | 89.93 | 95.58 | 95.49 | 95.45 |
| 95.58 | 95.27 | 95.75 | 95.58 | 95.45 | 95.09 | 94.06 | 94.20 | 94.37 |
| 94.37 | 94.24 | 94.27 | 94.27 | 94.24 | 93.97 | 94.02 | 94.10 | 93.93 |
| 94.10 | 94.06 | 93.93 | 93.93 | 92.55 | 93.89 | 88.70 | 92.33 | 89.38 |
| **66: Bos-Banteng** | | | <u>93.34</u> | 93.34 | 93.26 | 93.34 | 92.96 | 93.13 |
| 93.04 | 93.22 | 92.74 | 92.91 | 93.12 | 93.02 | 93.04 | 92.43 | 92.74 |
| 93.17 | 92.07 | 93.10 | 91.90 | 92.59 | 92.67 | 92.97 | 91.55 | 92.52 |

| | | | | | | | | |
|---|---|---|---|---|---|---|---|---|
| 92.41 | 91.17 | 92.41 | 93.93 | 92.20 | 91.22 | 91.39 | 91.17 | 91.55 |
| 91.05 | 88.97 | 90.57 | 90.06 | 90.46 | 94.72 | 94.71 | 89.74 | 89.69 |
| 88.94 | 90.00 | 90.17 | 89.92 | 90.35 | 89.41 | 92.65 | 92.89 | 92.61 |
| 89.38 | 94.32 | 92.03 | 89.67 | 92.57 | 89.45 | 89.49 | 89.84 | 95.97 |
| 95.80 | 91.33 | 91.29 | 99.70 | 99.53 | 100.00 | 99.31 | 98.41 | 98.63 |
| 98.24 | 99.40 | 92.64 | 92.60 | 99.01 | 96.17 | 93.80 | 92.07 | 89.07 |
| 95.34 | 94.32 | 94.23 | 94.54 | 94.15 | 90.23 | 95.84 | 95.84 | 95.79 |
| 95.84 | 95.62 | 96.01 | 95.84 | 95.79 | 95.34 | 94.41 | 94.54 | 94.71 |
| 94.67 | 94.54 | 94.49 | 94.49 | 94.54 | 94.19 | 94.23 | 94.32 | 94.15 |
| 94.32 | 94.28 | 94.15 | 94.15 | 92.76 | 94.10 | 88.78 | 92.59 | 89.59 |
| **67: Sheep** | | | 93.26 | 93.26 | 93.17 | 93.26 | 92.74 | 92.96 |
| 92.87 | 93.13 | 92.65 | 92.83 | 93.04 | 92.93 | 92.95 | 92.43 | 92.78 |
| 93.09 | 91.98 | 93.02 | 91.94 | 92.54 | 92.63 | 92.89 | 91.64 | 92.56 |
| 92.28 | 91.04 | 92.28 | 93.76 | 92.07 | 91.13 | 91.39 | 91.09 | 91.46 |
| 90.92 | 88.88 | 90.27 | 90.01 | 90.38 | 94.59 | 94.58 | 89.57 | 89.56 |
| 88.81 | 89.91 | 90.17 | 89.83 | 90.35 | 89.41 | 92.61 | 92.72 | 92.52 |
| 89.29 | 94.19 | 91.86 | 89.54 | 92.61 | 89.36 | 89.41 | 89.80 | 96.06 |
| 95.88 | 91.42 | 91.38 | 99.14 | 99.23 | 99.31 | 100.00 | 98.50 | 98.58 |
| 98.33 | 99.91 | 92.51 | 92.47 | 99.36 | 96.21 | 93.63 | 91.85 | 88.94 |
| 95.26 | 94.28 | 94.19 | 94.50 | 94.10 | 90.23 | 95.71 | 95.49 | 95.44 |
| 95.71 | 95.27 | 95.80 | 95.71 | 95.45 | 95.26 | 94.23 | 94.37 | 94.54 |
| 94.50 | 94.41 | 94.32 | 94.32 | 94.41 | 94.15 | 94.19 | 94.28 | 94.10 |
| 94.28 | 94.23 | 94.10 | 94.10 | 92.72 | 93.97 | 88.83 | 92.37 | 89.59 |
| **68: Deer** | | | 92.87 | 92.87 | 92.79 | 92.87 | 92.40 | 92.57 |
| 92.40 | 92.74 | 92.35 | 92.48 | 92.69 | 92.63 | 92.61 | 92.17 | 92.53 |
| 92.74 | 91.65 | 92.72 | 91.43 | 92.33 | 92.42 | 92.63 | 91.26 | 92.31 |
| 91.85 | 90.61 | 91.85 | 93.45 | 91.68 | 90.70 | 90.83 | 90.57 | 91.07 |
| 90.49 | 88.59 | 90.10 | 89.63 | 89.99 | 94.37 | 94.40 | 89.18 | 89.26 |
| 88.51 | 89.66 | 89.87 | 89.57 | 90.01 | 89.07 | 92.18 | 92.12 | 92.18 |
| 88.82 | 93.93 | 91.56 | 89.15 | 92.10 | 89.06 | 89.06 | 89.50 | 95.58 |
| 95.41 | 91.04 | 90.99 | 98.20 | 98.33 | 98.41 | 98.50 | 100.00 | 97.90 |
| 99.27 | 98.50 | 92.43 | 92.39 | 98.16 | 95.70 | 93.36 | 91.64 | 88.73 |
| 94.79 | 93.84 | 93.80 | 94.06 | 93.67 | 89.84 | 95.45 | 95.06 | 95.01 |
| 95.36 | 94.88 | 95.45 | 95.45 | 95.02 | 94.79 | 93.89 | 94.02 | 94.20 |
| 94.24 | 94.24 | 93.93 | 93.93 | 94.20 | 93.93 | 93.97 | 94.06 | 93.93 |
| 94.02 | 93.97 | 93.84 | 93.84 | 92.46 | 93.63 | 88.83 | 91.99 | 89.55 |
| **69: Musk-Deer** | | | 92.53 | 92.53 | 92.44 | 92.53 | 92.14 | 92.31 |
| 92.22 | 92.44 | 91.97 | 92.14 | 92.35 | 92.24 | 92.26 | 91.87 | 92.10 |
| 92.40 | 91.23 | 92.37 | 91.17 | 91.94 | 92.03 | 92.20 | 90.82 | 91.96 |
| 91.85 | 90.57 | 91.85 | 93.11 | 91.68 | 90.79 | 90.87 | 90.57 | 91.12 |
| 90.58 | 88.37 | 89.70 | 89.45 | 89.99 | 93.98 | 93.97 | 89.18 | 88.96 |
| 88.29 | 89.49 | 89.61 | 89.27 | 89.88 | 88.89 | 92.01 | 92.20 | 91.97 |
| 88.64 | 93.63 | 91.43 | 89.06 | 91.92 | 88.85 | 88.85 | 89.37 | 95.32 |
| 95.15 | 90.95 | 90.91 | 98.41 | 98.50 | 98.63 | 98.58 | 97.90 | 100.00 |
| 97.73 | 98.58 | 92.00 | 91.96 | 98.20 | 95.65 | 93.19 | 91.51 | 88.69 |
| 94.70 | 93.50 | 93.41 | 93.72 | 93.32 | 89.50 | 95.10 | 94.93 | 94.88 |
| 95.06 | 94.71 | 95.10 | 95.10 | 94.89 | 94.79 | 93.58 | 93.76 | 93.89 |
| 93.89 | 93.81 | 93.80 | 93.80 | 93.81 | 93.45 | 93.54 | 93.63 | 93.54 |
| 93.58 | 93.54 | 93.41 | 93.41 | 92.03 | 93.41 | 88.39 | 91.90 | 89.12 |

| | | | | | | | | |
|---|---|---|---|---|---|---|---|---|
| **70: Red-Deer** | | | 92.61 | 92.61 | 92.53 | 92.61 | 92.18 | 92.31 |
| 92.14 | 92.48 | 92.10 | 92.22 | 92.43 | 92.37 | 92.35 | 92.04 | 92.27 |
| 92.48 | 91.42 | 92.46 | 91.30 | 92.07 | 92.16 | 92.37 | 91.13 | 92.05 |
| 91.67 | 90.61 | 91.67 | 93.28 | 91.51 | 90.83 | 90.83 | 90.61 | 91.03 |
| 90.32 | 88.41 | 89.88 | 89.37 | 89.86 | 94.11 | 94.15 | 89.01 | 89.13 |
| 88.33 | 89.57 | 89.79 | 89.31 | 89.92 | 88.98 | 92.01 | 91.99 | 92.01 |
| 88.64 | 93.71 | 91.43 | 89.02 | 91.92 | 88.93 | 88.93 | 89.32 | 95.41 |
| 95.23 | 90.91 | 90.86 | 98.03 | 98.16 | 98.24 | 98.33 | 99.27 | 97.73 |
| 100.00 | 98.33 | 92.13 | 92.08 | 97.98 | 95.52 | 93.19 | 91.55 | 88.60 |
| 94.57 | 93.63 | 93.58 | 93.85 | 93.45 | 89.76 | 95.23 | 94.84 | 94.80 |
| 95.15 | 94.67 | 95.23 | 95.19 | 94.80 | 94.57 | 93.67 | 93.81 | 93.98 |
| 93.98 | 93.98 | 93.72 | 93.72 | 93.94 | 93.71 | 93.76 | 93.84 | 93.67 |
| 93.80 | 93.76 | 93.63 | 93.63 | 92.24 | 93.41 | 88.57 | 91.86 | 89.42 |

| | | | | | | | | |
|---|---|---|---|---|---|---|---|---|
| **71: Goat** | | | 93.34 | 93.34 | 93.26 | 93.34 | 92.83 | 93.04 |
| 92.96 | 93.22 | 92.74 | 92.91 | 93.12 | 93.02 | 93.04 | 92.52 | 92.78 |
| 93.17 | 92.07 | 93.10 | 92.03 | 92.63 | 92.72 | 92.97 | 91.73 | 92.65 |
| 92.32 | 91.09 | 92.32 | 93.84 | 92.11 | 91.17 | 91.43 | 91.13 | 91.50 |
| 90.96 | 88.97 | 90.35 | 90.10 | 90.46 | 94.63 | 94.62 | 89.66 | 89.65 |
| 88.89 | 90.00 | 90.26 | 89.92 | 90.44 | 89.50 | 92.70 | 92.80 | 92.61 |
| 89.38 | 94.23 | 91.94 | 89.62 | 92.70 | 89.45 | 89.49 | 89.88 | 96.14 |
| 95.97 | 91.42 | 91.38 | 99.14 | 99.31 | 99.40 | 99.91 | 98.50 | 98.58 |
| 98.33 | 100.00 | 92.60 | 92.56 | 99.40 | 96.30 | 93.71 | 91.94 | 89.03 |
| 95.34 | 94.32 | 94.23 | 94.54 | 94.15 | 90.23 | 95.79 | 95.58 | 95.53 |
| 95.79 | 95.36 | 95.88 | 95.80 | 95.53 | 95.34 | 94.28 | 94.41 | 94.58 |
| 94.54 | 94.45 | 94.36 | 94.36 | 94.45 | 94.19 | 94.23 | 94.32 | 94.15 |
| 94.32 | 94.28 | 94.15 | 94.15 | 92.76 | 94.02 | 88.83 | 92.46 | 89.59 |

| | | | | | | | | |
|---|---|---|---|---|---|---|---|---|
| **72: Zebra** | | | 92.44 | 92.44 | 92.35 | 92.44 | 92.01 | 92.18 |
| 92.01 | 92.27 | 92.05 | 91.96 | 92.17 | 92.42 | 92.09 | 91.39 | 91.75 |
| 92.22 | 91.28 | 92.46 | 91.36 | 92.07 | 92.16 | 92.38 | 91.01 | 91.79 |
| 91.55 | 90.17 | 91.55 | 93.34 | 91.42 | 90.39 | 90.52 | 90.30 | 91.08 |
| 90.79 | 87.75 | 89.52 | 88.62 | 88.77 | 93.52 | 93.68 | 88.91 | 88.82 |
| 87.80 | 88.87 | 88.78 | 88.39 | 89.05 | 88.10 | 91.88 | 91.82 | 91.83 |
| 88.71 | 93.25 | 90.65 | 88.32 | 91.70 | 88.27 | 88.36 | 88.49 | 93.34 |
| 93.47 | 90.69 | 90.69 | 92.34 | 92.39 | 92.64 | 92.51 | 92.43 | 92.00 |
| 92.13 | 92.60 | 100.00 | 99.91 | 92.43 | 92.68 | 94.58 | 91.24 | 88.50 |
| 92.89 | 93.77 | 93.73 | 93.99 | 93.60 | 89.84 | 93.07 | 92.81 | 92.77 |
| 92.77 | 92.59 | 93.04 | 93.12 | 92.77 | 92.76 | 93.60 | 93.60 | 93.69 |
| 93.34 | 93.43 | 93.85 | 93.85 | 93.43 | 93.68 | 93.77 | 93.82 | 93.68 |
| 93.82 | 93.86 | 93.73 | 93.73 | 91.94 | 93.86 | 89.04 | 91.60 | 89.54 |

| | | | | | | | | |
|---|---|---|---|---|---|---|---|---|
| **73: Horse** | | | 92.40 | 92.40 | 92.31 | 92.40 | 91.96 | 92.14 |
| 92.05 | 92.22 | 92.01 | 91.92 | 92.13 | 92.38 | 92.04 | 91.35 | 91.70 |
| 92.18 | 91.23 | 92.42 | 91.40 | 92.03 | 92.12 | 92.33 | 90.96 | 91.74 |
| 91.55 | 90.17 | 91.55 | 93.34 | 91.42 | 90.39 | 90.52 | 90.30 | 91.08 |
| 90.79 | 87.79 | 89.56 | 88.67 | 88.81 | 93.47 | 93.64 | 88.86 | 88.86 |
| 87.85 | 88.91 | 88.83 | 88.44 | 89.09 | 88.15 | 91.88 | 91.77 | 91.83 |
| 88.67 | 93.21 | 90.70 | 88.36 | 91.70 | 88.32 | 88.40 | 88.54 | 93.34 |
| 93.47 | 90.69 | 90.69 | 92.30 | 92.34 | 92.60 | 92.47 | 92.39 | 91.96 |
| 92.08 | 92.56 | 99.91 | 100.00 | 92.39 | 92.64 | 94.63 | 91.20 | 88.45 |
| 92.85 | 93.82 | 93.77 | 93.95 | 93.55 | 89.80 | 93.07 | 92.81 | 92.77 |
| 92.77 | 92.59 | 93.04 | 93.12 | 92.77 | 92.72 | 93.55 | 93.56 | 93.64 |

| | | | | | | | | |
|---|---|---|---|---|---|---|---|---|
| 93.30 | 93.38 | 93.81 | 93.81 | 93.38 | 93.64 | 93.73 | 93.77 | 93.64 |
| 93.77 | 93.82 | 93.68 | 93.68 | 91.90 | 93.82 | 89.00 | 91.64 | 89.49 |

| | | | | | | | | |
|---|---|---|---|---|---|---|---|---|
| **74: Oryx** | | | 93.09 | 93.09 | 93.00 | 93.09 | 92.66 | 92.87 |
| 92.79 | 92.96 | 92.61 | 92.66 | 92.87 | 92.76 | 92.78 | 92.26 | 92.53 |
| 92.92 | 91.84 | 92.85 | 91.86 | 92.37 | 92.46 | 92.72 | 91.52 | 92.44 |
| 92.06 | 90.83 | 92.06 | 93.76 | 91.81 | 90.87 | 91.18 | 90.87 | 91.29 |
| 90.75 | 88.89 | 90.10 | 89.93 | 90.29 | 94.42 | 94.36 | 89.44 | 89.52 |
| 88.81 | 89.83 | 90.05 | 89.75 | 90.18 | 89.28 | 92.61 | 92.51 | 92.48 |
| 89.21 | 94.02 | 91.77 | 89.41 | 92.57 | 89.24 | 89.28 | 89.71 | 95.84 |
| 95.62 | 91.25 | 91.21 | 98.76 | 98.89 | 99.01 | 99.36 | 98.16 | 98.20 |
| 97.98 | 99.40 | 92.43 | 92.39 | 100.00 | 96.00 | 93.62 | 91.77 | 89.08 |
| 95.05 | 94.02 | 93.93 | 94.24 | 93.84 | 90.10 | 95.41 | 95.32 | 95.27 |
| 95.41 | 95.10 | 95.58 | 95.49 | 95.28 | 95.05 | 94.02 | 94.20 | 94.37 |
| 94.33 | 94.24 | 94.10 | 94.10 | 94.24 | 93.97 | 94.02 | 94.10 | 93.93 |
| 94.02 | 93.97 | 93.84 | 93.84 | 92.46 | 93.80 | 88.66 | 92.20 | 89.38 |

| | | | | | | | | |
|---|---|---|---|---|---|---|---|---|
| **75: Hippopotamus** | | | 93.08 | 93.08 | 92.99 | 93.08 | 92.60 | 92.73 |
| 92.82 | 92.99 | 92.60 | 92.73 | 92.90 | 92.89 | 92.81 | 92.34 | 92.69 |
| 92.95 | 91.98 | 92.97 | 91.76 | 92.58 | 92.67 | 92.88 | 91.55 | 92.73 |
| 92.62 | 91.52 | 92.62 | 94.02 | 92.50 | 91.52 | 91.69 | 91.61 | 92.15 |
| 91.64 | 88.88 | 90.52 | 89.84 | 90.46 | 94.37 | 94.49 | 89.43 | 89.47 |
| 88.85 | 90.04 | 90.39 | 89.61 | 90.56 | 89.45 | 92.86 | 93.10 | 92.73 |
| 89.18 | 94.15 | 91.85 | 89.45 | 92.78 | 89.45 | 89.27 | 89.67 | 96.27 |
| 96.14 | 91.33 | 91.33 | 95.91 | 96.13 | 96.17 | 96.21 | 95.70 | 95.65 |
| 95.52 | 96.30 | 92.68 | 92.64 | 96.00 | 100.00 | 93.97 | 91.71 | 88.81 |
| 95.39 | 94.32 | 94.28 | 94.54 | 94.28 | 90.27 | 96.53 | 96.36 | 96.31 |
| 96.66 | 96.10 | 96.79 | 96.79 | 96.31 | 95.39 | 94.15 | 94.19 | 94.36 |
| 94.28 | 94.19 | 94.27 | 94.27 | 94.23 | 94.10 | 94.23 | 94.28 | 94.10 |
| 94.32 | 94.23 | 94.15 | 94.19 | 92.67 | 93.84 | 88.91 | 92.63 | 89.63 |

| | | | | | | | | |
|---|---|---|---|---|---|---|---|---|
| **76: Rhinoceros** | | | 93.16 | 93.25 | 93.12 | 93.16 | 92.77 | 92.90 |
| 92.86 | 93.16 | 92.81 | 92.73 | 92.94 | 93.19 | 92.94 | 92.20 | 92.68 |
| 92.99 | 92.20 | 93.18 | 92.15 | 92.75 | 92.84 | 93.09 | 91.71 | 92.16 |
| 92.62 | 91.39 | 92.62 | 94.14 | 92.50 | 91.61 | 91.82 | 91.43 | 91.93 |
| 91.82 | 88.84 | 90.38 | 89.66 | 90.06 | 94.45 | 94.70 | 89.47 | 89.43 |
| 88.75 | 89.86 | 89.94 | 89.35 | 90.21 | 89.14 | 92.55 | 92.71 | 92.55 |
| 89.23 | 94.40 | 91.89 | 89.31 | 92.42 | 89.23 | 89.18 | 89.53 | 94.41 |
| 94.54 | 91.59 | 91.54 | 93.50 | 93.49 | 93.80 | 93.63 | 93.36 | 93.19 |
| 93.19 | 93.71 | 94.58 | 94.63 | 93.62 | 93.97 | 100.00 | 91.66 | 88.46 |
| 93.96 | 94.75 | 94.66 | 94.88 | 94.75 | 90.74 | 94.36 | 94.19 | 94.14 |
| 94.02 | 93.97 | 94.36 | 94.41 | 94.19 | 93.87 | 94.58 | 94.58 | 94.67 |
| 94.32 | 94.32 | 94.66 | 94.66 | 94.41 | 94.62 | 94.62 | 94.75 | 94.66 |
| 94.79 | 94.75 | 94.62 | 94.71 | 93.06 | 94.58 | 89.17 | 92.54 | 89.98 |

| | | | | | | | | |
|---|---|---|---|---|---|---|---|---|
| **77: Rabbit** | | | 94.40 | 94.44 | 94.36 | 94.49 | 94.05 | 94.27 |
| 94.01 | 94.23 | 93.93 | 94.05 | 94.22 | 93.79 | 94.14 | 93.36 | 93.71 |
| 94.18 | 92.72 | 93.87 | 92.83 | 93.22 | 93.35 | 93.74 | 92.62 | 93.45 |
| 90.72 | 89.19 | 90.72 | 91.11 | 90.60 | 89.32 | 89.41 | 89.28 | 89.90 |
| 89.79 | 89.48 | 91.07 | 90.74 | 91.10 | 92.94 | 92.97 | 90.72 | 90.72 |
| 89.58 | 91.07 | 91.32 | 90.64 | 91.54 | 90.35 | 93.84 | 90.58 | 93.67 |
| 90.65 | 92.46 | 93.35 | 90.43 | 93.67 | 90.22 | 90.21 | 90.52 | 92.34 |
| 92.21 | 90.70 | 90.70 | 91.76 | 91.81 | 92.07 | 91.85 | 91.64 | 91.51 |
| 91.55 | 91.94 | 91.24 | 91.20 | 91.77 | 91.71 | 91.66 | 100.00 | 91.75 |

| | | | | | | | | |
|---|---|---|---|---|---|---|---|---|
| 91.66 | 92.46 | 92.25 | 92.59 | 92.29 | 89.33 | 91.55 | 91.55 | 91.50 |
| 91.61 | 91.33 | 91.65 | 91.65 | 91.47 | 91.66 | 92.33 | 92.56 | 92.60 |
| 92.86 | 92.82 | 92.75 | 92.75 | 92.82 | 92.59 | 92.55 | 92.59 | 92.46 |
| 92.68 | 92.64 | 92.51 | 92.59 | 90.89 | 92.51 | 89.01 | 90.63 | 89.51 |
| **78: Pika-hare** | | | <u>90.93</u> | 90.93 | 90.85 | 90.89 | 90.50 | 90.67 |
| 90.50 | 90.80 | 90.37 | 90.72 | 90.84 | 90.68 | 90.75 | 89.89 | 90.28 |
| 90.67 | 89.37 | 90.71 | 89.78 | 90.06 | 90.02 | 90.36 | 89.83 | 90.10 |
| 87.60 | 86.54 | 87.64 | 89.02 | 87.60 | 86.54 | 87.02 | 86.46 | 87.08 |
| 86.75 | 87.02 | 88.20 | 88.33 | 87.78 | 90.03 | 89.81 | 87.93 | 88.01 |
| 87.07 | 87.67 | 88.23 | 87.93 | 88.54 | 87.60 | 90.11 | 87.80 | 90.16 |
| 87.55 | 89.81 | 90.05 | 87.38 | 90.07 | 87.55 | 87.37 | 87.94 | 89.47 |
| 89.42 | 88.00 | 88.00 | 88.81 | 89.03 | 89.07 | 88.94 | 88.73 | 88.69 |
| 88.60 | 89.03 | 88.50 | 88.45 | 89.08 | 88.81 | 88.46 | 91.75 | 100.00 |
| 88.89 | 89.77 | 89.68 | 89.93 | 89.42 | 86.98 | 88.85 | 88.76 | 88.66 |
| 88.82 | 88.58 | 88.99 | 88.86 | 88.68 | 88.85 | 89.60 | 89.69 | 89.73 |
| 89.90 | 89.90 | 89.72 | 89.72 | 90.03 | 89.77 | 89.73 | 89.86 | 89.64 |
| 89.90 | 89.86 | 89.81 | 89.90 | 87.76 | 89.73 | 86.52 | 87.91 | 86.87 |
| **79: Pig** | | | <u>93.16</u> | 93.16 | 93.07 | 93.16 | 92.81 | 92.94 |
| 92.90 | 93.03 | 92.77 | 92.73 | 92.89 | 92.75 | 92.81 | 92.16 | 92.51 |
| 92.94 | 91.87 | 92.79 | 92.15 | 92.32 | 92.40 | 92.79 | 92.06 | 92.20 |
| 92.31 | 91.12 | 92.31 | 93.75 | 92.19 | 91.25 | 91.38 | 91.34 | 91.67 |
| 91.20 | 88.74 | 90.21 | 89.74 | 89.93 | 94.28 | 94.48 | 89.82 | 89.73 |
| 88.66 | 89.68 | 89.68 | 89.56 | 89.99 | 88.97 | 92.47 | 92.71 | 92.47 |
| 89.35 | 93.97 | 92.10 | 89.31 | 92.42 | 89.27 | 89.18 | 89.48 | 95.75 |
| 95.57 | 91.32 | 91.23 | 95.04 | 95.09 | 95.34 | 95.26 | 94.79 | 94.70 |
| 94.57 | 95.34 | 92.89 | 92.85 | 95.05 | 95.39 | 93.96 | 91.66 | 88.89 |
| 100.00 | 93.97 | 93.88 | 94.19 | 94.05 | 90.74 | 95.40 | 95.10 | 95.01 |
| 95.36 | 94.79 | 95.44 | 95.57 | 94.97 | 99.66 | 93.88 | 94.10 | 94.19 |
| 94.10 | 94.19 | 94.39 | 94.39 | 94.23 | 94.14 | 94.14 | 94.23 | 94.14 |
| 94.31 | 94.23 | 94.10 | 94.10 | 92.67 | 93.97 | 88.55 | 92.62 | 89.32 |
| **80: Seal** | | | <u>93.97</u> | 94.10 | 93.92 | 93.97 | 93.49 | 93.66 |
| 93.66 | 93.88 | 93.40 | 93.53 | 93.75 | 93.76 | 93.66 | 92.80 | 93.32 |
| 93.75 | 92.73 | 93.66 | 92.79 | 93.19 | 93.23 | 93.53 | 92.35 | 93.10 |
| 92.88 | 91.96 | 92.79 | 95.09 | 92.58 | 92.09 | 92.17 | 91.92 | 91.85 |
| 91.83 | 89.73 | 91.31 | 90.03 | 90.68 | 98.10 | 97.61 | 90.14 | 89.75 |
| 89.51 | 90.39 | 90.48 | 90.16 | 90.66 | 89.73 | 93.14 | 92.98 | 93.10 |
| 89.64 | 97.93 | 92.66 | 90.03 | 93.14 | 89.95 | 90.03 | 89.86 | 95.16 |
| 95.35 | 92.17 | 92.17 | 94.15 | 94.06 | 94.32 | 94.28 | 93.84 | 93.50 |
| 93.63 | 94.32 | 93.77 | 93.82 | 94.02 | 94.32 | 94.75 | 92.46 | 89.77 |
| 93.97 | 100.00 | 99.61 | 99.78 | 98.92 | 91.00 | 94.61 | 94.48 | 94.52 |
| 94.29 | 94.30 | 94.38 | 94.38 | 94.48 | 93.97 | 97.93 | 97.75 | 97.84 |
| 97.79 | 97.84 | 97.44 | 97.44 | 97.84 | 97.24 | 97.20 | 97.37 | 97.29 |
| 97.46 | 97.46 | 97.33 | 97.33 | 95.10 | 97.59 | 89.74 | 93.54 | 90.69 |
| **81: Monk-seal** | | | <u>93.84</u> | 93.97 | 93.79 | 93.84 | 93.36 | 93.53 |
| 93.49 | 93.75 | 93.27 | 93.40 | 93.62 | 93.63 | 93.53 | 92.66 | 93.19 |
| 93.62 | 92.58 | 93.53 | 92.66 | 93.06 | 93.10 | 93.40 | 92.13 | 92.97 |
| 92.79 | 91.96 | 92.70 | 95.04 | 92.58 | 92.09 | 92.17 | 91.92 | 91.85 |
| 91.83 | 89.56 | 91.09 | 89.86 | 90.51 | 97.92 | 97.44 | 89.92 | 89.54 |
| 89.34 | 90.13 | 90.30 | 89.98 | 90.48 | 89.56 | 92.97 | 92.98 | 92.88 |
| 89.51 | 97.76 | 92.44 | 89.86 | 92.97 | 89.78 | 89.86 | 89.69 | 95.07 |

| | | | | | | | | |
|---|---|---|---|---|---|---|---|---|
| 95.26 | 92.04 | 92.04 | 94.06 | 93.97 | 94.23 | 94.19 | 93.80 | 93.41 |
| 93.58 | 94.23 | 93.73 | 93.77 | 93.93 | 94.28 | 94.66 | 92.25 | 89.68 |
| 93.88 | 99.61 | 100.00 | 99.65 | 98.79 | 90.87 | 94.53 | 94.40 | 94.43 |
| 94.20 | 94.26 | 94.29 | 94.29 | 94.40 | 93.88 | 97.80 | 97.62 | 97.71 |
| 97.62 | 97.66 | 97.31 | 97.31 | 97.66 | 97.29 | 97.24 | 97.42 | 97.33 |
| 97.50 | 97.50 | 97.37 | 97.37 | 95.14 | 97.50 | 89.57 | 93.32 | 90.60 |
| **82: Grey-seal** | | | 94.14 | 94.27 | 94.10 | 94.14 | 93.67 | 93.84 |
| 93.75 | 94.06 | 93.58 | 93.71 | 93.92 | 93.94 | 93.84 | 92.93 | 93.49 |
| 93.93 | 92.92 | 93.84 | 92.88 | 93.36 | 93.41 | 93.71 | 92.40 | 93.27 |
| 93.05 | 92.21 | 92.96 | 95.25 | 92.75 | 92.30 | 92.38 | 92.12 | 92.06 |
| 91.92 | 89.77 | 91.27 | 90.11 | 90.76 | 98.27 | 97.79 | 90.23 | 89.74 |
| 89.55 | 90.38 | 90.56 | 90.20 | 90.74 | 89.77 | 93.19 | 93.20 | 93.15 |
| 89.73 | 98.14 | 92.66 | 90.03 | 93.19 | 89.99 | 90.08 | 89.94 | 95.33 |
| 95.56 | 92.33 | 92.33 | 94.36 | 94.28 | 94.54 | 94.50 | 94.06 | 93.72 |
| 93.85 | 94.54 | 93.99 | 93.95 | 94.24 | 94.54 | 94.88 | 92.59 | 89.93 |
| 94.19 | 99.78 | 99.65 | 100.00 | 99.14 | 91.22 | 94.82 | 94.69 | 94.65 |
| 94.46 | 94.43 | 94.55 | 94.55 | 94.69 | 94.19 | 98.19 | 97.97 | 98.05 |
| 97.97 | 98.01 | 97.66 | 97.66 | 98.01 | 97.50 | 97.45 | 97.63 | 97.54 |
| 97.71 | 97.71 | 97.58 | 97.58 | 95.31 | 97.84 | 89.87 | 93.58 | 90.78 |
| **83: Sealion** | | 93.71 | 93.84 | 93.66 | 93.71 | 93.23 | 93.40 | 93.32 |
| 93.62 | 93.19 | 93.36 | 93.49 | 93.55 | 93.40 | 92.58 | 93.14 | 93.53 |
| 92.54 | 93.45 | 92.70 | 92.97 | 93.10 | 93.36 | 92.18 | 92.97 | 92.62 |
| 91.61 | 92.57 | 94.78 | 92.45 | 91.92 | 91.92 | 91.57 | 91.72 | 91.79 |
| 89.25 | 90.96 | 89.73 | 90.38 | 97.79 | 97.35 | 89.83 | 89.49 | 89.04 |
| 90.04 | 90.17 | 89.77 | 90.31 | 89.43 | 92.84 | 92.89 | 92.88 | 89.56 |
| 97.63 | 92.22 | 89.56 | 92.80 | 89.52 | 89.55 | 89.56 | 94.90 | 95.09 |
| 91.99 | 91.99 | 93.97 | 93.89 | 94.15 | 94.10 | 93.67 | 93.32 | 93.45 |
| 94.15 | 93.60 | 93.55 | 93.84 | 94.28 | 94.75 | 92.29 | 89.42 | 94.05 |
| 98.92 | 98.79 | 99.14 | 100.00 | 90.87 | 94.61 | 94.31 | 94.34 | 94.12 |
| 94.13 | 94.38 | 94.38 | 94.31 | 93.97 | 97.85 | 97.66 | 97.75 | 97.49 |
| 97.53 | 97.31 | 97.31 | 97.58 | 97.07 | 97.03 | 97.20 | 97.11 | 97.29 |
| 97.29 | 97.16 | 97.16 | 94.97 | 97.46 | 89.31 | 93.32 | 90.21 | |
| **84: Manatee** | | 90.96 | 90.96 | 90.87 | 90.96 | 90.66 | 90.87 | 90.74 |
| 90.83 | 90.61 | 90.74 | 90.87 | 91.04 | 90.78 | 90.39 | 90.87 | 90.96 |
| 89.55 | 90.66 | 90.25 | 90.48 | 90.53 | 90.61 | 89.86 | 90.22 | 89.54 |
| 88.64 | 89.50 | 90.48 | 89.64 | 88.73 | 88.94 | 88.86 | 88.73 | 88.76 |
| 87.19 | 88.93 | 88.44 | 88.57 | 91.11 | 90.75 | 88.10 | 87.73 | 86.97 |
| 88.28 | 88.19 | 87.88 | 88.37 | 87.45 | 90.66 | 89.14 | 90.74 | 87.51 |
| 90.66 | 90.21 | 87.79 | 90.35 | 87.40 | 87.75 | 88.31 | 90.89 | 90.66 |
| 94.94 | 94.86 | 90.10 | 89.93 | 90.23 | 90.23 | 89.84 | 89.50 | 89.76 |
| 90.23 | 89.84 | 89.80 | 90.10 | 90.27 | 90.74 | 89.33 | 86.98 | 90.74 |
| 91.00 | 90.87 | 91.22 | 90.87 | 100.00 | 90.83 | 90.78 | 90.83 | 90.98 |
| 90.70 | 91.06 | 90.89 | 90.70 | 90.61 | 90.57 | 90.76 | 90.85 | 90.80 |
| 90.85 | 90.61 | 90.61 | 90.93 | 90.66 | 90.70 | 90.74 | 90.74 | 90.70 |
| 90.66 | 90.61 | 90.57 | 88.83 | 90.48 | 88.69 | 89.31 | 89.17 | |
| **85: Blue-Whale** | | 93.01 | 93.01 | 92.93 | 93.01 | 92.49 | 92.66 | 92.71 |
| 92.93 | 92.62 | 92.62 | 92.79 | 92.85 | 92.71 | 92.40 | 92.71 | 92.93 |
| 91.78 | 92.75 | 92.00 | 92.58 | 92.62 | 92.80 | 91.66 | 92.40 | 92.61 |
| 91.66 | 92.57 | 94.48 | 92.49 | 91.70 | 91.79 | 91.74 | 91.97 | 91.66 |
| 89.16 | 90.65 | 90.07 | 90.60 | 94.72 | 94.36 | 89.61 | 89.36 | 89.08 |

| | | | | | | | | |
|---|---|---|---|---|---|---|---|---|
| 90.17 | 90.22 | 89.89 | 90.31 | 89.42 | 92.71 | 93.02 | 92.75 | 89.46 |
| 94.44 | 92.05 | 89.77 | 92.62 | 89.64 | 89.42 | 90.03 | 96.36 | 96.51 |
| 91.82 | 91.82 | 95.53 | 95.58 | 95.84 | 95.71 | 95.45 | 95.10 | 95.23 |
| 95.79 | 93.07 | 93.07 | 95.41 | 96.53 | 94.36 | 91.55 | 88.85 | 95.40 |
| 94.61 | 94.53 | 94.82 | 94.61 | 90.83 | 100.00 | 98.75 | 98.75 | 98.70 |
| 98.53 | 98.92 | 99.39 | 98.66 | 95.31 | 94.83 | 94.59 | 94.76 | 94.46 |
| 94.50 | 94.19 | 94.19 | 94.46 | 94.66 | 94.70 | 94.74 | 94.66 | 94.83 |
| 94.78 | 94.66 | 94.66 | 92.75 | 94.31 | 89.17 | 92.67 | 89.81 | |

| | | | | | | | | |
|---|---|---|---|---|---|---|---|---|
| **86: Dolphin** | | 92.84 | 92.84 | 92.75 | 92.84 | 92.32 | 92.49 | 92.53 |
| 92.75 | 92.45 | 92.45 | 92.62 | 92.67 | 92.53 | 92.19 | 92.45 | 92.75 |
| 91.64 | 92.58 | 91.79 | 92.32 | 92.36 | 92.62 | 91.48 | 92.31 | 92.44 |
| 91.92 | 92.39 | 94.31 | 92.27 | 91.83 | 91.87 | 91.92 | 92.06 | 91.44 |
| 88.90 | 90.61 | 89.94 | 90.55 | 94.41 | 94.10 | 89.57 | 89.49 | 88.90 |
| 90.13 | 90.26 | 89.72 | 90.35 | 89.38 | 92.40 | 92.85 | 92.45 | 89.42 |
| 94.18 | 92.05 | 89.47 | 92.32 | 89.47 | 89.20 | 89.68 | 96.32 | 96.47 |
| 91.78 | 91.78 | 95.53 | 95.49 | 95.84 | 95.49 | 95.06 | 94.93 | 94.84 |
| 95.58 | 92.81 | 92.81 | 95.32 | 96.36 | 94.19 | 91.55 | 88.76 | 95.10 |
| 94.48 | 94.40 | 94.69 | 94.31 | 90.78 | 98.75 | 100.00 | 99.91 | 98.35 |
| 99.70 | 98.87 | 98.49 | 99.83 | 95.10 | 94.61 | 94.37 | 94.55 | 94.16 |
| 94.29 | 93.88 | 93.88 | 94.16 | 94.35 | 94.40 | 94.44 | 94.35 | 94.48 |
| 94.44 | 94.31 | 94.31 | 92.40 | 94.01 | 89.08 | 92.41 | 89.73 | |

**87: Killer-Whale**

| | | | | | | | | |
|---|---|---|---|---|---|---|---|---|
| 92.79 | 92.79 | 92.71 | 92.79 | 92.27 | 92.44 | 92.49 | 92.71 | 92.40 |
| 92.40 | 92.57 | 92.63 | 92.48 | 92.14 | 92.40 | 92.71 | 91.63 | 92.53 |
| 91.78 | 92.27 | 92.31 | 92.57 | 91.35 | 92.27 | 92.39 | 91.86 | 92.34 |
| 94.30 | 92.27 | 91.73 | 91.77 | 91.82 | 92.01 | 91.35 | 88.69 | 90.53 |
| 89.94 | 90.54 | 94.33 | 94.06 | 89.57 | 89.40 | 88.81 | 90.07 | 90.20 |
| 89.64 | 90.30 | 89.29 | 92.36 | 92.85 | 92.40 | 89.33 | 94.17 | 91.96 |
| 89.42 | 92.27 | 89.42 | 89.13 | 89.68 | 96.24 | 96.46 | 91.76 | 91.76 |
| 95.49 | 95.45 | 95.79 | 95.44 | 95.01 | 94.88 | 94.80 | 95.53 | 92.77 |
| 92.77 | 95.27 | 96.31 | 94.14 | 91.50 | 88.66 | 95.01 | 94.52 | 94.43 |
| 94.65 | 94.34 | 90.83 | 98.75 | 99.91 | 100.00 | 98.28 | 99.66 | 98.80 |
| 98.41 | 99.83 | 95.01 | 94.61 | 94.29 | 94.46 | 94.07 | 94.20 | 93.88 |
| 93.88 | 94.07 | 94.35 | 94.39 | 94.43 | 94.35 | 94.48 | 94.43 | 94.30 |
| 94.30 | 92.36 | 94.04 | 89.04 | 92.37 | 89.68 | | | |

| | | | | | | | | |
|---|---|---|---|---|---|---|---|---|
| **88: Sperm-Whale** | | 93.06 | 93.06 | 92.97 | 93.06 | 92.58 | 92.76 | 92.80 |
| 92.97 | 92.63 | 92.67 | 92.84 | 92.59 | 92.75 | 92.54 | 92.80 | 92.97 |
| 91.77 | 92.71 | 91.88 | 92.62 | 92.66 | 92.71 | 91.49 | 92.41 | 92.42 |
| 91.53 | 92.38 | 93.94 | 92.51 | 91.53 | 91.57 | 91.61 | 92.25 | 91.44 |
| 88.81 | 90.28 | 89.97 | 90.48 | 94.33 | 94.19 | 89.46 | 89.34 | 88.88 |
| 89.97 | 90.15 | 89.79 | 90.24 | 89.41 | 92.72 | 92.90 | 92.85 | 89.36 |
| 93.94 | 91.80 | 89.45 | 92.63 | 89.53 | 89.28 | 89.97 | 96.28 | 96.28 |
| 91.75 | 91.75 | 95.58 | 95.58 | 95.84 | 95.71 | 95.36 | 95.06 | 95.15 |
| 95.79 | 92.77 | 92.77 | 95.41 | 96.66 | 94.02 | 91.61 | 88.82 | 95.36 |
| 94.29 | 94.20 | 94.46 | 94.12 | 90.98 | 98.70 | 98.35 | 98.28 | 100.00 |
| 98.11 | 99.06 | 98.97 | 98.27 | 95.23 | 94.37 | 94.38 | 94.55 | 94.12 |
| 94.17 | 94.19 | 94.19 | 94.12 | 94.29 | 94.33 | 94.37 | 94.29 | 94.42 |
| 94.37 | 94.29 | 94.24 | 92.62 | 93.94 | 89.09 | 92.89 | 89.83 | |

**89: Whitebeak-Dolp**

92.62 92.62 92.53 92.62 92.10 92.27 92.27 92.53 92.27
92.27 92.44 92.50 92.35 91.97 92.23 92.57 91.49 92.40
91.61 92.10 92.14 92.44 91.17 92.14 92.17 91.65 92.13
94.09 92.10 91.60 91.56 91.60 91.80 91.18 88.52 90.40
89.77 90.37 94.11 93.84 89.48 89.22 88.64 89.90 90.03
89.46 90.12 89.12 92.18 92.76 92.31 89.16 93.96 91.78
89.25 92.10 89.25 88.96 89.51 96.02 96.25 91.59 91.59
95.31 95.27 95.62 95.27 94.88 94.71 94.67 95.36 92.59
92.59 95.10 96.10 93.97 91.33 88.58 94.79 94.30 94.26
94.43 94.13 90.70 98.53 99.70 99.66 98.11 100.00 98.62
98.24 99.61 94.79 94.39 94.07 94.24 93.86 93.99 93.62
93.62 93.86 94.17 94.26 94.26 94.17 94.30 94.26 94.13
94.13 92.18 93.83 88.86 92.15 89.55

**90: Beaked-whale**

93.11 93.11 93.02 93.11 92.63 92.80 92.85 93.02 92.72
92.80 92.89 92.85 92.80 92.37 92.72 93.02 92.01 92.93
92.10 92.66 92.71 92.97 91.76 92.58 92.47 91.65 92.42
94.03 92.43 91.57 91.65 91.65 92.17 91.45 89.03 90.49
90.23 90.75 94.37 94.23 89.45 89.77 89.15 90.41 90.46
90.05 90.55 89.67 92.72 93.16 92.85 89.44 94.03 92.02
89.75 92.63 89.71 89.45 90.06 96.33 96.28 91.79 91.79
95.71 95.75 96.01 95.80 95.45 95.10 95.23 95.88 93.04
93.04 95.58 96.79 94.36 91.65 88.99 95.44 94.38 94.29
94.55 94.38 91.06 98.92 98.87 98.80 99.06 98.62 100.00
99.36 98.83 95.31 94.51 94.47 94.64 94.16 94.21 94.23
94.23 94.16 94.38 94.42 94.46 94.33 94.51 94.46 94.38
94.33 92.67 94.03 89.08 92.68 89.78

**91: Minke-whale** 93.15 93.15 93.06 93.15 92.67 92.85 92.89
93.06 92.76 92.76 92.93 92.81 92.84 92.46 92.80 93.06
91.97 92.88 91.97 92.66 92.71 92.93 91.71 92.45 92.55
91.70 92.51 94.07 92.56 91.65 91.74 91.74 92.17 91.58
89.03 90.41 90.14 90.75 94.50 94.36 89.46 89.64 89.11
90.37 90.37 89.97 90.46 89.59 92.76 93.12 92.85 89.44
94.07 92.02 89.80 92.68 89.75 89.49 90.01 96.33 96.24
91.71 91.71 95.54 95.58 95.84 95.71 95.45 95.10 95.19
95.80 93.12 93.12 95.49 96.79 94.41 91.65 88.86 95.57
94.38 94.29 94.55 94.38 90.89 99.39 98.49 98.41 98.97
98.24 99.36 100.00 98.40 95.44 94.55 94.55 94.73 94.34
94.34 94.36 94.36 94.29 94.46 94.51 94.55 94.42 94.64
94.59 94.51 94.46 92.84 94.12 89.09 92.68 89.82

**92: Common-Dolphin**

92.75 92.75 92.66 92.75 92.23 92.40 92.45 92.66 92.36
92.36 92.53 92.59 92.44 92.10 92.36 92.66 91.54 92.49
91.83 92.23 92.27 92.53 91.44 92.36 92.35 91.96 92.31
94.39 92.18 91.79 91.83 91.87 92.01 91.35 88.90 90.57
89.90 90.55 94.33 94.02 89.52 89.44 88.86 90.13 90.26
89.72 90.35 89.38 92.36 92.85 92.45 89.33 94.09 91.96
89.47 92.32 89.42 89.20 89.64 96.23 96.38 91.69 91.69
95.49 95.45 95.79 95.45 95.02 94.89 94.80 95.53 92.77
92.77 95.28 96.31 94.19 91.47 88.68 94.97 94.48 94.40

94.69 94.31 90.70 98.66 99.83 99.83 98.27 99.61 98.83
98.40 100.00 94.97 94.57 94.33 94.50 94.07 94.20 93.88
93.88 94.07 94.35 94.40 94.44 94.35 94.48 94.44 94.31
94.31 92.40 94.01 89.08 92.37 89.73

**93: Warthog** 93.12 93.12 93.03 93.12 92.77 92.86 92.86
92.99 92.73 92.68 92.85 92.71 92.76 92.12 92.47 92.90
91.83 92.75 92.06 92.27 92.36 92.75 91.97 92.12 92.31
91.25 92.31 93.71 92.19 91.38 91.51 91.47 91.71 91.11
88.74 90.21 89.74 89.93 94.28 94.48 89.73 89.77 88.66
89.68 89.64 89.52 89.95 88.92 92.38 92.58 92.38 89.26
93.97 92.10 89.31 92.34 89.35 89.18 89.48 95.66 95.49
91.32 91.23 95.04 95.09 95.34 95.26 94.79 94.79 94.57
95.34 92.76 92.72 95.05 95.39 93.87 91.66 88.85 99.66
93.97 93.88 94.19 93.97 90.61 95.31 95.10 95.01 95.23
94.79 95.31 95.44 94.97 100.00 93.79 94.01 94.10 94.10
94.19 94.26 94.26 94.23 94.10 94.10 94.18 94.10 94.27
94.18 94.05 94.05 92.62 93.84 88.51 92.54 89.19

**94: Brown-Bear** 94.14 94.18 94.10 94.14 93.58 93.75 93.71
93.97 93.49 93.62 93.83 93.85 93.75 92.97 93.32 93.92
92.82 93.75 92.70 93.14 93.27 93.62 92.22 92.92 92.66
91.74 92.62 94.70 92.45 91.96 92.13 91.70 91.89 91.79
89.38 91.01 89.82 90.64 97.71 97.31 90.01 89.45 89.25
90.30 90.39 89.85 90.70 89.69 92.93 92.72 93.06 89.21
97.55 92.18 89.82 92.88 89.69 89.77 89.60 95.07 95.26
91.86 91.86 94.19 94.06 94.41 94.23 93.89 93.58 93.67
94.28 93.60 93.55 94.02 94.15 94.58 92.33 89.60 93.88
97.93 97.80 98.19 97.85 90.57 94.83 94.61 94.61 94.37
94.39 94.51 94.55 94.57 93.79 100.00 99.52 99.70 97.40
97.45 97.14 97.14 97.45 97.12 97.03 97.20 97.16 97.29
97.29 97.16 97.16 94.92 97.37 89.22 93.15 90.17

**95: Black-Bear** 94.32 94.36 94.28 94.32 93.80 93.97 93.93
94.10 93.67 93.80 94.01 93.73 93.93 93.02 93.45 94.10
92.86 93.84 92.80 93.14 93.27 93.66 92.36 93.02 92.82
91.66 92.78 94.59 92.69 91.96 92.09 91.70 92.17 91.88
89.55 91.04 89.99 90.65 97.80 97.49 90.05 89.58 89.43
90.36 90.40 89.95 90.71 89.74 93.06 92.94 93.19 89.36
97.32 92.36 89.99 92.98 89.74 89.87 89.77 95.14 95.03
91.79 91.79 94.32 94.20 94.54 94.37 94.02 93.76 93.81
94.41 93.60 93.56 94.20 94.19 94.58 92.56 89.69 94.10
97.75 97.62 97.97 97.66 90.76 94.59 94.37 94.29 94.38
94.07 94.47 94.55 94.33 94.01 99.52 100.00 99.83 97.59
97.59 97.31 97.31 97.59 96.93 96.84 97.02 96.97 97.10
97.10 96.97 96.97 95.10 97.15 89.29 93.28 90.36

**96: Polar-Bear** 94.41 94.45 94.36 94.41 93.89 94.06 94.02
94.19 93.76 93.89 94.10 93.82 94.01 93.19 93.54 94.19
92.96 93.92 92.88 93.23 93.36 93.75 92.45 93.10 92.91
91.75 92.86 94.68 92.73 92.01 92.18 91.79 92.26 91.97
89.60 91.13 90.03 90.78 97.84 97.53 90.09 89.62 89.47
90.40 90.44 89.99 90.75 89.78 93.15 92.94 93.28 89.44
97.40 92.44 90.03 93.06 89.78 89.91 89.82 95.31 95.20

| | | | | | | | | |
|---|---|---|---|---|---|---|---|---|
| 91.88 | 91.88 | 94.50 | 94.37 | 94.71 | 94.54 | 94.20 | 93.89 | 93.98 |
| 94.58 | 93.69 | 93.64 | 94.37 | 94.36 | 94.67 | 92.60 | 89.73 | 94.19 |
| 97.84 | 97.71 | 98.05 | 97.75 | 90.85 | 94.76 | 94.55 | 94.46 | 94.55 |
| 94.24 | 94.64 | 94.73 | 94.50 | 94.10 | 99.70 | 99.83 | 100.00 | 97.63 |
| 97.63 | 97.40 | 97.40 | 97.63 | 97.02 | 96.93 | 97.10 | 97.06 | 97.19 |
| 97.19 | 97.06 | 97.06 | 95.18 | 97.23 | 89.33 | 93.46 | 90.36 | |
| **97: Wolverine** | | 93.93 | 93.97 | 93.89 | 93.93 | 93.50 | 93.67 | 93.58 |
| 93.80 | 93.37 | 93.58 | 93.71 | 93.42 | 93.62 | 92.67 | 93.19 | 93.71 |
| 92.49 | 93.58 | 92.49 | 92.88 | 93.01 | 93.36 | 92.19 | 93.10 | 92.91 |
| 91.53 | 92.91 | 94.76 | 92.90 | 91.83 | 91.83 | 91.66 | 92.00 | 91.75 |
| 89.90 | 91.13 | 90.29 | 91.08 | 99.35 | 98.87 | 90.26 | 90.01 | 89.82 |
| 90.75 | 90.83 | 90.55 | 90.92 | 90.00 | 92.89 | 92.81 | 92.93 | 89.97 |
| 98.92 | 92.57 | 90.25 | 92.80 | 90.04 | 90.34 | 90.03 | 94.96 | 94.94 |
| 92.01 | 92.01 | 94.45 | 94.37 | 94.67 | 94.50 | 94.24 | 93.89 | 93.98 |
| 94.54 | 93.34 | 93.30 | 94.33 | 94.28 | 94.32 | 92.86 | 89.90 | 94.10 |
| 97.79 | 97.62 | 97.97 | 97.49 | 90.80 | 94.46 | 94.16 | 94.07 | 94.12 |
| 93.86 | 94.16 | 94.34 | 94.07 | 94.10 | 97.40 | 97.59 | 97.63 | 100.00 |
| 99.14 | 97.31 | 97.31 | 99.14 | 96.80 | 96.71 | 96.89 | 96.89 | 96.97 |
| 96.97 | 96.84 | 96.84 | 95.01 | 97.10 | 89.68 | 93.28 | 90.41 | |
| **98: Ermine** | | 93.84 | 93.89 | 93.80 | 93.84 | 93.41 | 93.58 | 93.50 |
| 93.71 | 93.37 | 93.54 | 93.62 | 93.38 | 93.54 | 92.59 | 93.15 | 93.63 |
| 92.44 | 93.49 | 92.40 | 92.84 | 92.97 | 93.36 | 92.06 | 92.97 | 92.91 |
| 91.49 | 92.91 | 94.68 | 92.86 | 91.83 | 91.79 | 91.62 | 91.87 | 91.71 |
| 89.94 | 91.22 | 90.34 | 91.17 | 99.61 | 99.31 | 90.39 | 90.10 | 89.86 |
| 90.83 | 90.83 | 90.60 | 90.97 | 90.04 | 92.89 | 92.77 | 92.93 | 89.75 |
| 98.88 | 92.49 | 90.25 | 92.80 | 90.17 | 90.30 | 90.12 | 94.96 | 94.98 |
| 91.83 | 91.83 | 94.32 | 94.24 | 94.54 | 94.41 | 94.24 | 93.81 | 93.98 |
| 94.45 | 93.43 | 93.38 | 94.24 | 94.19 | 94.32 | 92.82 | 89.90 | 94.19 |
| 97.84 | 97.66 | 98.01 | 97.53 | 90.85 | 94.50 | 94.29 | 94.20 | 94.17 |
| 93.99 | 94.21 | 94.34 | 94.20 | 94.19 | 97.45 | 97.59 | 97.63 | 99.14 |
| 100.00 | 97.22 | 97.22 | 99.57 | 96.84 | 96.67 | 96.84 | 96.84 | 96.89 |
| 96.93 | 96.80 | 96.76 | 94.92 | 97.06 | 89.68 | 93.24 | 90.45 | |
| **99: Dog** | | 93.94 | 93.99 | 93.90 | 93.94 | 93.55 | 93.73 | 93.64 |
| 93.81 | 93.38 | 93.60 | 93.81 | 93.67 | 93.72 | 92.68 | 93.25 | 93.73 |
| 92.77 | 93.70 | 92.63 | 92.97 | 93.10 | 93.53 | 92.24 | 93.20 | 92.84 |
| 91.73 | 92.84 | 94.49 | 92.67 | 92.03 | 92.03 | 91.73 | 92.19 | 91.68 |
| 89.23 | 90.86 | 89.92 | 90.54 | 97.40 | 97.50 | 89.95 | 89.69 | 89.19 |
| 90.21 | 90.38 | 89.91 | 90.56 | 89.62 | 92.82 | 92.84 | 92.86 | 89.66 |
| 97.14 | 92.24 | 89.75 | 92.73 | 89.66 | 89.79 | 89.83 | 94.93 | 94.84 |
| 91.76 | 91.76 | 94.28 | 94.27 | 94.49 | 94.32 | 93.93 | 93.80 | 93.72 |
| 94.36 | 93.85 | 93.81 | 94.10 | 94.27 | 94.66 | 92.75 | 89.72 | 94.39 |
| 97.44 | 97.31 | 97.66 | 97.31 | 90.61 | 94.19 | 93.88 | 93.88 | 94.19 |
| 93.62 | 94.23 | 94.36 | 93.88 | 94.26 | 97.14 | 97.31 | 97.40 | 97.31 |
| 97.22 | 100.00 | 100.00 | 97.22 | 96.70 | 96.62 | 96.79 | 96.70 | 96.92 |
| 96.92 | 96.79 | 96.79 | 95.31 | 99.18 | 89.43 | 93.53 | 90.36 | |
| **100: Dingo** | | 93.94 | 93.99 | 93.90 | 93.94 | 93.55 | 93.73 | 93.64 |
| 93.81 | 93.38 | 93.60 | 93.81 | 93.67 | 93.72 | 92.68 | 93.25 | 93.73 |
| 92.77 | 93.70 | 92.63 | 92.97 | 93.10 | 93.53 | 92.24 | 93.20 | 92.84 |
| 91.73 | 92.84 | 94.49 | 92.67 | 92.03 | 92.03 | 91.73 | 92.19 | 91.68 |
| 89.23 | 90.86 | 89.92 | 90.54 | 97.40 | 97.50 | 89.95 | 89.69 | 89.19 |

| | | | | | | | | |
|---|---|---|---|---|---|---|---|---|
| 90.21 | 90.38 | 89.91 | 90.56 | 89.62 | 92.82 | 92.84 | 92.86 | 89.66 |
| 97.14 | 92.24 | 89.75 | 92.73 | 89.66 | 89.79 | 89.83 | 94.93 | 94.84 |
| 91.76 | 91.76 | 94.28 | 94.27 | 94.49 | 94.32 | 93.93 | 93.80 | 93.72 |
| 94.36 | 93.85 | 93.81 | 94.10 | 94.27 | 94.66 | 92.75 | 89.72 | 94.39 |
| 97.44 | 97.31 | 97.66 | 97.31 | 90.61 | 94.19 | 93.88 | 93.88 | 94.19 |
| 93.62 | 94.23 | 94.36 | 93.88 | 94.26 | 97.14 | 97.31 | 97.40 | 97.31 |
| 97.22 | 100.00 | 100.00 | 97.22 | 96.70 | 96.62 | 96.79 | 96.70 | 96.92 |
| 96.92 | 96.79 | 96.79 | 95.31 | 99.18 | 89.43 | 93.53 | 90.36 | |
| **101: Ferret** | | 93.89 | 93.93 | 93.84 | 93.89 | 93.45 | 93.63 | 93.54 |
| 93.76 | 93.32 | 93.58 | 93.67 | 93.42 | 93.58 | 92.72 | 93.24 | 93.67 |
| 92.49 | 93.53 | 92.49 | 92.93 | 93.06 | 93.32 | 92.14 | 93.02 | 92.91 |
| 91.49 | 92.91 | 94.72 | 92.82 | 91.79 | 91.79 | 91.62 | 91.96 | 91.71 |
| 90.07 | 91.26 | 90.42 | 91.26 | 99.61 | 99.48 | 90.48 | 90.06 | 89.99 |
| 90.92 | 90.92 | 90.64 | 91.06 | 90.13 | 92.93 | 92.81 | 93.06 | 89.79 |
| 98.88 | 92.57 | 90.29 | 92.85 | 90.21 | 90.39 | 90.21 | 94.96 | 94.98 |
| 91.88 | 91.88 | 94.32 | 94.24 | 94.54 | 94.41 | 94.20 | 93.81 | 93.94 |
| 94.45 | 93.43 | 93.38 | 94.24 | 94.23 | 94.41 | 92.82 | 90.03 | 94.23 |
| 97.84 | 97.66 | 98.01 | 97.58 | 90.93 | 94.46 | 94.16 | 94.07 | 94.12 |
| 93.86 | 94.16 | 94.29 | 94.07 | 94.23 | 97.45 | 97.59 | 97.63 | 99.14 |
| 99.57 | 97.22 | 97.22 | 100.00 | 96.89 | 96.71 | 96.89 | 96.89 | 96.93 |
| 96.93 | 96.80 | 96.80 | 94.97 | 97.06 | 89.68 | 93.24 | 90.45 | |
| **102: Cat** | | 93.75 | 93.79 | 93.71 | 93.75 | 93.23 | 93.49 | 93.27 |
| 93.66 | 93.19 | 93.45 | 93.53 | 93.63 | 93.44 | 92.62 | 93.14 | 93.53 |
| 92.63 | 93.49 | 92.74 | 93.06 | 93.14 | 93.36 | 92.31 | 93.31 | 92.66 |
| 91.79 | 92.62 | 94.61 | 92.41 | 91.83 | 92.09 | 91.70 | 92.15 | 91.79 |
| 89.47 | 90.96 | 89.99 | 90.68 | 97.10 | 96.75 | 89.97 | 89.58 | 89.43 |
| 90.30 | 90.39 | 90.07 | 90.61 | 89.60 | 92.97 | 92.55 | 93.01 | 89.73 |
| 96.94 | 92.18 | 89.77 | 92.88 | 89.78 | 89.81 | 89.73 | 95.03 | 95.22 |
| 91.91 | 91.91 | 93.97 | 93.97 | 94.19 | 94.15 | 93.93 | 93.45 | 93.71 |
| 94.19 | 93.68 | 93.64 | 93.97 | 94.10 | 94.62 | 92.59 | 89.77 | 94.14 |
| 97.24 | 97.29 | 97.50 | 97.07 | 90.66 | 94.66 | 94.35 | 94.35 | 94.29 |
| 94.17 | 94.38 | 94.46 | 94.35 | 94.10 | 97.12 | 96.93 | 97.02 | 96.80 |
| 96.84 | 96.70 | 96.70 | 96.89 | 100.00 | 99.44 | 99.61 | 99.48 | 99.48 |
| 99.48 | 99.35 | 99.35 | 97.01 | 96.94 | 89.61 | 93.28 | 90.39 | |
| **103: Puma** | | 93.66 | 93.71 | 93.62 | 93.66 | 93.14 | 93.40 | 93.19 |
| 93.58 | 93.10 | 93.32 | 93.44 | 93.50 | 93.36 | 92.58 | 93.10 | 93.45 |
| 92.58 | 93.40 | 92.57 | 93.01 | 93.10 | 93.27 | 92.13 | 93.36 | 92.62 |
| 91.83 | 92.57 | 94.53 | 92.36 | 91.83 | 92.00 | 91.79 | 92.11 | 91.83 |
| 89.47 | 90.79 | 89.90 | 90.47 | 96.93 | 96.57 | 90.01 | 89.45 | 89.43 |
| 90.09 | 90.26 | 90.07 | 90.48 | 89.47 | 92.93 | 92.46 | 92.97 | 89.69 |
| 96.86 | 92.05 | 89.77 | 92.84 | 89.82 | 89.77 | 89.60 | 94.98 | 95.26 |
| 91.95 | 91.95 | 94.02 | 94.02 | 94.23 | 94.19 | 93.97 | 93.54 | 93.76 |
| 94.23 | 93.77 | 93.73 | 94.02 | 94.23 | 94.62 | 92.55 | 89.73 | 94.14 |
| 97.20 | 97.24 | 97.45 | 97.03 | 90.70 | 94.70 | 94.40 | 94.39 | 94.33 |
| 94.26 | 94.42 | 94.51 | 94.40 | 94.10 | 97.03 | 96.84 | 96.93 | 96.71 |
| 96.67 | 96.62 | 96.62 | 96.71 | 99.44 | 100.00 | 99.74 | 99.44 | 99.53 |
| 99.44 | 99.31 | 99.35 | 96.88 | 96.90 | 89.79 | 93.24 | 90.47 | |
| **104: Cheetah** | | 93.84 | 93.88 | 93.79 | 93.84 | 93.32 | 93.58 | 93.36 |
| 93.75 | 93.27 | 93.49 | 93.62 | 93.72 | 93.53 | 92.71 | 93.23 | 93.62 |
| 92.82 | 93.62 | 92.74 | 93.19 | 93.27 | 93.49 | 92.31 | 93.44 | 92.79 |

| | | | | | | | | |
|---|---|---|---|---|---|---|---|---|
| 92.00 | 92.75 | 94.70 | 92.54 | 92.00 | 92.17 | 91.92 | 92.23 | 92.01 |
| 89.56 | 90.96 | 90.03 | 90.64 | 97.10 | 96.75 | 90.10 | 89.67 | 89.51 |
| 90.26 | 90.43 | 90.16 | 90.66 | 89.64 | 93.06 | 92.59 | 93.10 | 89.77 |
| 97.03 | 92.22 | 89.86 | 92.97 | 89.91 | 89.94 | 89.82 | 95.16 | 95.44 |
| 92.08 | 92.08 | 94.10 | 94.10 | 94.32 | 94.28 | 94.06 | 93.63 | 93.84 |
| 94.32 | 93.82 | 93.77 | 94.10 | 94.28 | 94.75 | 92.59 | 89.86 | 94.23 |
| 97.37 | 97.42 | 97.63 | 97.20 | 90.74 | 94.74 | 94.44 | 94.43 | 94.37 |
| 94.26 | 94.46 | 94.55 | 94.44 | 94.18 | 97.20 | 97.02 | 97.10 | 96.89 |
| 96.84 | 96.79 | 96.79 | 96.89 | 99.61 | 99.74 | 100.00 | 99.61 | 99.61 |
| 99.61 | 99.48 | 99.48 | 97.05 | 97.03 | 89.83 | 93.41 | 90.52 | |

**105: Bengal-Panther**

| | | | | | | | | |
|---|---|---|---|---|---|---|---|---|
| 93.75 | 93.79 | 93.71 | 93.75 | 93.23 | 93.49 | 93.27 | 93.66 | 93.19 |
| 93.40 | 93.53 | 93.59 | 93.44 | 92.71 | 93.14 | 93.53 | 92.63 | 93.49 |
| 92.70 | 93.14 | 93.23 | 93.36 | 92.26 | 93.23 | 92.66 | 91.83 | 92.62 |
| 94.53 | 92.41 | 91.87 | 92.04 | 91.74 | 92.15 | 91.92 | 89.47 | 91.01 |
| 89.90 | 90.68 | 97.14 | 96.75 | 90.01 | 89.49 | 89.43 | 90.30 | 90.43 |
| 90.07 | 90.66 | 89.64 | 93.06 | 92.55 | 93.10 | 89.77 | 97.03 | 92.13 |
| 89.77 | 92.97 | 89.82 | 89.86 | 89.64 | 94.94 | 95.22 | 92.08 | 92.08 |
| 93.93 | 93.93 | 94.15 | 94.10 | 93.93 | 93.54 | 93.67 | 94.15 | 93.68 |
| 93.64 | 93.93 | 94.10 | 94.66 | 92.46 | 89.64 | 94.14 | 97.29 | 97.33 |
| 97.54 | 97.11 | 90.74 | 94.66 | 94.35 | 94.35 | 94.29 | 94.17 | 94.33 |
| 94.42 | 94.35 | 94.10 | 97.16 | 96.97 | 97.06 | 96.89 | 96.84 | 96.70 |
| 96.70 | 96.89 | 99.48 | 99.44 | 99.61 | 100.00 | 99.44 | 99.44 | 99.31 |
| 99.31 | 96.92 | 96.90 | 89.79 | 93.28 | 90.39 | | | |

| | | | | | | | | |
|---|---|---|---|---|---|---|---|---|
| **106: Tiger** | | 93.92 | 93.97 | 93.88 | 93.92 | 93.40 | 93.66 | 93.45 |
| 93.84 | 93.36 | 93.58 | 93.70 | 93.76 | 93.62 | 92.80 | 93.32 | 93.71 |
| 92.82 | 93.66 | 92.74 | 93.23 | 93.32 | 93.53 | 92.35 | 93.40 | 92.75 |
| 91.96 | 92.70 | 94.74 | 92.58 | 92.00 | 92.35 | 91.92 | 92.32 | 91.97 |
| 89.60 | 90.96 | 90.08 | 90.77 | 97.14 | 96.79 | 90.05 | 89.67 | 89.56 |
| 90.39 | 90.52 | 90.11 | 90.79 | 89.77 | 93.14 | 92.72 | 93.10 | 89.86 |
| 97.07 | 92.35 | 89.90 | 93.06 | 89.91 | 89.99 | 89.82 | 95.16 | 95.44 |
| 92.04 | 92.04 | 94.10 | 94.10 | 94.32 | 94.28 | 94.02 | 93.58 | 93.80 |
| 94.32 | 93.82 | 93.77 | 94.02 | 94.32 | 94.79 | 92.68 | 89.90 | 94.31 |
| 97.46 | 97.50 | 97.71 | 97.29 | 90.70 | 94.83 | 94.48 | 94.48 | 94.42 |
| 94.30 | 94.51 | 94.64 | 94.48 | 94.27 | 97.29 | 97.10 | 97.19 | 96.97 |
| 96.89 | 96.92 | 96.92 | 96.93 | 99.48 | 99.53 | 99.61 | 99.44 | 100.00 |
| 99.91 | 99.78 | 99.78 | 97.35 | 97.20 | 89.74 | 93.50 | 90.47 | |

| | | | | | | | | |
|---|---|---|---|---|---|---|---|---|
| **107: Lion** | | 93.88 | 93.92 | 93.84 | 93.88 | 93.36 | 93.62 | 93.40 |
| 93.79 | 93.32 | 93.53 | 93.66 | 93.72 | 93.57 | 92.75 | 93.27 | 93.66 |
| 92.77 | 93.62 | 92.70 | 93.19 | 93.27 | 93.49 | 92.31 | 93.36 | 92.75 |
| 91.96 | 92.70 | 94.74 | 92.58 | 92.00 | 92.35 | 91.87 | 92.28 | 91.97 |
| 89.56 | 90.92 | 90.03 | 90.73 | 97.14 | 96.79 | 90.01 | 89.62 | 89.51 |
| 90.35 | 90.48 | 90.07 | 90.74 | 89.73 | 93.10 | 92.72 | 93.06 | 89.82 |
| 97.07 | 92.31 | 89.86 | 93.01 | 89.87 | 89.94 | 89.77 | 95.07 | 95.35 |
| 91.99 | 91.99 | 94.06 | 94.06 | 94.28 | 94.23 | 93.97 | 93.54 | 93.76 |
| 94.28 | 93.86 | 93.82 | 93.97 | 94.23 | 94.75 | 92.64 | 89.86 | 94.23 |
| 97.46 | 97.50 | 97.71 | 97.29 | 90.66 | 94.78 | 94.44 | 94.43 | 94.37 |
| 94.26 | 94.46 | 94.59 | 94.44 | 94.18 | 97.29 | 97.10 | 97.19 | 96.97 |
| 96.93 | 96.92 | 96.92 | 96.93 | 99.48 | 99.44 | 99.61 | 99.44 | 99.91 |
| 100.00 | 99.87 | 99.78 | 97.35 | 97.16 | 89.70 | 93.50 | 90.43 | |

**108: Leopard** 93.75 93.79 93.71 93.75 93.23 93.49 93.27
93.66 93.19 93.40 93.53 93.59 93.44 92.62 93.14 93.53
92.63 93.49 92.57 93.06 93.14 93.36 92.22 93.23 92.62
91.87 92.57 94.61 92.45 91.87 92.22 91.74 92.15 91.83
89.43 90.79 89.90 90.60 97.01 96.66 89.88 89.54 89.38
90.22 90.35 89.94 90.70 89.60 92.97 92.59 92.93 89.69
96.94 92.18 89.73 92.88 89.74 89.81 89.64 95.03 95.31
91.86 91.86 93.93 93.93 94.15 94.10 93.84 93.41 93.63
94.15 93.73 93.68 93.84 94.15 94.62 92.51 89.81 94.10
97.33 97.37 97.58 97.16 90.61 94.66 94.31 94.30 94.29
94.13 94.38 94.51 94.31 94.05 97.16 96.97 97.06 96.84
96.80 96.79 96.79 96.80 99.35 99.31 99.48 99.31 99.78
99.87 100.00 99.66 97.22 97.03 89.61 93.37 90.34

**109: Snow-Leopard**

93.79 93.84 93.75 93.79 93.27 93.53 93.32 93.71 93.23
93.45 93.57 93.68 93.49 92.71 93.23 93.58 92.63 93.49
92.61 93.10 93.19 93.36 92.22 93.27 92.62 91.83 92.57
94.61 92.45 91.87 92.22 91.74 92.15 91.92 89.43 90.88
89.95 90.64 97.01 96.66 89.92 89.54 89.38 90.26 90.48
89.94 90.74 89.69 93.19 92.68 93.06 89.73 96.94 92.22
89.73 93.01 89.74 89.81 89.64 94.94 95.22 91.99 91.99
93.93 93.93 94.15 94.10 93.84 93.41 93.63 94.15 93.73
93.68 93.84 94.19 94.71 92.59 89.90 94.10 97.33 97.37
97.58 97.16 90.57 94.66 94.31 94.30 94.24 94.13 94.33
94.46 94.31 94.05 97.16 96.97 97.06 96.84 96.76 96.79
96.79 96.80 99.35 99.35 99.48 99.31 99.78 99.78 99.66
100.00 97.22 97.03 89.57 93.37 90.34

**110: Jaguar** 92.25 92.25 92.21 92.21 91.73 91.95 91.77
92.08 91.69 91.90 92.03 91.89 91.94 91.08 91.60 92.03
90.98 91.97 90.85 91.41 91.50 91.84 90.54 91.73 91.14
90.00 91.14 92.84 90.93 90.04 90.30 90.00 90.54 90.11
87.61 89.25 88.14 88.80 95.06 95.26 88.33 87.90 87.62
88.43 88.60 88.21 88.82 87.88 91.47 91.07 91.47 88.04
94.75 90.63 87.96 91.39 88.01 88.01 87.92 93.36 93.23
89.89 89.89 92.55 92.55 92.76 92.72 92.46 92.03 92.24
92.76 91.94 91.90 92.46 92.67 93.06 90.89 87.76 92.67
95.10 95.14 95.31 94.97 88.83 92.75 92.40 92.36 92.62
92.18 92.67 92.84 92.40 92.62 94.92 95.10 95.18 95.01
94.92 95.31 95.31 94.97 97.01 96.88 97.05 96.92 97.35
97.35 97.22 97.22 100.00 94.75 87.85 91.89 88.71

**111: Arctic-Fox** 93.84 93.88 93.79 93.84 93.45 93.62 93.45
93.75 93.27 93.49 93.70 93.72 93.62 92.58 93.14 93.62
92.63 93.58 92.66 92.93 93.06 93.40 92.22 93.14 92.53
91.57 92.49 94.44 92.36 91.87 91.92 91.61 91.89 91.62
89.25 90.70 89.82 90.47 97.36 96.92 89.75 89.54 89.17
90.13 90.26 89.90 90.48 89.47 92.62 92.55 92.62 89.51
97.42 92.13 89.69 92.53 89.65 89.77 89.69 94.64 94.92
91.95 91.95 93.89 93.89 94.10 93.97 93.63 93.41 93.41
94.02 93.86 93.82 93.80 93.84 94.58 92.51 89.73 93.97
97.59 97.50 97.84 97.46 90.48 94.31 94.01 94.04 93.94

| | | | | | | | | |
|---|---|---|---|---|---|---|---|---|
| 93.83 | 94.03 | 94.12 | 94.01 | 93.84 | 97.37 | 97.15 | 97.23 | 97.10 |
| 97.06 | 99.18 | 99.18 | 97.06 | 96.94 | 96.90 | 97.03 | 96.90 | 97.20 |
| 97.16 | 97.03 | 97.03 | 94.75 | 100.00 | 89.39 | 92.93 | 90.39 | |
| **112: Armadillo** | | 89.69 | 89.74 | 89.69 | 89.69 | 89.30 | 89.48 | 89.48 |
| 89.52 | 89.21 | 89.61 | 89.69 | 89.64 | 89.61 | 89.21 | 89.43 | 89.74 |
| 88.33 | 89.68 | 88.72 | 89.38 | 89.47 | 89.55 | 88.72 | 89.03 | 87.87 |
| 86.87 | 87.91 | 88.91 | 87.96 | 86.92 | 87.18 | 87.05 | 87.75 | 87.45 |
| 85.92 | 87.09 | 86.80 | 86.95 | 89.85 | 89.69 | 87.00 | 86.59 | 85.80 |
| 86.70 | 86.91 | 86.79 | 87.05 | 86.10 | 89.26 | 87.40 | 89.26 | 86.08 |
| 89.48 | 88.90 | 86.71 | 89.17 | 86.33 | 86.49 | 86.67 | 89.37 | 89.48 |
| 89.46 | 89.46 | 88.74 | 88.70 | 88.78 | 88.83 | 88.83 | 88.39 | 88.57 |
| 88.83 | 89.04 | 89.00 | 88.66 | 88.91 | 89.17 | 89.01 | 86.52 | 88.55 |
| 89.74 | 89.57 | 89.87 | 89.31 | 88.69 | 89.17 | 89.08 | 89.04 | 89.09 |
| 88.86 | 89.08 | 89.09 | 89.08 | 88.51 | 89.22 | 89.29 | 89.33 | 89.68 |
| 89.68 | 89.43 | 89.43 | 89.68 | 89.61 | 89.79 | 89.83 | 89.79 | 89.74 |
| 89.70 | 89.61 | 89.57 | 87.85 | 89.39 | 100.00 | 87.82 | 91.34 | |
| **113: Pangolin** | | 91.92 | 91.92 | 91.83 | 91.92 | 91.57 | 91.61 | 91.74 |
| 91.79 | 91.48 | 91.57 | 91.65 | 91.42 | 91.56 | 91.14 | 91.53 | 91.74 |
| 90.56 | 91.50 | 91.12 | 91.29 | 91.37 | 91.46 | 90.60 | 91.05 | 91.24 |
| 90.52 | 91.24 | 93.28 | 91.16 | 90.57 | 90.74 | 90.52 | 90.86 | 90.29 |
| 87.98 | 89.48 | 88.85 | 89.16 | 93.33 | 93.45 | 88.53 | 87.87 | 87.94 |
| 88.87 | 88.79 | 88.88 | 89.14 | 88.24 | 90.92 | 90.86 | 91.05 | 87.84 |
| 93.11 | 90.42 | 88.45 | 90.83 | 88.15 | 88.41 | 88.72 | 93.02 | 92.85 |
| 90.20 | 90.16 | 92.33 | 92.33 | 92.59 | 92.37 | 91.99 | 91.90 | 91.86 |
| 92.46 | 91.60 | 91.64 | 92.20 | 92.63 | 92.54 | 90.63 | 87.91 | 92.62 |
| 93.54 | 93.32 | 93.58 | 93.32 | 89.31 | 92.67 | 92.41 | 92.37 | 92.89 |
| 92.15 | 92.68 | 92.68 | 92.37 | 92.54 | 93.15 | 93.28 | 93.46 | 93.28 |
| 93.24 | 93.53 | 93.53 | 93.24 | 93.28 | 93.24 | 93.41 | 93.28 | 93.50 |
| 93.50 | 93.37 | 93.37 | 91.89 | 92.93 | 87.82 | 100.00 | 88.59 | |
| **114: Sloth** | | 90.76 | 90.72 | 90.67 | 90.67 | 90.20 | 90.37 | 90.20 |
| 90.50 | 90.28 | 90.41 | 90.59 | 90.24 | 90.50 | 89.85 | 90.41 | 90.67 |
| 89.03 | 90.36 | 89.79 | 89.84 | 89.97 | 90.36 | 89.66 | 89.89 | 88.44 |
| 87.49 | 88.44 | 89.73 | 88.30 | 87.41 | 87.49 | 87.62 | 88.04 | 87.81 |
| 86.20 | 87.98 | 87.03 | 87.24 | 90.66 | 90.46 | 87.34 | 87.22 | 86.12 |
| 86.91 | 87.22 | 86.81 | 87.52 | 86.38 | 89.59 | 88.34 | 89.76 | 86.86 |
| 90.21 | 89.54 | 86.90 | 89.59 | 86.77 | 86.77 | 86.85 | 90.36 | 90.34 |
| 89.62 | 89.62 | 89.41 | 89.38 | 89.59 | 89.59 | 89.55 | 89.12 | 89.42 |
| 89.59 | 89.54 | 89.49 | 89.38 | 89.63 | 89.98 | 89.51 | 86.87 | 89.32 |
| 90.69 | 90.60 | 90.78 | 90.21 | 89.17 | 89.81 | 89.73 | 89.68 | 89.83 |
| 89.55 | 89.78 | 89.82 | 89.73 | 89.19 | 90.17 | 90.36 | 90.36 | 90.41 |
| 90.45 | 90.36 | 90.36 | 90.45 | 90.39 | 90.47 | 90.52 | 90.39 | 90.47 |
| 90.43 | 90.34 | 90.34 | 88.71 | 90.39 | 91.34 | 88.59 | 100.00 | |

**Supplementary Figure 1.** The multiple sequence alignment of human NOTCH3 with 113 mammals using CLUSTAL-Omega. The underlined figures against each mammal correspond to the % identity with human.

**a**

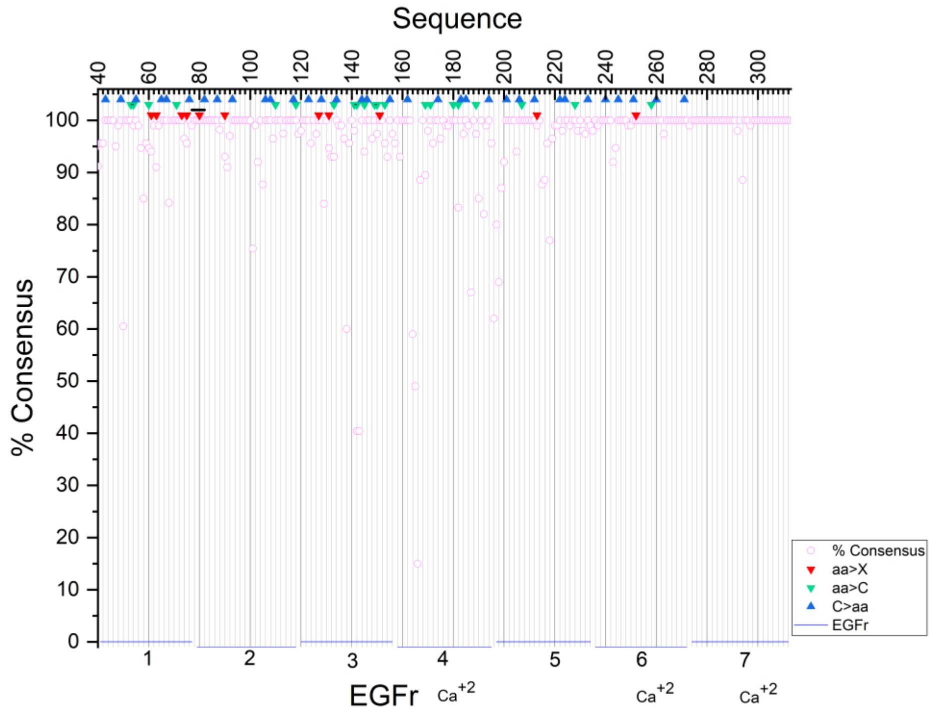
Sequence
% Consensus
EGFr
Ca+2
Ca+2
Ca+2
% Consensus
aa>X
aa>C
C>aa
EGFr

**b**

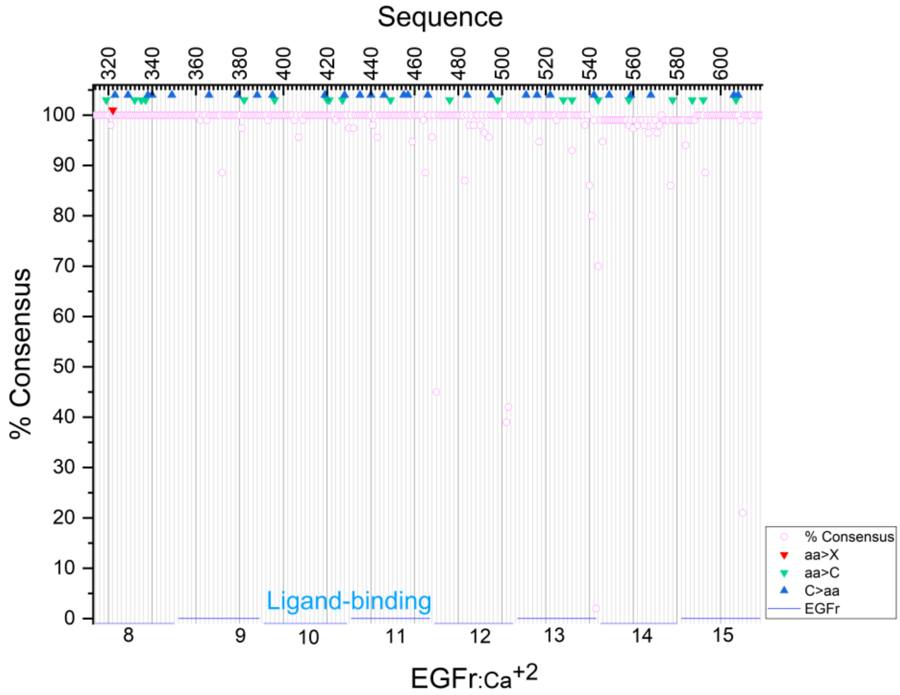
Sequence
% Consensus
Ligand-binding
EGFr:Ca+2
% Consensus
aa>X
aa>C
C>aa
EGFr

c

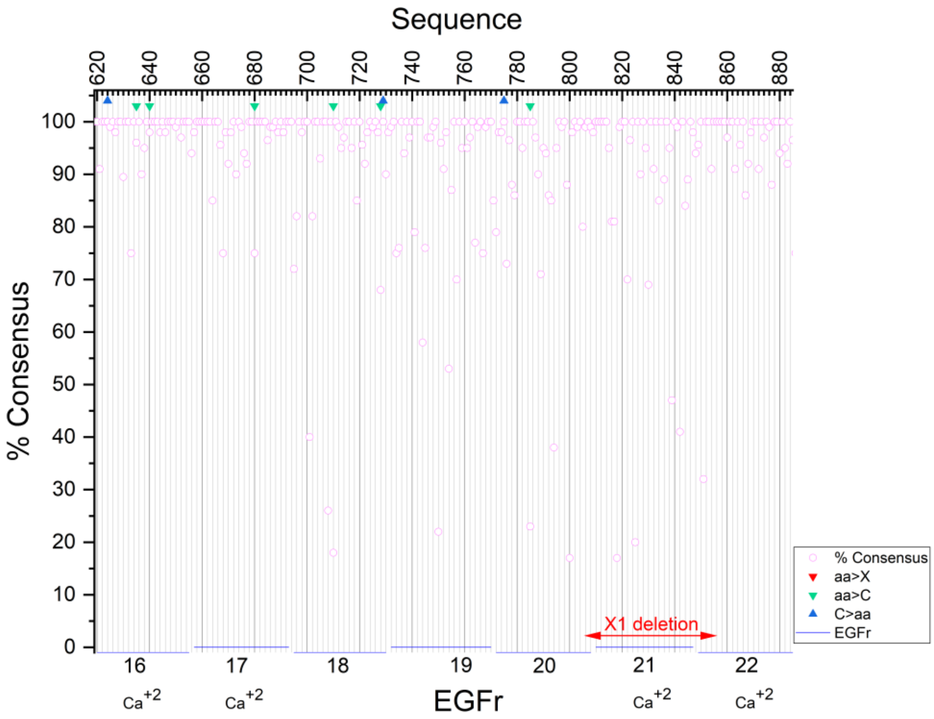
Sequence
620
640
660
680
700
720
740
760
780
800
820
840
860
880
% Consensus
100
90
80
70
60
50
40
30
20
10
0
X1 deletion
16
17
18
19
20
21
22
Ca+2
EGFr
% Consensus
aa>X
aa>C
C>aa
EGFr

d

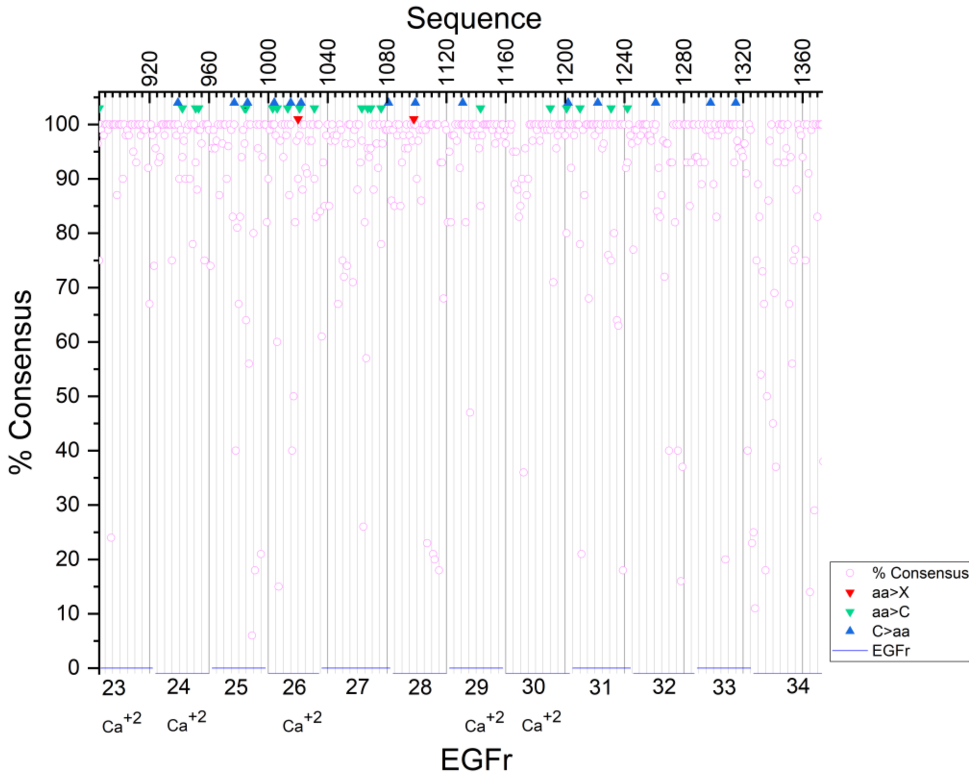
Sequence
920
960
1000
1040
1080
1120
1160
1200
1240
1280
1320
1360
% Consensus
100
90
80
70
60
50
40
30
20
10
0
Ca+2
EGFr
% Consensus
aa>X
aa>C
C>aa
EGFr

**e**

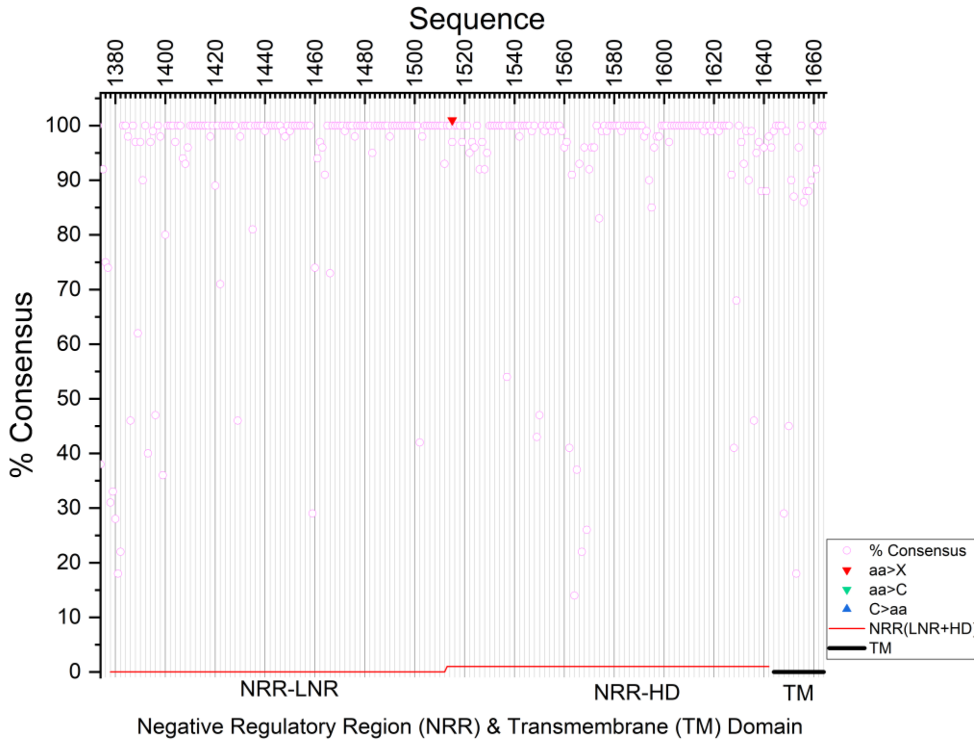
Sequence
% Consensus
% Consensus
aa>X
aa>C
C>aa
NRR(LNR+HD
TM
NRR-LNR
NRR-HD
TM
Negative Regulatory Region (NRR) & Transmembrane (TM) Domain

**f**

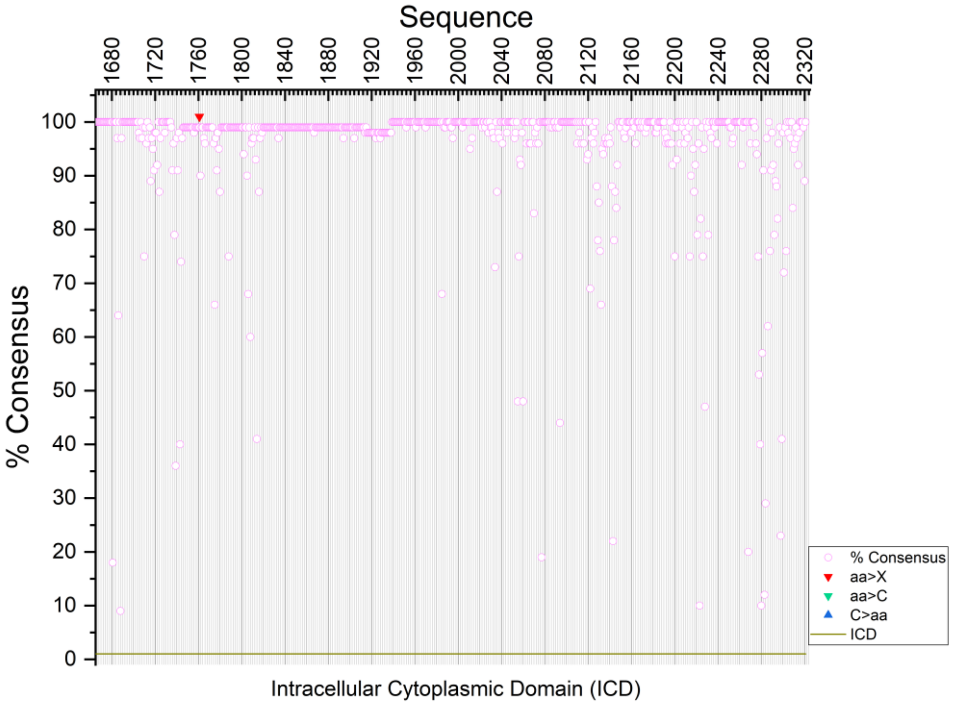
Sequence
% Consensus
% Consensus
aa>X
aa>C
C>aa
ICD
Intracellular Cytoplasmic Domain (ICD)

**Supplementary Figure 2.** The Origin plots showing the NOTCH3 sequence with various domains, pathogenic mutations and % consensus with mammalian protein. Domains are shown on X-axis as colored segments spanning six plots. 34 EGFr in ECD, blue; NRR (LNR + HD); TM, black; ICD, green; % consensus, pink circle. Various types of pathogenic mutations are shown as triangles: aa>X, red; aa>C, green; C>aa, blue. The interactive Origin plot file (**Supplementary Software 2**) features tooltips, where hovering the cursor over data points reveals the exact percent consensus, amino acid sequence, residue name, and mutation information.

Full length:

| | | | | |
|---|---|---|---|---|
| 1: Human | 100.00 | 99.09 | 91.05 | 92.83 |
| 2: Rhesus | 99.09 | 100.00 | 90.80 | 92.44 |
| 3: Mouse | 91.05 | 90.80 | 100.00 | 89.45 |
| 4: Mole-rat | 92.83 | 92.44 | 89.45 | 100.00 |

EGFr 1-34:

| | | | | |
|---|---|---|---|---|
| 1: Human | 100.00 | 98.88 | 89.23 | 92.74 |
| 2: Rhesus | 98.88 | 100.00 | 88.71 | 92.07 |
| 3: Mouse | 89.23 | 88.71 | 100.00 | 88.10 |
| 4: Mole-rat | 92.74 | 92.07 | 88.10 | 100.00 |

NRR:

| | | | | |
|---|---|---|---|---|
| 1: Human | 100.00 | 100.00 | 94.38 | 93.21 |
| 2: Rhesus | 100.00 | 100.00 | 94.36 | 93.56 |
| 3: Mouse | 94.38 | 94.36 | 100.00 | 92.45 |
| 4: Mole-rat | 93.21 | 93.56 | 92.45 | 100.00 |

ICD:

| | | | | |
|---|---|---|---|---|
| 1: Human | 100.00 | 99.09 | 94.17 | 93.77 |
| 2: Rhesus | 99.09 | 100.00 | 94.17 | 93.77 |
| 3: Mouse | 94.17 | 94.17 | 100.00 | 92.20 |
| 4: Mole-rat | 93.77 | 93.77 | 92.20 | 100.00 |

```
                                                                 EGFr1
            Exon1                                          Exon2(A)
Human       MGPGARGRRRRRRPMSPPPPPP-PVRA---LPLLLLLAGPGAAAPPCLDGSPCANGGRCT     56
Rhesus      MGPGARGRRRRRRPMSPPPPPPP-VR---ALPLLLLLAGPGAAVPPCLDGSPCANGGRCT     56
Mouse       MGLGARGRRRRRRLMALPPPPPPM-R---ALPLLLLLAGLGAAAPPCLDGSPCANGGRCT     56
Mole-rat    MGPGARGRRGRS--MSPPPPPPPPKGTLPQLLLLPLLAGLGAAAPACLDGSPCANGGRCT     58
            ** ****** *   *: *****        * ** **** ***.* **************

                Exon3(L)             EGFr2                       Exon4(G)
Human       Q-LPSREAACLCPPGWVGERCQLEDPCHSGPCAGRGVCQSSVVAGTARFSCRCPRGFRGP     115
Rhesus      Q-LPSREAACLCPPGWVGERCQLEDPCHSGPCAGRGVCQSSVVAGTARFSCRCPRGFRGP     115
Mouse       HQQPSLEAACLCLPGWVGERCQLEDPCHSGPCAGRGVCQSSVVAGTARFSCRCLRGFQGP     116
Mole-rat    Q-LPSQEAACLCPPGWVGERCQLEDPCHSGPCAGRGVCQSSVVAGTARFSCRCPRGFRGP     117
            :  ** ****** **************************************** ***:**

                                    ↑
```

```
            EGFr3                                     Ca binding-EGFr4
Human       DCSLPDPCLSSPCAHGARCSVGPDGRFLCSCPPGYQGRSCRSDVDECRVGEPCRHGGTCL   175
Rhesus      DCSLPDPCLSSPCAHSARCSVGPDGRFLCSCPPGYQGRSCRSDVDECRVGEPCRHGGTCL   175
Mouse       DCSQPDPCVSRPCVHGAPCSVGPDGRFACACPPGYQGQSCQSDIDECRSGTTCRHGGTCL   176
Mole-rat    DCSLPDPCLSSPCAHTARCSVGSDGRFVCSCPPGYQGRSCQSDMDECRVAGLCRHGGTCL   177
            *** ****:* **.* * **** **** *:*******:**:**:**** .  ********
                ↑

                                EGFr5                 Exon5(G)
Human       NTPGSFRCQCPAGYTGPLCENPAVPCAPSPCRNGGTCRQSGDLTYDCACLPGFEGQNCEV   235
Rhesus      NTPGSFRCQCPAGYTGPLCENPAVPCAPSPCRNGGTCRQSGDLTYDCACLPGFEGQNCEV   235
Mouse       NTPGSFRCQCPLGYTGLLCENPVVPCAPSPCRNGGTCRQSSDVTYDCACLPGFEGQNCEV   236
Mole-rat    NMPGSFRCQCPGGYTGPLCEDPTVPCAPSPCRNGGTCRQSGDLTYDCACLPGFEGQNCEV   237
            * ********* ********:*.****************.*:*****************
                               ↑

                                                     Ca-EGFr7
            Ca-EGFr6                    Exon6(G)
Human       NVDDCPGHRCLNGGTCVDGVNTYNCQCPPEWTGQFCTEDVDECQLQPNACHNGGTCFNTL   295
Rhesus      NVDDCPGHRCLNGGTCVDGVNTYNCQCPPEWTGQFCTEDVDECQLQPNACHNGGTCFNTL   295
Mouse       NVDDCPGHRCLNGGTCVDGVNTYNCQCPPEWTGQFCTEDVDECQLQPNACHNGGTCFNLL   296
Mole-rat    NVDDCPGHRCLNGGTCVDGVNTYNCQCPPEWTGQFCTEDVDECQLQPNACHNGGTCFNTL   297
            ********************************************************* *

                                                                   EGFr9
                             Ca-EGFr8                  Exon7(G)
Human       GGHSCVCVNGWTGESCSQNIDDCATAVCFHGATCHDRVASFYCACPMGKTGLLCHLDDAC   355
Rhesus      GGHSCVCVNGWTGESCSQNIDDCATAVCFHGATCHDRVASFYCACPMGKTGLLCHLDDAC   355
Mouse       GGHSCVCVNGWTGESCSQNIDDCATAVCFHGATCHDRVASFYCACPMGKTGLLCHLDDAC   356
Mole-rat    GGHSCVCVNGWTGESCSQNIDDCATAVCFHGATCHDRVASFYCACPMGKTGLLCHLDDAC   357
            ************************************************************

                                                  Ca-EGFr10
                                                   Exon8(G)
Human       VSNPCHEDAICDTNPVNGRAICTCPPGFTGGACDQDVDECSIGANPCEHLGRCVNTQGSF   415
Rhesus      VSNPCHEDAICDTNPVNGRAICTCPPGFTGGACDQDVDECSIGANPCEHLGRCVNTQGSF   415
Mouse       VSNPCHEDAICDTNPVSGRAICTCPPGFTGGACDQDVDECSIGANPCEHLGRCVNTQGSF   416
Mole-rat    VSNPCHEDAVCDTNPVNGRAICTCPPGFTGGACDQDVDECSIGANPCEHLGRCVNTQGSF   417
            *********:******.*******************************************

                          Ca-EGFr11                Exon9(G)      Ca-EGFr12
Human       LCQCGRGYTGPRCETDVNECLSGPCRNQATCLDRIGQFTCICMAGFTGTYCEVDIDECQS   475
Rhesus      LCQCGRGYTGPRCETDVNECLSGPCRNQATCLDRIGQFTCICMAGFTGTYCEVDIDECQS   475
Mouse       LCQCGRGYTGPRCETDVNECLSGPCRNQATCLDRIGQFTCICMAGFTGTYCEVDIDECQS   476
Mole-rat    LCQCGRGYTGPRCETDVNECLSGPCRNQATCLDRIGQFTCICMAGFTGTYCEVDIDECQS   477
            ************************************************************

                          Exon10(G)           Ca-EGFr13
Human       SPCVNGGVCKDRVNGFSCTCPSGFSGSTCQLDVDECASTPCRNGAKCVDQPDGYECRCAE   535
Rhesus      SPCVNGGICKDRVNGFSCTCPSGFSGSTCQLDVDECASTPCRNGAKCVDQPDGYECRCAE   535
Mouse       SPCVNGGVCKDRVNGFSCTCPSGFSGSMCQLDVDECASTPCRNGAKCVDQPDGYECRCAE   536
Mole-rat    SPCVNGGVCKDRVNGFSCTCPSGFSGTTCQLDVDECASTPCKNGAKCVDQPDGYECRCAE   537
            *******:******************: *************:******************

             Exon11(G)  Ca-EGFr14                            Ca-EGFr15
Human        GFEGTLCDRNVDDCSPDPCHHGRCVDGIASFSCACAPGYTGTRCESQVDECRSQPCRHGG   595
Rhesus       GFEGMLCERNVDDCSPDPCHHGRCVDGIASFSCACAPGYTGTRCESQVDECRSQPCRHGG   595
Mouse        GFEGTLCERNVDDCSPDPCHHGRCVDGIASFSCACAPGYTGIRCESQVDECRSQPCRYGG   596
Mole-rat     GFEGTLCERNVDDCSPDPCHHGRCVDGIASFTCACVPGYTGTRCESQVDECRSQPCRHGG   597
             **** **:**********************:***.***** ***************:**
                              ↑
```

```
                                   Ca-EGFr16
                    Exon12(G)                            Exon13(G)
Human     KCLDLVDKYLCRCPSGTTGVNCEVNIDDCASNPCTFGVCRDGINRYDCVCQPGFTGPLCN   655
Rhesus    KCLDLVDKYLCRCPSGTTGVNCEVNIDDCASNPCSFGVCRDGINRYDCVCQPGFTGPLCN   655
Mouse     KCLDLVDKYLCRCPPGTTGVNCEVNIDDCASNPCTFGVCRDGINRYDCVCQPGFTGPLCN   656
Mole-rat  KCLDLVDKYLCRCPPGTTGVNCEVNIDDCASNPCTFGVCRDGINRYDCVCQPGFTGPLCN   657
          ************** ******************* :*************************

           Ca-EGFr17                            EGFr18         Exon14(G)
Human     VEINECASSPCGEGGSCVDGENGFRCLCPPGSLPPLCLPPSHPCAHEPCSHGICYDAPGG   715
Rhesus    VEINECASSPCGEGGSCVDGENGFRCLCPPGSLPPLCLPPSHPCAHDPCSHGICYDAPGG   715
Mouse     VEINECASSPCGEGGSCVDGENGFHCLCPPGSLPPLCLPANHPCAHKPCSHGVCHDAPGG   716
Mole-rat  VEINECASSPCGEGGSCLDVENGFRCLCPPGSLPPLCLPPSHACAHEPCSHGVCHDAPGG   717
          *****************:* ****:************** .* ***.*****:*:*****
                                                       ↑
                                                                EGFr20
                      EGFr19                   Exon15(G)
Human     FRCVCEPGWSGPRCSQSLARDACESQPCRAGGTCSSDGMGFHCTCPPGVQGRQCELLSPC   775
Rhesus    FRCVCEPGWSGPRCSQSLARDACESQPCRAGGTCSSDGIGFHCTCPPGVQGRQCELLSPC   775
Mouse     FRCVCEPGWSGPRCSQSLAPDACESQPCQAGGTCTSDGIGFRCTCAPGFQGHQCEVLSPC   776
Mole-rat  FRCVCEPGWSGPRCSQSLARDACESQPCRGGGTCVSDRMSFHCTCPPGVQGRQCEVLSPC   777
          ******************* ********:.**** ** :.*:*** **.**:***:****

                                             Ca-EGFr21
                              Exon16(G)(deleted in human X1 isoform)
Human     TPNPCEHGGRCESAPGQLPVCSCPQGWQGPRCQQDVDECAGPAPCGPHGICTNLAGSFSC   835
Rhesus    TPNPCEHGGRCESAPGQLPVCSCPQGWQGPRCQQDVDECAGPAPCGPHGICTNLAGSFSC   835
Mouse     TPSLCEHGGHCESDPDRLTVCSCPPGWQGPRCQQDVDECAGASPCGPHGTCTNLPGNFRC   836
Mole-rat  TPNPCEHGGRCESAPGQPVVCSCPTGWQGPRCQQDVDECASSAPCGPHGTCSNLAGSFSC   837
          **. *****:*** *.:  ***** ***************. :****** *:** *.* *
                    ↑

                      Ca-EGFr22
                     Exon17(N)                             Ca-EGFr23
Human     TCHGGYTGPSCDQDINDCDPNPCLNGGSCQDGVGSFSCSCLPGFAGPRCARDVDECLSNP   895
Rhesus    TCHGGYTGPSCDQDINDCDPNPCLNGGSCQDGVGSFSCSCLPGFAGPRCARDVDECLSNP   895
Mouse     ICHRGYTGPFCDQDIDDCDPNPCLHGGSCQDGVGSFSCSCLDGFAGPRCARDVDECLSSP   896
Mole-rat  TCHGGYTGPSCDQDIDDCDPNPCLNGGSCQDSVGSFSCSCLPGFAGPRCARDVDECLSSP   897
          ** ***** *****:********:******.********* ****************.*
                            ↑
                                      Ca-EGFr24
                                      Exon18(S)
Human     CGPGTCTDHVASFTCTCPPGYGGFHCEQDLPDCSPSSCFNGGTCVDGVNSFSCLCRPGYT   955
Rhesus    CGPGTCTDHVASFTCTCPPGYGGFHCEQDLPDCSPSSCFNGGTCVDGVNSFSCLCRPGYT   955
Mouse     CGPGTCTDHVASFTCACPPGYGGFHCEIDLPDCSPSSCFNGGTCVDGVSSFSCLCRPGYT   956
Mole-rat  CGPGTCTDHVASFACTCPPGYGGFHCEKDLPDCSPSSCFNGGTCVDRVNSFSCLCRPGYT   957
          *************:*:*********** ****************** *.***********

                                                    EGFr26
              EGFr25                       Exon19(T)
Human     GAHCQHEADPCLSRPCLHGGVCSAAHPGFRCTCLESFTGPQCQTLVDWCSRQPCQNGGRC   1015
Rhesus    GAHCQHEADPCLSRPCLHGGVCSAAHPGFRCTCPQSFTGPQCQTLVDWCSRQPCQNGGRC   1015
Mouse     GTHCQYEADPCFSRPCLHGGICNPTHPGFECTCREGFTGSQCQNPVDWCSQAPCQNGGRC   1016
Mole-rat  GAHCQYEADPCLSRPCLNGGVCSTTHPGFHCACLEGFAGSQCQTLVDWCSQAPCQNGGHC   1017
          *:***:*****:*****:**:*. :****.*:* :.*:* ***. *****: ******:*
               ↑
                              EGFr27Exon20(G)
Human     VQTGAYCLCPPGWSGRLCDIRSLPCREAAAQIGVRLEQLCQAGGQCVDEDSSHYCVCPEG   1075
Rhesus    VQTGAYCLCPPGWSGRLCDIRSLPCREAAAQIGVRLEQLCQAGGQCVDEDSSHYCVCPEG   1075
Mouse     VQTGAYCICPPGWSGRLCDIQSLPCTEAAAQMGVRLEQLCQEGGKCIDKGRSHYCVCPEG   1076
Mole-rat  VQTGAYCLCRPGWSGRLCDIRSLPCKEAAAQIGVRLEHLCQAGGQCVDKGSSHSCVCPEG   1077
          *******:* **********:**** *****:*****:*** **:*:*:. ** ******
```

```
                    EGFr28          Exon21(C)           Ca-EGFr29
Human      RTGSHCEQEVDPCLAQPCQHGGTCRGYMGGYMCECLPGYNGDNCEDDVDECASQPCQHGG   1135
Rhesus     RTGSHCEQEVDPCLAQPCQHGGTCRGYMGGYMCECLPGYNGENCEDDVDECASQPCQHGG   1135
Mouse      RTGSHCEHEVDPCTAQPCQHGGTCRGYMGGYVCECPAGYAGDSCEDNIDECASQPCQNGG   1136
Mole-rat   RTGSHCEQEVDPCLAQPCQHGGTCRGYMGGYMCECPAGYSGDNCEDDVDECASQPCQHGG   1137
           *******:***** ****************:***  ** *:.***::*********:**
                        ↑
                                  Ca-EGFr30
                        Exon22(G)
Human      SCIDLVARYLCSCPPGTLGVLCEINEDDCGPGPPLDSGPRCLHNGTCVDLVGGFRCTCPP   1195
Rhesus     SCIDLVARYLCSCPPRTLGVLCEINEDDCGPGPPLDSGPRCLHNGTCVDLVGGFRCTCPP   1195
Mouse      SCIDLVARYLCSCPPGTLGVLCEINEDDCDLGPSLDSGVQCLHNGTCVDLVGGFRCNCPP   1196
Mole-rat   SCIDLVARYLCSCPPGTLGVLCEINEDDCGPGPSLDLRPRCLHNGTCVDLVGGFRCNCPP   1197
           *************** *************. ** **   :****************.***

                                                       EGFr32
                     EGFr31                  Exon23(G)
Human      GYTGLRCEADINECRSGACHAAHTRDCLQDPGGGFRCLCHAGFSGPRCQTVLSPCESQPC   1255
Rhesus     GYTGLRCEADINECRSGACHAAHTRDCLQDPGGGFRCLCHAGFSGPRCQTVLSPCESQPC   1255
Mouse      GYTGLHCEADINECRPGACHAAHTRDCLQDPGGHFRCVCHPGFTGPRCQIALSPCESQPC   1256
Mole-rat   GYTGLHCEADINECRPGACHAAHTRDCLQDPGGRFHCLCHTGFTGPRCQTALSPCESQPC   1257
           *****:********* ***************** *:*:** **:***** .*********
                                          ↑
                          Exon24(P)      EGFr33
Human      QHGGQCRPSPGPGGGLTFTCHCAQPFWGPRCERVARSCRELQCPVGVPCQQTPRGPRCAC   1315
Rhesus     QHGGQCRPSPGPGGGLTFTCHCAQPFWGPRCERVARACRELQCPVGVPCQQTPRGPRCAC   1315
Mouse      QHGGQCRHSLGRGGGLTFTCHCVPPFWGLRCERVARSCRELQCPVGIPCQQTARGPRCAC   1316
Mole-rat   HNVGQCRPSPGPGGVLTFICHCVQPFWGPHCEHVARTCRELQCPPGVPCQQTPRGPRCAC   1317
           :: **** * * ** *** ***. **** :**:***:******* *:***** *******

                              EGFr34
Human      PPGLSGPSCRSFPGSPPGASNASCAAAPCLHGGSCRPAPLAPFFRCACAQGWTGPRCEAP   1375
Rhesus     PPGLSGPSCRSFSGSPPGASNASCAAAPCLHGGSCRPAPLAPFFRCTCAQGWTGPRCEAP   1375
Mouse      PPGLSGPSCRVSRASPSGATNASCASAPCLHGGSCLPVQSVPFFRCVCAPGWGGPRCETP   1376
Mole-rat   PPGLSGLSCRSR-AVPPGGTNASCAAAPCLHGGSCRPAPLAPFFRCGCAPGWAGLRCETP   1376
           ****** ***   . * *.:*****:********* *.  .***** ** ** * ***:*

              NRR-LNR
Human      A--AAPEVSEEPRCPRAACQAKRGDQRCDRECNSPGCGWDGGDCSLSVGDPWRQCEALQC   1433
Rhesus     A--AAPEVSEEPRCPRAACQAKRGDQRCDRECNSPGCGWDGGDCSLSVGDPWRQCEALQC   1433
Mouse      S--AAPEVPEEPRCPRAACQAKRGDQNCDRECNTPGCGWDGGDCSLNVDDPWRQCEALQC   1434
Mole-rat   AAAAAPEVAEEPQCPRAACQAKRGDRRCDRQCNTRGCGWDGGDCSLNVDDPWRQCEALQC   1436
           :  ***** ***:*************:.***:**: ***********.*.**********

           NRR-LNR                    Exon25(N)
Human      WRLFNNSRCDPACSSPACLYDNFDCHAGGRERTCNPVYEKYCADHFADGRCDQGCNTEEC   1493
Rhesus     WRLFNNSRCDPACSSPACLYDNFDCHAGGRERTCNPVYEKYCADHFADGRCDQGCNTEEC   1493
Mouse      WRLFNNSRCDPACSSPACLYDNFDCYSGGRDRTCNPVYEKYCADHFADGRCDQGCNTEEC   1494
Mole-rat   WRLFNNSRCDLACSSPACLYDNFDCH--GRERTCNPVYEKYCADHFADGRCDQGCNTEEC   1494
           ********** **************:  **:*****************************

              NRR-LNR          NRR-HD
Human      GWDGLDCASEVPALLARGVLVLTVLLPPEELLRSSADFLQRLSAILRTSLRFRLDAHGQA   1553
Rhesus     GWDGLDCASEVPALLARGVLVLTVLLPPEELLRSSADFLQRLSAILRTSLRFRLDAHGQA   1553
Mouse      GWDGLDCASEVPALLARGVLVLTVLLPPEELLRSSADFLQRLSAILRTSLRFRLDARGQA   1554
Mole-rat   GWDGLDCASEVPALLARGVLVLTVLMPPEELLRSSADFLQRLSAILRTSLRFRLDSRGQA   1554
           *************************:*****************************::***

                              Exon26(G)
Human      MVFPYHRPSPGSEPRARR↓ELAPEVIGSVVMLEIDNRLCLQSPENDHCFPDAQSAADYLGA   1613
Rhesus     MVFPYHRPSPGSEPRARR↓ELAPEVIGSVVMLEIDNRLCLQSPENDHCFPDAQSAADYLGA   1613
Mouse      MVFPYHRPSPGSESRVRR↓ELGPEVIGSVVMLEIDNRLCLQSAENDHCFPDAQSAADYLGA   1614
Mole-rat   MVFPYHRPSSASESRTRR↓ELAPEVIGSVVMLEIDNRLCLQSPENDHCFPDAQSAADYLGA   1614
           ********* .** *.** **.******************** ******************
                 1571-1572 (RE) S1 Cleavage site by Furin-like protease
```

```
                    NRR-HD      Exon27(G)             TRANS MEMBRANE REGION
Human        LSAVERLDFPYPLRD↓VRGEPLEPPEPSVPLLPLLVAGAVLLLVILVLG↓VMVARRKREHST  1673
Rhesus       LSAVERLDFPYPLRD↓VRGEPLEPPEPSVPLLPLLVAGAVLLLVILVLG↓VMVARRKREHST  1673
Mouse        LSAVERLDFPYPLRD↓VRGEPLEAPEQSVPLLPLLVAGAVFLLIIFILG↓VMVARRKREHST  1674
Mole-rat     LSAVERLDFPYPLRD↓VRGEPLEPPEPTVPVLPLLAAGAVFLLVILVLG↓VMVARRKREHST  1674
             *************** ******* ** :**:****.****:**:*::** ************
             1628-1629 (DV) S2 cleavage site by ADAM-10 1661-1662 (GV)
             S3 Cleavage site by gamma-secretase complex

                                       Exon28(K)
Human        LWFPEGFSLHKDVASGHKGRREPVGQDALGMKNMAKGESLMGEVATDWMDTECPEAKRLK  1733
Rhesus       LWFPEGFSLHKDVAAGHKGRREPVGQDALGMKNMAKGESLMGEVATDWMDTECPEAKRLK  1733
Mouse        LWFPEGFALHKDIAAGHKGRREPVGQDALGMKNMAKGESLMGEVVTDLNDSECPEAKRLK  1734
Mole-rat     LWFPEGFSLHKDVAAGHKGRREPVGQDALGMKSMAKGESLMGEVATDWMDTDCPEAKRLK  1734
             *******:****:*:*****************.***********.**  *::********

             Exon29(V)                                      Exon30(D)
Human        VEEPGMGAEEAVDCRQWTQHHLVAADIRVAPAMALTPPQGDADADGMDVNVRGPDGFTPL  1793
Rhesus       VEELGMGAEEAVDCRQWTQHHLVAADIRVAPAMALTPPQGDADADGMDVNVRGPDGFTPL  1793
Mouse        VEEPGMGAEEPEDCRQWTQHHLVAADIRVAPATALTPPQGDADADGVDVNVRGPDGFTPL  1794
Mole-rat     MEEPGVGAKEPVDCRQWTQHHLVAADIRVAPAMALTPPQGDADADGMDVNVRGPDGFTPL  1794
             :** *:**:*  ******************* *************:*************

Human        MLASFCGGALEPMPTEEDEADDTSASIISDLICQGAQLGARTDRTGETALHLAARYARAD  1853
Rhesus       MLASFCGGALEPMPTEEDEADDTSASIISDLICQGAQLGARTDRTGETALHLAARYARAD  1853
Mouse        MLASFCGGALEPMPAEEDEADDTSASIISDLICQGAQLGARTDRTGETALHLAARYARAD  1854
Mole-rat     MLASFCGGALEPVPAEEEEVEDTSASIISDLICQGAQLGARTDRTGETALHLAARYARAD  1854
             ************:*:**:*.:***************************************

                                                Exon31(I)
Human        AAKRLLDAGADTNAQDHSGRTPLHTAVTADAQGVFQILIRNRSTDLDARMADGSTALILA  1913
Rhesus       AAKRLLDAGADTNAQDHSGRTPLHTAVTADAQGVFQILIRNRSTDLDARMADGSTALILA  1913
Mouse        AAKRLLDAGADTNAQDHSGRTPLHTAVTADAQGVFQILIRNRSTDLDARMADGSTALILA  1914
Mole-rat     AAKRLLDAGADTNAQDHSGRTPLHTAVTADAQGVFQILIRNRSTDLDARMADGSTALILA  1914
             ************************************ ***********************

                                  Exon32(G)                      Exon33(E)
Human        ARLAVEGMVEELIASHADVNAVDELGKSALHWAAAVNNVEATLALLKNGANKDMQDSKEE  1973
Rhesus       ARLAVEGMVEELIASHADVNAVDELGKSALHWAAAVNNVEATLALLKNGANKDMQDSKEE  1973
Mouse        ARLAVEGMVEELIASHADVNAVDELGKSALHWAAAVNNVEATLALLKNGANKDMQDSKEE  1974
Mole-rat     ARLAVEGMVEELIASHADVNAVDELGKSALHWAAAVNNVEATLALLKNGANKDMQDSKEE  1974
             ************************************************************

Human        TPLFLAAREGSYEAAKLLLDHFANREITDHLDRLPRDVAQERLHQDIVRLLDQPSGPRSP  2033
Rhesus       TPLFLAAREGSYEAAKLLLDHFANREITDHLDRLPRDVAQERLHQDIVRLLDQPSGPRSP  2033
Mouse        TPLFLAAREGSYEAAKLLLDHLANREITDHLDRLPRDVAQERLHQDIVRLLDQPSGPRSP  2034
Mole-rat     TPLFLAAREGSYEAAKLLLDHFANREITDHLDRLPRDVAQERLHQDIVRLLDQPSGPRSP  2034
             *********************:**************************************

Human        PGPHGLGPLLCPPGAFLPGLKAAQSGSKKSRRPPGKAGLGPQGPRGRGKKLTLACPGPLA  2093
Rhesus       PGTHGLGPLLCPPGAFLPGLKATQSGSKKSRRPPGKAGLGPQGPRGRGKKLTLACPGPLA  2093
Mouse        SGPHGLGPLLCPPGAFLPGLKAVQSGTKKSRRPPGKTGLGPQGTRGRGKKLTLACPGPLA  2094
Mole-rat     PGPHGLGPLLCPPGAFLPGLKTAPSGTKKSRRPPGKAGLGPQGTRGRGKKLTLACPGPLA  2094
              * *******************:. **:*********:****** ****************

Human        DSSVTLSPVDSLDSPRPFGGPPASPGGFPLEGPYAAATATAVSLAQLGGPGRAG-LGRQP  2152
Rhesus       DSSVTLSPVDSLDSPRPFGGPPASPGGFPLEGPYAAATATAVSLAQLGGPGRAG-LGRQP  2152
Mouse        DSSVTLSPVDSLDSPRPFSGPPASPGGFPLEGPYAT-TATAVSLAQLG-ASRAGPLGRQP  2152
Mole-rat     ESSVTLSPVDSLDSPRAFGGPPASPGSFPIEGPYAAATATAVSLAQLGGPGRAGPLGRQP  2154
             :*************** *.*******.**:*****: *********** .*** *****
```

```
Human         PGGCVLSLGLLNPVAVPLDWARLPPPAPPGPSFLLPLAPGPQLLNPGTPVSPQERPPPYL     2212
Rhesus        PGGCVLSLGLLNPVAVPLDWARLPPPAPPGPSFLLPLAPGPQLLNPGTPVSPQERPPPYL     2212
Mouse         PGGCVLSFGLLNPVAVPLDWARLPPPAPPGPSFLLPLAPGPQLLNPGAPVSPQERPPPYL     2212
Mole-rat      PGGCVLSLGLLNPVAVPLDWARLPPPAPPGPSFLLPLAPGPQLLNPGAPVSPQERPPPYL     2214
              *******:***************************************:************

Human         AVPGHGEEYPVAGAHSSPPKARFLRVPSEHPYLTPSPESPEHWASPSPPSLSDWSESTPS     2272
Rhesus        AVPGHGEEYPAAGAHSSPPKARFLRVPSEHPYLTPSPESPEHWASPSPPSLSDWSESTPS     2272
Mouse         AAPGHGEEYPAAGTRSSPTKARFLRVPSEHPYLTPSPESPEHWASPSPPSLSDWSDSTPS     2272
Mole-rat      AAPGHGEEYPASGTQSSPPKPRFLRVPSEHPYLTPSPESPEHWASPSPPSLSDWSDSTPS     2274
              *.********.:*::*** * **********************************:****

Human         PATATGAMATT-TGALPAQPLPLSVPSSLAQAQTQLGPQPEVTPKRQVLA 2321
Rhesus        PATATGAMATA-TGALPAQPLPLSVPSSLAQAQTQLGPQPEVTPKRQVLA 2321
Mouse         PATATNAT---ASGALPAQPHPISVPS-LPQSQTQLGPQPEVTPKRQVMA 2318
Mole-rat      PAIAAGATATAAAAGLPSQPHPLSIPGSLGQAQTQLGPQPEATPKRQVLA 2324
              ** *:.*     :..**:** *:*:*. * *:*********.******:*
```

**Key:**
**Highlighted residues:**
Yellow, Cysteine
Green, Putative $Ca^{+2}$-binding (predicted by CD/NCBI)

**Colored/boxed residues (Human):**
**Red/bold, Pathogenic (confirmed by clinical data from UniProt, NextProt and the references)**
**Bold**/underlined: Deletion
Horizontal box: EGFr domains
Blue residue: Exon starting aa.
Dashed box: NRR-LNR region
Solid box: NRR-HD region
Arrows: DP bonds within EGF domains

**-Cleaved region for signal peptide in NOTCH3 (aa):** 1-39

**-NOTCH3 receptor (ectosomal) domain (aa):** 40-1643
34 EGFr containing region from 25 exons (aa): 40-1373 (18 Ca-binding EGFr domain. Many of the multiple tandem repeats of the Notch ECD contain the following consensus [D/E/N]-X-[D/N]-[D/E/N/Q]-Xm-[D/N/Q]*-Xn-[F/Y] (where * indicates possible β-hydroxylation, and m and n are variables) which is predictive for calcium binding.)
There is total 33 exons in NOTCH3 gene.
RAM [RBP-Jκ (recombination-signal-sequence binding protein for Jκ genes)-associated molecule (RBPJ association module) region:
LNR region (3 cysteine-rich Lin12/Notch repeats) (aa): 1387-1505 (or 1384-1500)
NOD (NOTCH protein domain) region (aa): 1505-1561 (or 1560 NCBI)
NODP region (aa): 1576-1640 (or 1577-1637 NCBI)
Heterodimerization domain (HD):
**S1** cleavage site by furin-like protease (aa): 1571-1572 (RE residues)
**Negative regulatory region (NRR):** RAM, LNRs, HD, NOD and NODP altogether called NRR. The region maintains receptor quiescence by preventing protease cleavage prior to ligand binding and averts the activation of the signaling cascade in the absence of the ligand. (1378-1642 aa)
**S2** cleavage site by ADAM-10 (aa): 1628-1629 (DV residues)

**-Transmembrane (TM) region (aa):** 1644-1664

**S3** Cleavage site by gama-secretase complex (aa): 1661-1662 (GV residues)

**-NOTCH3 intracellular (cytoplasmic) domain (aa)**: 1665-2321
Juxtamembrane and transmembrane (JMTM) domain: 1618-1709
Ankyrin repeat (ANK) region: 1838-2000 (1788-1901 NCBI)
ANK region: 1905-2000
Oligomer interface: 1905-2002
Proline/glutamate/serine/threonine-rich (PEST) motifs:
Ligand-binding site: EGFRs 10-11

**Posttranslational modifications (PTMs) count**: There are 35 PTMs annotated in this protein:
Phosphorylation 16
Acetylation 9
Ubiquitination 4
Proteolytic cleavage 3
Hydroxylation 2: Regions 1864-1867 (DT**N**A motif) and 1931-1934 (DV**N**A motif). Green "**N**" residue.
N-linked glycosylation 1: Three residues 1179N, 1336N and 1438N (N, Asparagine).

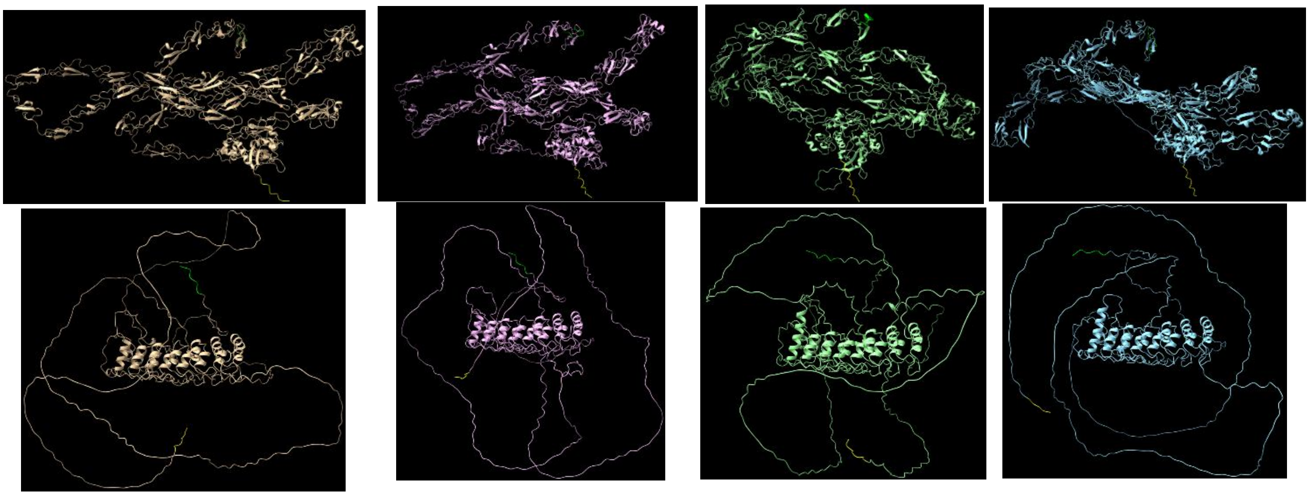

**Supplementary Figure 3.** The MSA shows sequence conservation among all EGFs, exons, and domains while AF3 modeling shows structural conservation. Upper: ECD, lower: ICD. Tan, human; pink, rhesus; blue, naked mole-rat; green, mouse. N-terminal ends, top; C-terminal ends, bottom.

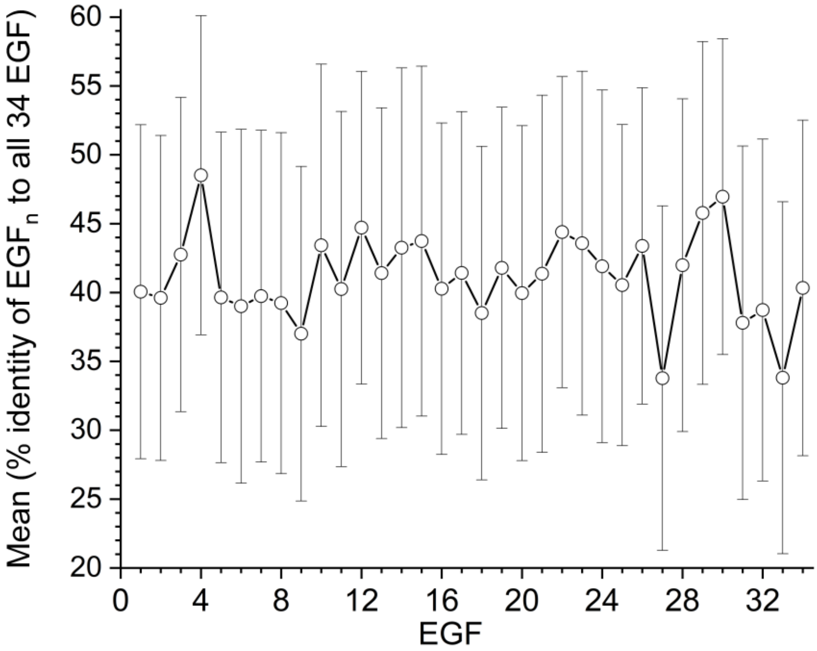

**Supplementary Figure 4.** Plot showing the mean percentage identity and standard deviation for each individual EGFr domain, calculated by comparing its sequence against all 34 EGFr domains within the human dataset.

**Supplementary Table 1.** The stability and flexibility scores of jaguar reverse aa>Cys mutations analysed by DynaMut tool.

DynaMut - Predictions Outcomes

| # | AA from | AA to | Position | Prediction ΔΔG mCSM | Prediction ΔΔG SDM | Prediction ΔΔG DUET | Prediction ΔΔG ENCoM | ΔΔS ENCoM | ΔΔG DynaMut |
|---|---|---|---|---|---|---|---|---|---|
| 1 | G | C | 81 | -1.272 kcal/mol | -2.14 kcal/mol | -1.551 kcal/mol | -2.501 kcal/mol | 3.127 kcal.mol$^{-1}$.K$^{-1}$ | **-0.688 kcal/mol** |
| 2 | R | C | 86 | -0.024 kcal/mol | -0.15 kcal/mol | -0.014 kcal/mol | -0.972 kcal/mol | 1.214 kcal.mol$^{-1}$.K$^{-1}$ | **0.012 kcal/mol** |
| 3 | G | C | 122 | -0.729 kcal/mol | -0.19 kcal/mol | -0.67 kcal/mol | 0.196 kcal/mol | -0.245 kcal.mol$^{-1}$.K$^{-1}$ | **0.33 kcal/mol** |
| 4 | P | C | 115 | -0.917 kcal/mol | -0.26 kcal/mol | -0.88 kcal/mol | -0.23 kcal/mol | 0.288 kcal.mol$^{-1}$.K$^{-1}$ | **-0.595 kcal/mol** |
| 5 | G | C | 75 | -1.054 kcal/mol | -0.27 kcal/mol | -0.9 kcal/mol | 0.053 kcal/mol | -0.066 kcal.mol$^{-1}$.K$^{-1}$ | **-0.2 kcal/mol** |
| 6 | L | C | 101 | -0.749 kcal/mol | -0.51 kcal/mol | -0.634 kcal/mol | 0.123 kcal/mol | -0.154 kcal.mol$^{-1}$.K$^{-1}$ | **-0.187 kcal/mol** |
| 7 | P | C | 110 | -0.25 kcal/mol | -0.32 kcal/mol | -0.076 kcal/mol | 0.248 kcal/mol | -0.31 kcal.mol$^{-1}$.K$^{-1}$ | **0.736 kcal/mol** |
